# Long-Read Deep Sequencing Reveals High Rates of Multilineage Transmission and Rapid Viral Population Changes in Acute HIV Infection

**DOI:** 10.1101/2025.10.15.682663

**Authors:** James I. Mullins, Wenjie Deng, Elena E. Giorgi, Craig A. Magaret, Morgane Rolland, Tanmoy Bhattacharya, Dylan H. Westfall, Anna E.J. Yssel, Roger E. Bumgarner, Ben Murrell, Thumbi Ndung’u, Merlin L. Robb, Raabya Rossenkhan, Paul T. Edlefsen, Krista L. Dong, Lennie Chen, Asanda Gwashu-Nyangiri, Hong Zhao, Ruwayhida Thebus, Fredrick Sawe, Sorachai Nitayaphan, Talita York, David Matten, Hugh Murrell, Alec P. Pankow, Michal Juraska, James Ludwig, John Hural, Myron S. Cohen, Lawrence Corey, M. Juliana McElrath, Peter B. Gilbert, Carolyn Williamson

**Author notes:** Corresponding author: Department of Microbiology, University of Washington School of Medicine, Seattle, WA, US. WD, DHW, LC: Vaccine and Infectious Disease Division, Fred Hutchinson Cancer Center, Seattle, WA, USA. AEJY: Sand Technologies, Cape Town, ZA. APP: Department of Microbiology, Icahn School of Medicine at Mount Sinai, New York, NY, US. HZ: Department of Obstetrics and Gynecology, University of Washington, Seattle, WA, US.

## Abstract

Understanding the selective forces acting upon HIV early in infection is crucial to design prevention strategies. By leveraging deep sequencing and the short diagnostic intervals of the FRESH and RV217 cohorts (median 4 days) between the last-negative and first-positive RNA tests, we captured a precise and early snapshot of acute HIV infection. The frequency of multiple transmitted viruses of 38% in these as well as placebo recipients from the AMP trials was higher than previously published, with the true frequency likely to be higher. The relative abundance of lineages fluctuated substantially over time in two-thirds of the multilineage infections, generating uncertainty in identifying the specific viruses that were transmitted and founding the infection. Viral populations exhibited diversity and selection on the Gag and Env proteins at the earliest times examined, with sites inferred to be undergoing negative selection most evident. These data may help explain vaccination failures and provide new targets for prevention.

## Introduction

Recent HIV prevention studies have shed new light on the essential components necessary for an efficacious vaccine^1-3^. However, major challenges remain, including the high and increasing genetic and antigenic diversity of HIV likely to be encountered at exposure. It is therefore crucial to understand characteristics of the acquired viruses that expand exponentially during early infection.

Despite a potentially large amount and diversity of HIV in the source (the donor), when a person is infected, a substantial genetic bottleneck occurs, with only a single or very small subset of viruses successfully establishing ongoing infection^4-9^. These variants may arise from random (stochastic) processes, higher fitness in the immunologically naïve host and initially encountered target cells, as well as their ability to survive innate immune responses^10-13^. A recent meta-analysis found an overall probability of acquiring more than one viral lineage of 25%, and the probability was higher for male-to-male transmission (30%) than for male-to-female transmission (21%)^14^.

To gain an in-depth understanding of early HIV populations we targeted sequencing of 100 molecules for each of two regions (totaling 5.5kb) of the HIV genome using a long-read, PacBio single-molecule-real-time (SMRT) platform with unique molecular identifiers (SMRT-UMI)^15^. Samples from four prospective cohorts of adult males and females were studied: RV217^16^, Females Rising through Education, Support, and Health (FRESH)^17,18^, and individuals from the placebo arms in the two Antibody Mediated Protection (AMP) trials, from the Americas (HVTN 704) and from southern Africa (HVTN 703)^1^. The RV217 and FRESH cohort participants were sampled twice weekly for viral RNA in plasma whereas the AMP trial participants were sampled monthly and then again within 1-2 weeks after the first detection of viral RNA. Viral populations were then assessed for distinct lineages, diversification and lineage fluctuations, and selection on the *gag* and *env* genes. This provided unprecedented insight into virus characteristics and lineage dynamics in early HIV infection, including prior to the onset of detectable adaptive immune-driven selection.

## Results

We generated 104,157 single-molecule, long-read viral sequences from 310 plasma samples taken from 123 prospectively identified persons living with HIV (Table 1). Two regions of the viral genome were sequenced: 2.5kb amplicons encompassing the *gag* gene and the first kb of the *pol* gene (GP fragments), and 3.0kb amplicons encompassing the *rev, vpu, env* and the first third of the *nef* gene (REN fragments, also used for construction of Env pseudotype viruses in AMP trial studies^1^) (Supplementary Fig. 1).

**Table 1.**
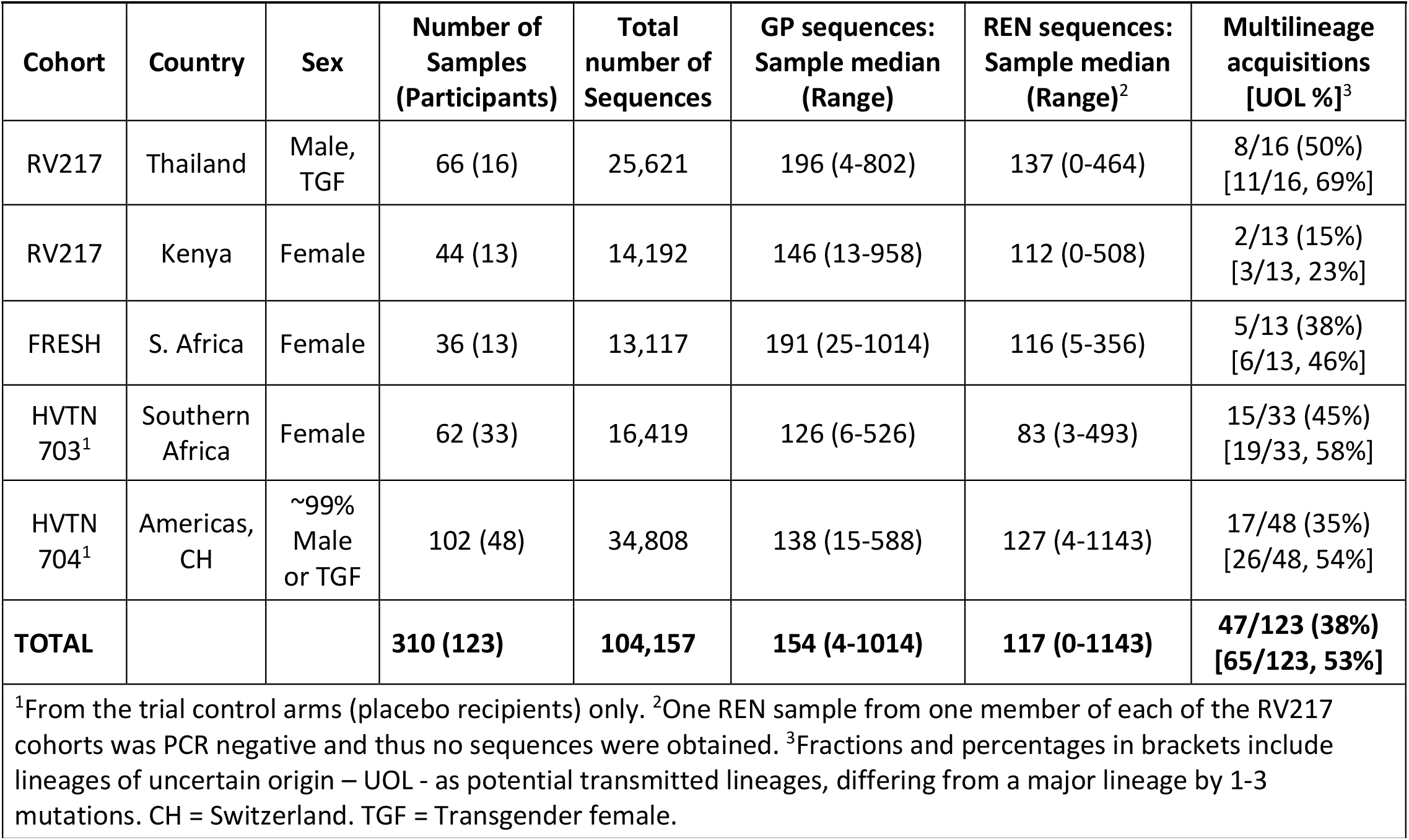
Participants, HIV sequences and multilineage acquisitions.

| **Table 1. Participants, HIV sequences and multilinesage acquisitions** |  |  |  |  |  |  |  |
| --- | --- | --- | --- | --- | --- | --- | --- |
| **Cohort** | **Country** | **Sex** | **Number of Samples (Participants)** | **Total number of Sequences** | **GP sequences: Sample median (Range)** | **REN sequences: Sample median (Range)<sup>2</sup>** | **Multilinesage acquisitions [UOL %]<sup>3</sup>** |
| RV217 | Thailand | Male, TGF | 66 (16) | 25,621 | 196 (4-802) | 137 (0-464) | 8/16 (50%)<br>[11/16, 69%] |
| RV217 | Kenya | Female | 44 (13) | 14,192 | 146 (13-958) | 112 (0-508) | 2/13 (15%)<br>[3/13, 23%] |
| FRESH | S. Africa | Female | 36 (13) | 13,117 | 191 (25-1014) | 116 (5-356) | 5/13 (38%)<br>[6/13, 46%] |
| HVTN 703<sup>1</sup> | Southern Africa | Female | 62 (33) | 16,419 | 126 (6-526) | 83 (3-493) | 15/33 (45%)<br>[19/33, 58%] |
| HVTN 704<sup>1</sup> | Americas, CH | ~99% Male or TGF | 102 (48) | 34,808 | 138 (15-588) | 127 (4-1143) | 17/48 (35%)<br>[26/48, 54%] |
| **TOTAL** |  |  | **310 (123)** | **104,157** | **154 (4-1014)** | **117 (0-1143)** | **47/123 (38%)**<br>**[65/123, 53%]** |
| <sup>1</sup>From the trial control arms (placebo recipients) only. <sup>2</sup>One REN sample from one member of each of the RV217 cohorts was PCR negative and thus no sequences were obtained. <sup>3</sup>Fractions and percentages in brackets include lineages of uncertain origin – UOL - as potential transmitted lineages, differing from a major lineage by 1-3 mutations. CH = Switzerland. TGF = Transgender female. |  |  |  |  |  |  |  |

Viral sequences were obtained from up to 5 time points (denoted by black dots along the viral load curves in Fig. 1) from the RV217 (N=29 participants) and FRESH (N=13) cohorts, over a period of 2-61 days from the estimated date of detectable HIV infection (EDDI). This date was taken to be the center of bounds (COB) between the last negative and first viral RNA positive test dates, with the COB corresponding to approximately 7 days post-acquisition^19^. The same amplicons were sequenced from 81 participants in the placebo arms from the two AMP trials^1^. These individuals were screened for HIV monthly, and plasma virus from the first HIV diagnostic timepoint was sequenced, plus 1-2 timepoints generally obtained 2-4 weeks later (Fig. 1). For these cohorts, the EDDI were derived from HIV diagnostics and viral sequence data^2,20^. Each cohort consisted of adults and included a total of 59 individuals assigned as female at birth with heterosexual transmission risk, and 64 individuals assigned as male at birth with primarily homosexual transmission risk (Table 1). All plasma samples were taken prior to participants initiating antiretroviral therapy, although 3 individuals from the HVTN 704 cohort reported taking pre-exposure prophylaxis around the time of HIV acquisition (dotted lines in Fig 1F).

**Fig 1.**
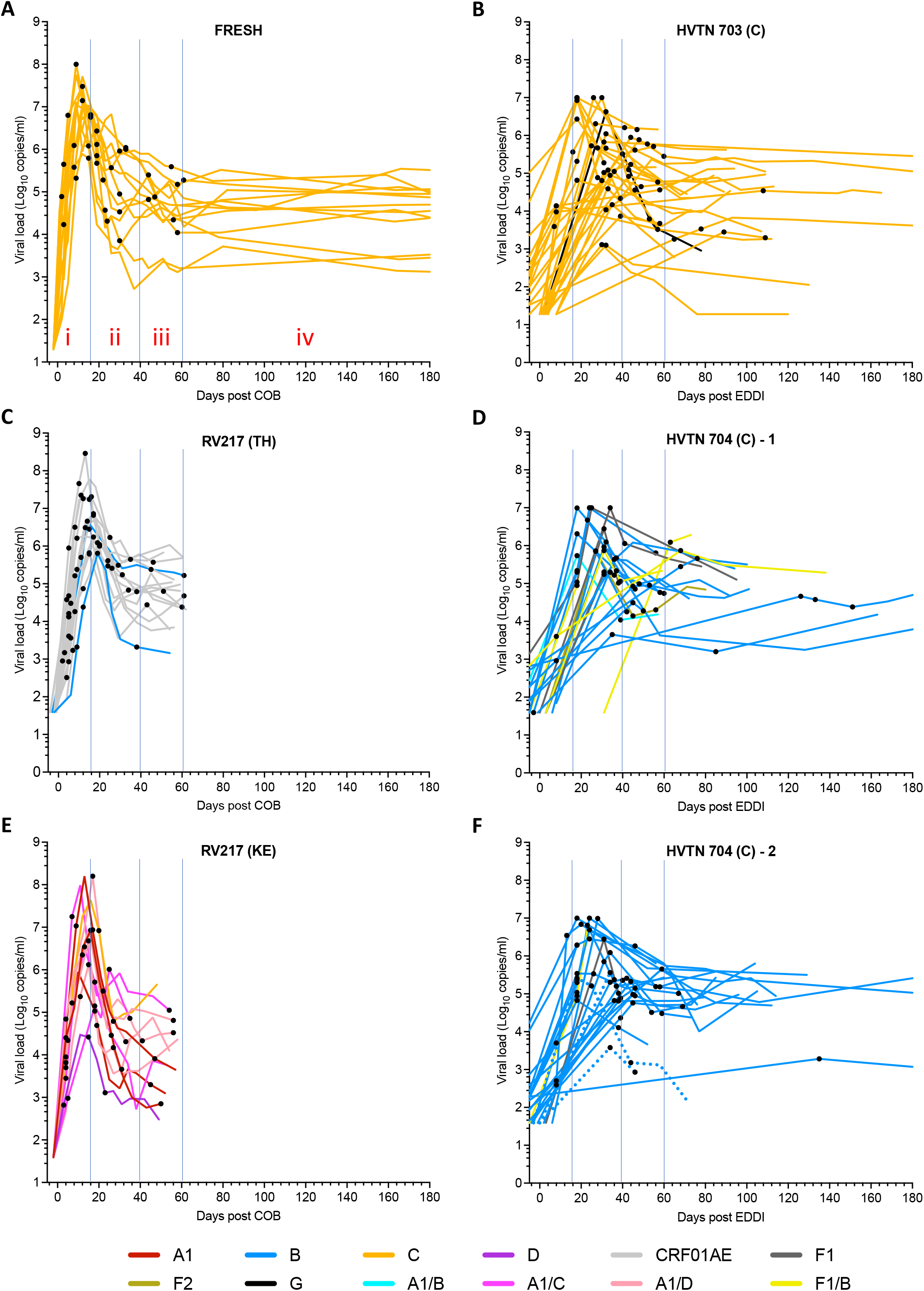
Viral load and sampling times. The name of each cohort is listed at the top of each panel. The FRESH cohort from South Africa is shown in panel **A**, and the AMP trial HVTN 703 control cohort is in panel **B**. The RV217 study included samples derived from two countries, Thailand (TH, panel **C**) and Kenya (KE, panel **E**). AMP trial participants from the HVTN 704 control cohort were the largest group studied and split over two panels (**D, F**). REN viral subtypes are indicated by colored lines, with subtypes listed in the key at the bottom of the figure. Inter-subtype recombinants are also indicated. Sample times from which HIV sequences were obtained are indicated as black dots on the curves in each panel. For the FRESH and RV217 cohorts, days post COB (Center of Bounds; corresponding to the day midway between the last HIV negative test and the first RNA positive test) is plotted along the x-axis. For the HVTN 703 and HVTN 704 cohorts, which were sampled less frequently, the estimated dates of detectable infection (EDDI) were derived from HIV diagnostics and viral sequence data as described^2,20^. For some analyses, sequence datasets were divided into four time frames (i-iv) see panel A), as demarcated by thin blue vertical lines (see S2 Table).

Viral subtype analyses of the REN region revealed 45 acquisitions of subtype C, 38 with subtype B, 15 with CRF01AE, as well as smaller numbers of individuals acquiring subtypes A1, F, G, D (N=10) or inter-subtype recombinants (N=15) (Supplementary Table 1, Supplementary Fig. 2). In 12 participants, subtype discordance was found between the two genomic regions sequenced, 11 of which involved intersubtype recombinants. This was most evident in the RV217 cohort from Kenya (in 5 of 13 participants), reflecting the long-standing presence of multiple subtypes in this region of Africa. Similarly, cocirculation of subtype B and F1 viruses in Peru and Brazil was reflected in the large number (N=9 of 48) of B-F1 intersubtype recombinants.

Mutational differences that were gene- and subtype-specific were noted: *gag* genes were significantly less likely than *env* genes to have putative inactivating mutations in each of two time windows (approximately 30 and 45 days EDDI, respectively; Supplementary Tables 2-4, Supplementary Fig. 3A-B; See Online Methods). Subtype C viruses had lower rates of intact genes in both Gag and Env than the subtype B and in Gag when compared to the combined group “other” (composed of other subtypes and recombinants) sequence datasets (Supplementary Table 4A-B, Supplementary Fig. 3C-D). Surprisingly, despite a lower rate of intact genes, subtype C genes were not more likely to be hypermutated (Supplementary Fig. 3E-F; Supplementary Discussion 1), which frequently results in the formation of stop codons. No correlation was found between viral loads and intact gene proportions in either gene (Supplementary Discussion 2). However, when sample datasets with no hypermutated sequences were excluded, there was a strong trend for hypermutated sequences to decline with increasing viral loads: p=0.0006 and 0.06 for *gag* at the first and second time windows, respectively, and p=0.005 and 0.02 for *env*. (Supplementary Fig. 4).

Two approaches were taken to discern whether an infection resulted from single or multiple founding lineages and to count lineages: Poisson Fitter^21^ (Supplementary Discussion 4) was used to examine sequences from the first available time point, and; an iterative method was devised that included clustering of sequences from all time points into distinct phylogenetic clades, followed by a manual review of pairwise distances to refine lineage designations (See Online methods and Supplementary Fig. 5). After removal of recombinants between lineages, pairwise distance distributions typically clustered the sequences unambiguously (Supplementary Fig. 6). The results from the earliest sample from all participants was 100% concordant between the Poisson Fitter and phylogenetic/distance methods. We report results for the phylogenetic/distance method, given that it included sequences from all time points.

The frequency of multilineage acquisitions (MLA) was determined by considering both the GP and REN regions and all time points sampled. These considerations as well as the deep sequencing performed resulted in a higher than previously reported frequency of MLA (38%) compared to an authoritative prior meta analysis (25%)^14^. Results from GP and REN were typically concordant. However, in 6 (13%) of the 47 individuals in which multiple lineages were identified, only one region (4 in GP only and 2 in REN only) had evidence of an MLA, and in 3 of 47 (6%) a second lineage was only identified at the second time point. Delayed outgrowth of some lineages (noted in 4 cases in GP and 7 cases in REN), may contribute to this discordance and to the high MLA frequency. However, a lack of distinction between lineages in one region, or low viral loads and the consequent failure to achieve deep sequencing at the first time point was likely to be the cause of this discrepancy in some cases, as the median number of viral sequences recovered at the first time point in these cases was somewhat lower (Supplementary Fig. 7).

In addition to multiple lineages, 18 (24%) of the 76 individuals with otherwise single lineage acquisitions had evidence of lineages whose origin as transmitted or evolving in the new host was uncertain due to a limited number of lineage-defining differences. These uncertain-origin-lineages (UOLs) were defined if detected at the first time point and if they had 1-3 substitution mutations distinguishing them from a more common lineage (e.g., Supplementary Fig. 8). UOL sequences in these individuals represented a median of 26.5% (95% CI 15-35%) of the total number of sequences from the first time point in otherwise single lineage acquisitions. In three cases, a UOL became the dominant lineage at later time points. If indeed each UOL actually corresponded to a transmitted lineage, then 53% of the individuals we studied would have acquired multiple transmitted lineages (Table 1).

The representation of lineages or recombinants over the first two months post COB fluctuated ten-fold or more in 9 of 15 (60%) cases of multilineage acquisitions in the FRESH and RV217 cohorts (see FRESH participants 079, 267, 271 and 318 in Fig. 2, and RV217 participants 20337, 20502, 40061, 40265 and 40436 in Supplementary Fig. 9). Furthermore, initially minor variants or recombinants came to represent the major sequence population at a later time point in GP and/or REN in 6 of 15 (40%) cases (see FRESH participants 079, 271 and 318 in Fig 2, and RV217 participants 20337, 20502 and 40363 in and 318 in Fig. 2, and RV217 participants 20337, 20502, 40061, 40265 and 40436 in Supplementary Fig. 9). Overall, these major lineage shifts were found in a total of 10 of 15 (67%) multilineage acquisitions in the FRESH and RV217 cohorts.

**Fig 2.**
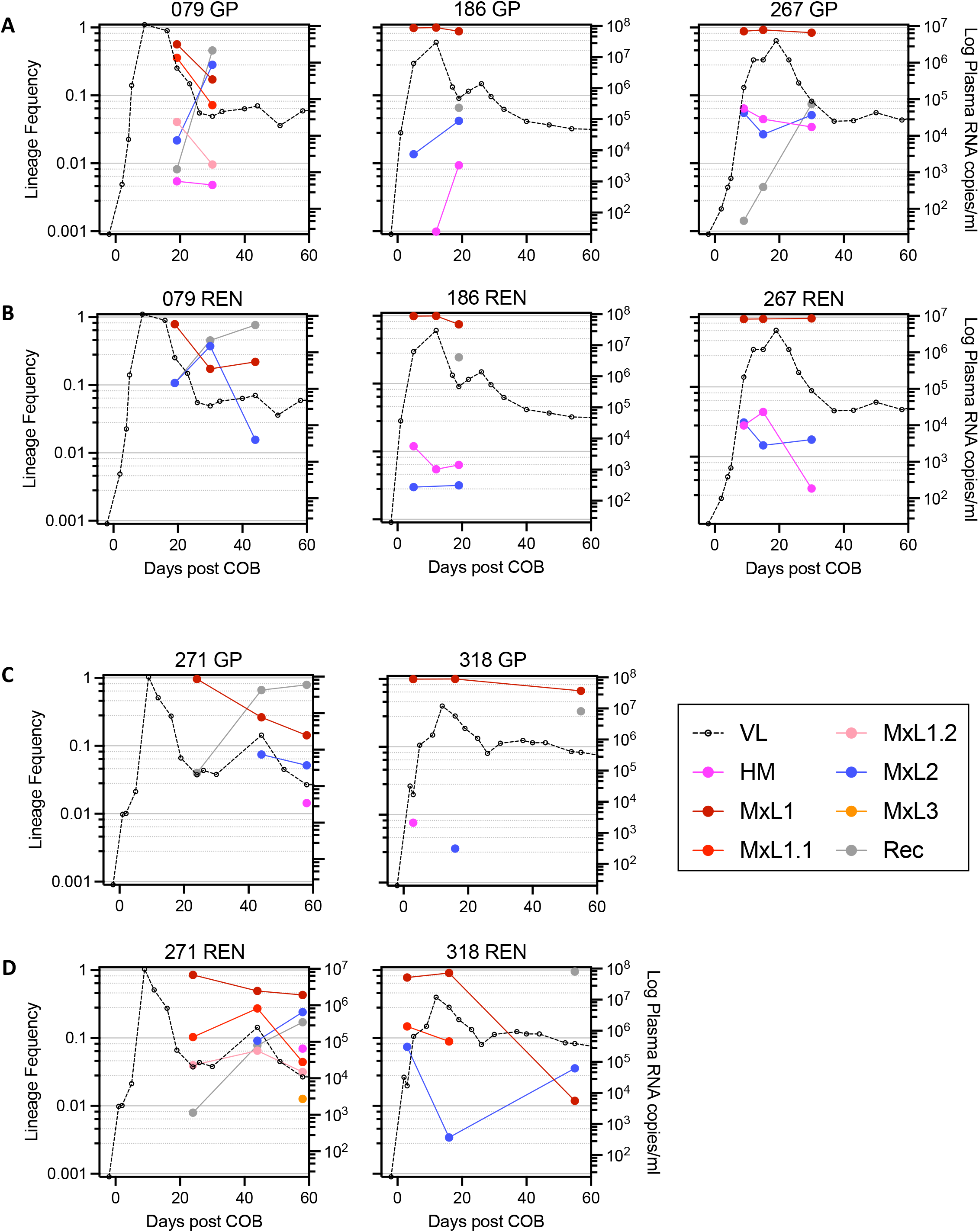
Plasma HIV RNA viral load and lineage frequencies over time. Data from the 5 participants from the FRESH cohort with multiple transmitted lineages are shown with stacked panels showing GP (**A, C**) and REN data (**B, D**), respectively. Key abbreviations: VL, plasma HIV RNA; HM, hypermutated sequences; MxL(n), lineage designations with numbers (n) assigned in decreasing order of the abundance of the lineages at the 1^st^ time point; “x” is used instead of a given stage of infection since data from all time points are shown; MxL(n.m), UOL sequences closely related to MxL(n), numbered (m) in order of frequency at the first time point; REC, recombinants.

Analysis of pairwise maximum likelihood distances, as noted in previous studies^12,22^, showed that the diversity of individual viral lineages often contracted relative to the first time point at either the second or third time point sampled, especially in the RV217 cohort (Supplementary Fig. 10), which on average was initially sampled earliest in infection (Fig. 1). The diversity of the *gag* and *env* lineages within each individual were evaluated by calculating Shannon entropies. Average entropy values were determined across both the *gag* and *env* genes and deduced viral proteins. Sliding window displays of entropy variation comparing genes and proteins were similar (Supplementary Fig. 11) and for subsequent analyses only amino acid data will be discussed. Because averages can be substantially impacted by outliers within small datasets, we restricted the following analyses to datasets comprised of at least 10 members in each lineage. For the same reason, we employed a second measure, the proportion of lineages that had nonzero entropy at each position. The two types of entropy plots showed diversity in the same regions, but with different relative amplitudes in some regions (Supplementary Fig. 11). Similar patterns were observed when comparing entropies in viral Subtype C vs other “Not Subtype C” subtypes (Supplementary Figs. 12, 13); the number of samples of Non-C subtypes of sufficient lineage size was judged to be too few to assess by individual subtype).

In addition to entropy we also identified amino acid sites that were inferred to be undergoing pervasive negative (purifying) or positive (diversifying) selection across the Gag and Env genes, again pooling data from all cohorts and lineages. Overall, 7 fold more sites were inferred to be undergoing purifying versus diversifying selection in Gag, and 1.5 fold more in Env (Supplementary Table 5). Next, we examined selection in the 6 individual coding regions within Gag and after separating Env into 3 regions: gp120 and the extracellular and intracellular regions of gp41. In addition, we split the datasets into four time-based stages following COB/EDDI as noted in Fig. 1 and Supplementary Table 6. The earliest ‘i’ stage was a period of exponential viral load increase. Stage ‘ii’ corresponded to a period of rapid decrease in viral load, ‘iii’ corresponded to a slower decrease in viral load and ‘iv’ the beginning of a period of approximately steady state viral load. Stage ‘iv’ included only AMP trial participants, had the least available data (Fig. 1), and thus will not be discussed further. Adaptive immune responses leading to positive selection and immune escape are typically first detected in stage ‘ii’, with escape mutations first noted at one or very few sites within the viral proteome in stages ‘ii’ and “iii”^23-25^.

The number of sites inferred to be undergoing negative selection increased through the 3 stages in nearly all Gag and Env coding regions. Positive selection was less consistent, with the highest levels reached in the intracellular segment of gp41 within Env, p2 within Gag, and gp120 within Env. The ratio of negative to positively selected sites was substantially higher in the p24 than other coding regions (Fig. 3A-B). Interestingly, this was not due to a higher level of negative selection, but rather an atypically low level of positive selection (Fig 3C). The greater negative to positive selection ratio in Gag vs. Env was associated with both lower levels of negatively selected sites and higher levels of positively selected sites (Supplementary Table 5).

**Fig 3.**
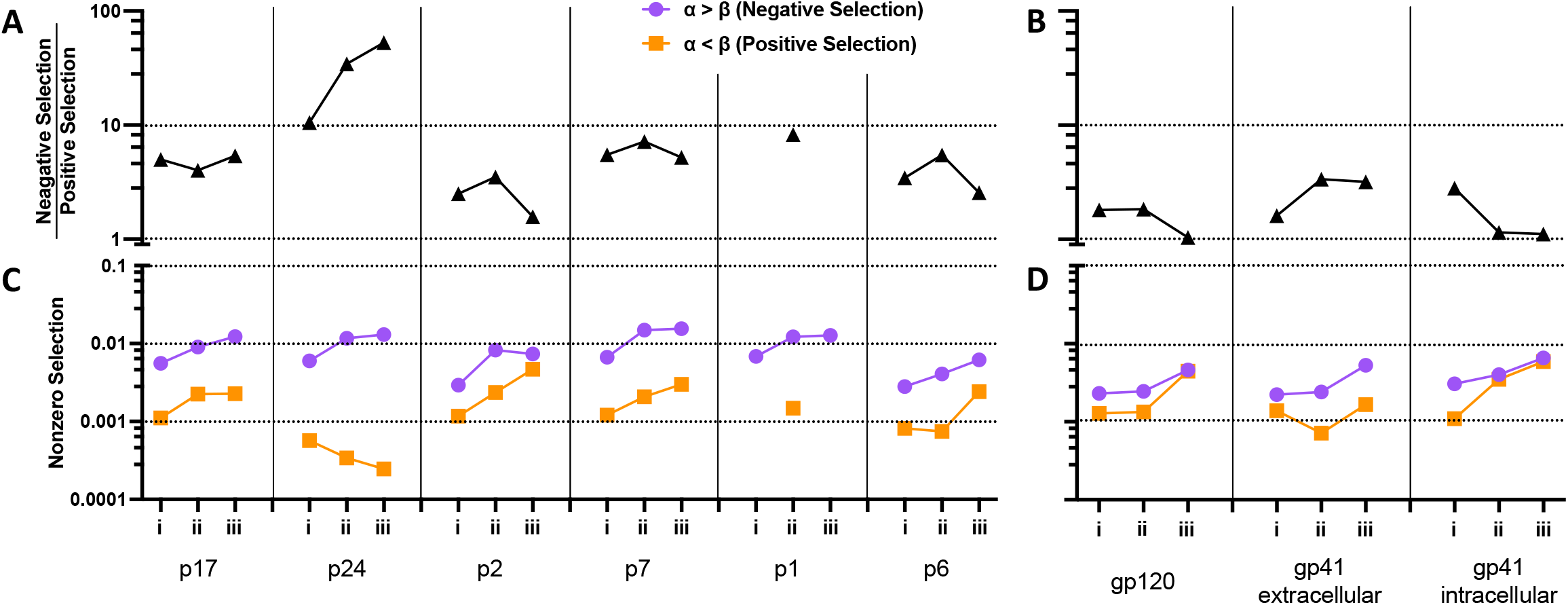
Inferred selection in Gag and Env coding regions as a function of stage of acute/early HIV infection. The top panels show the ratio of negatively and positively selected sites across separate coding regions of Gag (**A**) and Env (**B**), respectively. Panels **C** and **D** show the frequencies of negatively and positively selected sites separately.

We next assessed entropy and selected sites along each coding region and through post COB/EDDI stages i-iii, combining data from all time points, participants and lineages. Entropy was evident within both Gag and Env at the initial sampling times. Regions of higher entropy were found near the N and C termini of p17 (matrix protein), in p2-p7(nucleocapsid)-p1 and the C-terminal portion of p6 in Gag (Fig 4A). These levels were substantially driven by changes in stage iii in p17, p7 and portions of p6 (Fig. 4C). Positive selection was most evident in stage iii in the N-terminal region of p17, p2, the N-terminal region of p7 and segments within p6, and in stage ii in regions of p17 and p7 (Fig. 4E). Interestingly, positive selection was focused on the N-terminal region of p7, whereas negative selection peaked toward the C-terminal region of p7 (Fig. 4D).

**Fig 4.**
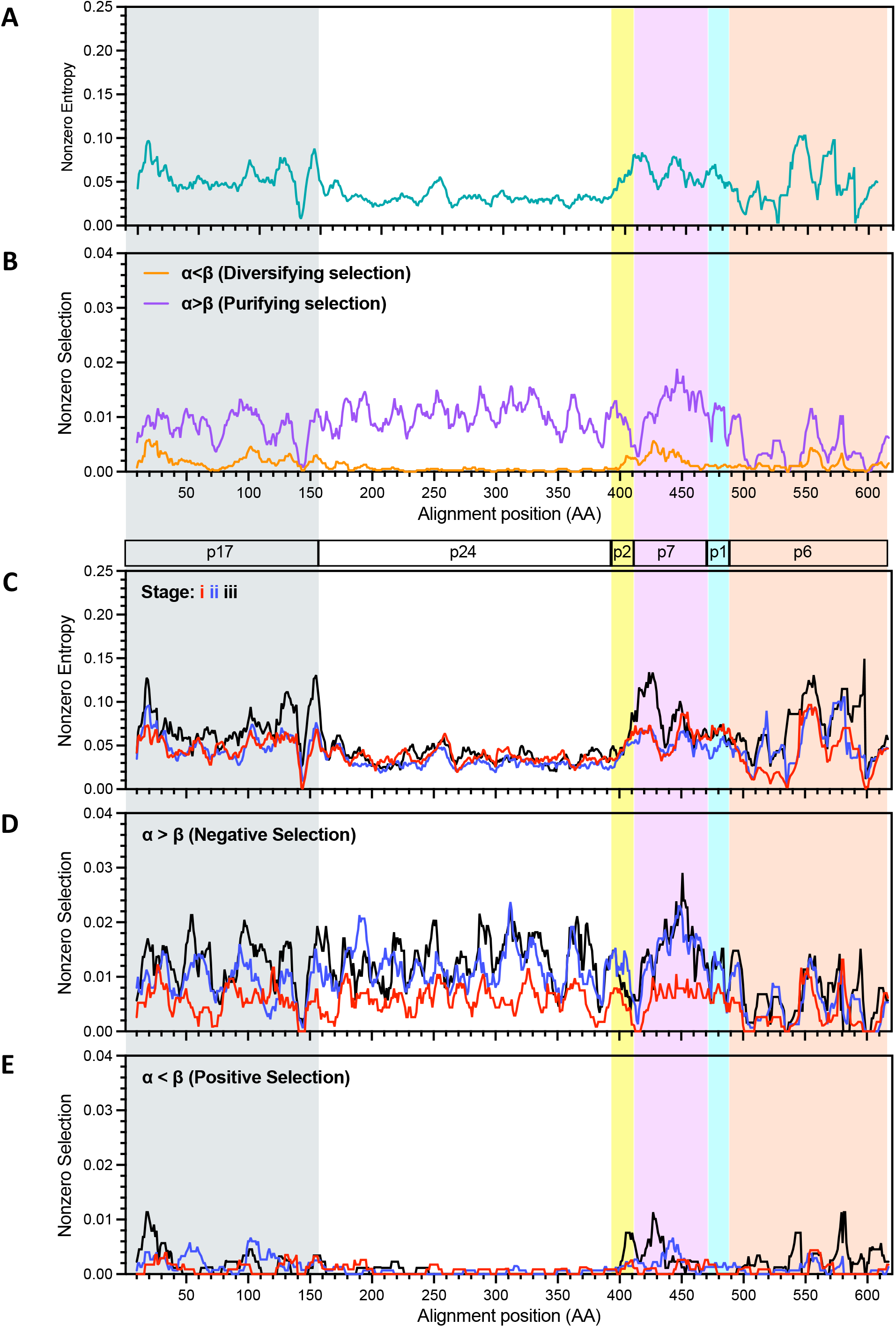
Entropy and selection within Gag. Plots of the proportion of lineages with nonzero entropy (**A**) and (**C**) and selection (**B, D** and **E)** derived from an alignment from all lineages having at least 10 members and plotted in sliding windows of 10 amino acids. Identity of coding regions are indicated within bars above panel **C** and shaded with different colors. Panels **A** and **B** show data from all time points combined. Panel C shows nonzero entropy, panel D negative selection and E positive selection from each of the three stages of acute/early HIV infection. Data included derives from all lineages with at least 10 members. Stage i, red lines; ii, blue; iii, black.

**Fig 5.**
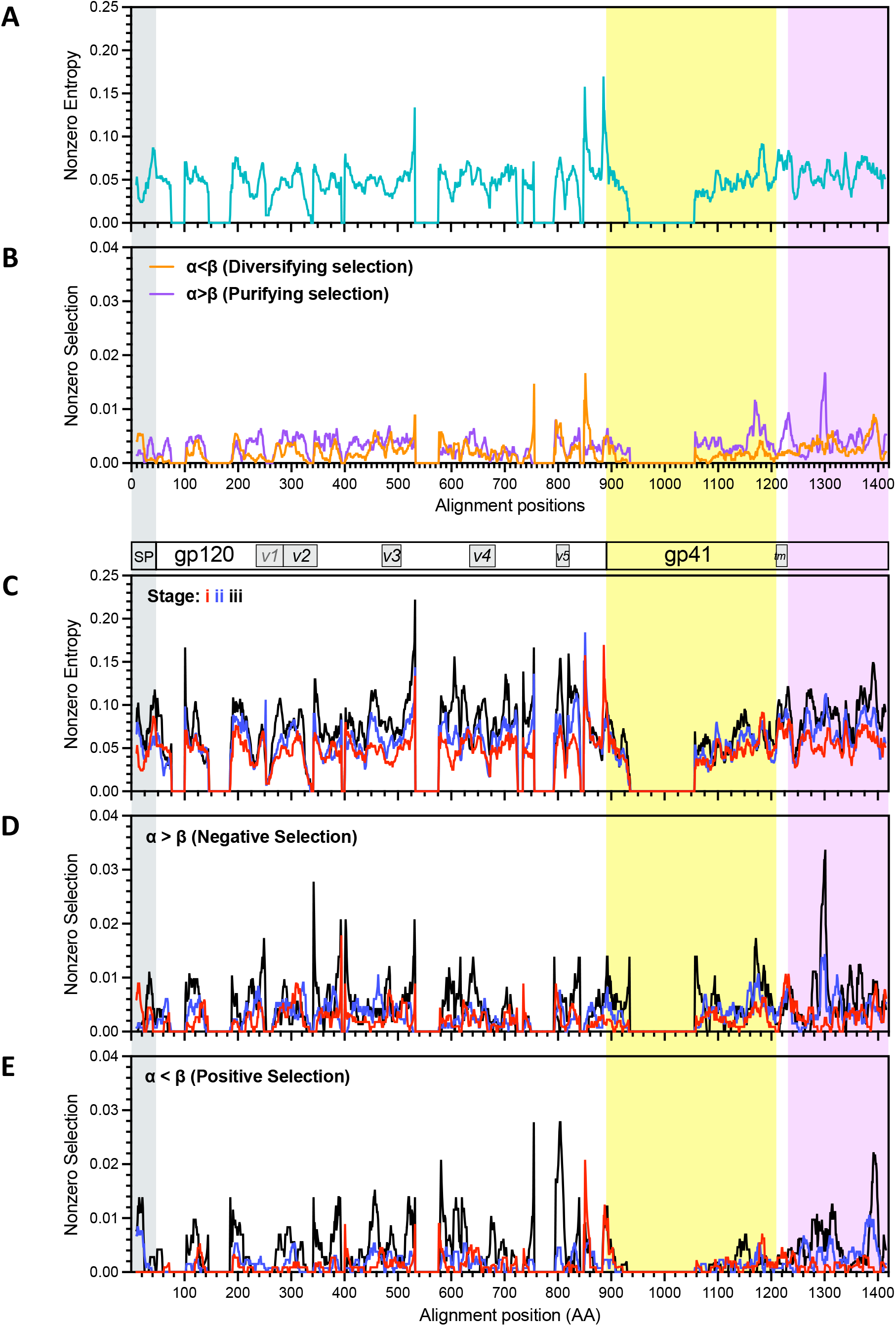
Entropy and selection within Env. Plots of the proportion of lineages with nonzero entropy (**A**) and (**C**) and selection (**B, D** and **E)** derived from an alignment from all lineages having at least 10 members and plotted in sliding windows of 10 amino acids. Identity of coding regions are indicated within bars above panel **C** and shaded with different colors. Panels **A** and **B** show data from all time points combined. Panel C shows nonzero entropy, panel D negative selection and E positive selection from each of the three stages of acute/early HIV infection. Data included derives from all lineages with at least 10 members. Stage i, red lines; ii, blue; iii, black.

Entropy as well as selected sites were more distributed over Env, with several peaks of positive and negative selection. The positive selection noted in the cytoplasmic domain of gp41 (Fig. 3) was associated with peaks in the central and C-terminal regions.

In a study of HIV subtype C infection of donor-recipient pairs, the envelope genes in recipients were found to have shorter variable loop lengths and fewer potential N-linked glycosylation sites (PNGS)^10^ although this was less consistent for donor-recipient pairs infected with HIV Subtype B^26,27^. Here too, subtype C viruses had variable regions that were shorter than other subtypes, and shorter than those found in the LANL database, the latter of which have sequences from viruses from acute as well as chronic infection and AIDS (Supplementary Table 7). Also consistent with prior studies, subtype B viruses had the longest variable loops and were similar to or longer than those found in the LANL database. A previous study of the RV217 cohort failed to demonstrate a change in PNGS over the first 6 months of infection^28^. Here too, the median number of PNGS did not vary through stages i-iii (Supplementary Figure 14), although subtype C viruses had on average one fewer site vs Subtype B. We also tallied the sites within and nearby NLGS sequons that experienced positive and negative selection. Neither sequons nor the 9 amino acid regions centered on sequons had a higher level of selected sites compared to random expectations, although sequons and immediately N-terminal and C-terminal amino acids appeared to be enriched for negatively selected sites (Supplementary Fig. 15) and a significant difference between the ratio of T vs S was noted when comparing positive and negative selection levels in the 3^rd^ position (p = 0.03941, OR = 3.33, 95% CI = (1.1, 11.4), 2-way exact Fisher test).

## Discussion

This study offers an unparalleled view into the dynamics of HIV viral populations during early infection and provides a glimpse into the processes that underly the establishment and early maintenance of HIV infection. Our advanced methods detected a higher frequency of transmission of multiple distinct virus lineages compared to previous studies^14^. A major strength of our study derives from sampling: larger numbers of sequences, i.e., sampling deeper to identify minor variants; over larger stretches of the genome, which allows greater opportunity for detecting distinguishing mutations, and; over multiple time points, which increased the number of sequences evaluated and allowed for detection of potentially late appearing lineages. This also allowed opportunities to view dynamic changes in variant detection and representation.

Massive virus population expansion occurs during the approximately 7-day eclipse phase of infection prior to detectability and continuing during initial acute infection^19^. Multiple forces may shape these emerging virus populations very early in infection, including: the stochastic process of variants entering cells with different division rates; inherent variant replication rate; purifying selection for more rapid growth, including possible loss of mutations selected for immune escape in the prior host that also caused a loss in replication fitness, and; diversifying selection for the emergence of escape mutants from adaptive immunity^11,12,29-31^.

The significant increase in the percentage of multilineage acquisitions from approximately 25% (95% CI of 21-29%) in a prior meta-analysis^14^ to a minimum 38% (95% CI of 30-47%) reported here, was largely due to the detection of minor variant lineages, and there was no evident bias associated with sex. However, 38% is very likely an underestimate due to multiple factors: **1)** Rapid and large changes in lineage representation. We found major shifts in lineage representation over the first ∼2 months of infection in two-thirds of the multilineage acquisitions in the RV217 and FRESH cohorts. These two cohorts were sampled at high temporal resolution and the intensive sampling provided a uniquely detailed view of viral dynamics, offering insights that were previously inaccessible due to less frequent sampling. However, in one case (RV217 participant 40061 from Thailand), a previous study using short-read length sequences from the same individual found that a minor variant at day 14 was dominant between days 19 and 28 and not detected at day 42^32^ (Supplementary Fig. 15C). In both the previous and current studies, this lineage represented about 25% of the sequences at day 14. but in the current study no time points were observed when this lineage dominated (Supplementary Fig. 15A-B). This illustrates that even with the closely spaced sampling we employed, major transient shifts in virus populations may fail to be observed, including the failure to detect variants that at other times may appear to be the founder of infection. **2)** Shallow sampling relative to the *in vivo* population. Any feasible scale of virus population sampling is very shallow relative to the *in vivo* population, and we found that variants that only appeared subsequent to the first time point tended to have lower numbers of sequences recovered at the first time point compared to the individuals in which multiple lineages were detected at the first time point. We estimate that the average number of sequences we obtained using the SMRT-UMI method is likely to detect 99-100% of all variants with a frequency of at least 5% whereas when using Sanger SGA to obtain 20 sequences, variants present at 5% are missed 36% of the time. HIV sequences from the RV217 and FRESH participants studied here were previously determined using Sanger SGA and in the case of the Thailand cohort, 5/16 (31%) of individuals were found to have MLA^19^, compared to 50% in the current study. The same percentage of participants from the Kenyan cohort (2/13, 15%) were found to have MLA by both Sanger SGA^19^ and here using SMRT-UMI. In the case of the FRESH cohort, only one of the 13 individuals (8%) were found to have MLA by Sanger SGA, compared to 5/13 (38%) here (Ndung’u et al, unpublished). **3)** The presence of lineages of uncertain origin (UOL), as identified here in 27% of what we have characterized as single lineage acquisitions, could increase the tally of multilineage acquisitions to as high as 53% (95% CI 46-62%) in our datasets. UOL may represent early evolution or transmission of multiple lineages from a newly infected donor with a near homogeneous viral population. **4)** A large fraction of transmissions occur when the transmitter is in acute infection^33,34^ when the infecting population often has little diversity. Hence, not all multilineage transmissions would be discernable by viral genetic analysis in these cases. **5)** We studied 5.5kb of the 9kb viral genome, whereas inclusion of the remaining 40% of the genome may have identified additional lineages.

Recognizing that most individuals are sampled only once early in infection and precise staging (e.g., Fiebig^35^ or other staging^21,36,37^) is often not be possible, nuance is appropriate in labelling virus lineages. In the current study of RV217 and FRESH cohort individuals, for which the threshold of RNA positivity is known within a few days, the lineages observed were in the earliest recognizable stages following HIV acquisition. We propose that the largely homogeneous variant populations observed very early in infection should nonetheless be considered representatives of a transmitted lineage, rather than definitively characterized as the actual transmitted variant(s). As such, the common term “transmitted/founder”^25^ or T/F lacks precision, in part because it combines two different properties that are not necessarily linked: a “transmitted” variant does not necessarily indicate the “founder” of infection over the long term, and there is no clear definition implied of how long after infection a “transmitted” lineage is discernible. Simiarly, “founder”^38^ can be misleading since in some cases the major variant(s), i.e., the presumed founder, appears different depending on the precise timing of sampling. Another rationale for use of “lineage” comes from the observation that entropy and selection on the virus population was observed at the very earliest times of infection (i.e., stage ‘i’). As time of infection progresses, detection of transmitted variants and even lineages becomes harder because of selection (both positive and negative) and recombination, and our inability to discern complex recombination patterns. From this conceptual advancement we propose the use of the term “transmitted founder lineages” (TFL) to describe virus populations detected early in infection.

Two observations were made that may be associated with viral subtype. First, subtype C virus populations were found to harbor significantly more defective *gag* and *env* genes than other subtypes, but this was not due to higher rates of APOBEC-mediated hypermutation which frequently leads to the formation of stop codons. This suggests that the fidelity of reverse transcriptase (RT) is lower in subtype C viruses, although very limited *in vitro* studies comparing the fidelity of subtype B and C viruses have not shown differences^39^. Given the variability in the levels of defective viruses in the infections we studied, a comprehensive study that examines multiple representatives of each of the subtypes of interest is warranted. Second, atypically low levels of MLA were found in the RV217 Kenyan female cohort. While the reason for this is unclear, it cannot be attributed simply to sex, as the Kenyan as well as individuals from the FRESH and HVTN 703 cohorts were assigned as female at birth. Some differences in mode of transmission in the Kenyan vs. the FRESH and HVTN 703 cohorts (e.g., receptive vaginal vs. anal intercourse) cannot be ruled out. However, there was a difference in subtypes – nearly all acquisitions were with subtype C in the FRESH and HVTN 703 cohorts whereas 11/13 individuals in the Kenyan cohort were subtype A1 or recombinants involving subtype A1.

Glycosylation can negatively affect the rate of protein folding^40^ and importantly, can impact viral infectivity and provide a shield against antibody recognition^41-43^. Thus, selection for the loss of PNGS may occur in the absence of neutralizing antibody responses in acute infection, as a more accessible and compact envelope protein may increase viral replication fitness. Consistent with this hypothesis was the finding that negative selection was associated with a higher prevalence of S over T at the 3^rd^ position of sequons, preserving the sequon, and a higher prevalence of T over S in sequons that were lost as a result of positive selection at this site. Since glycosylation is up to 40 times more likely to occur when T is in the 3^rd^ position^44,45^, both of the differences noted favor unglycosylated sequons, and increased viral fitness in the absence of adaptive immunity.

Overall, we found MLA in at least 37% (CI 26-50) of male-female transmissions and 39% (CI 28-51%) of male-to-male transmissions, both potentially higher and with less sex differences than reported in the previous meta-analysis (21%, 14-31 and 30%, 22-40%, respectively)^14^.

The potential implications of this study for HIV prevention are threefold. Given that we consider it probable that substantial numbers of variants are transmitted in a large fraction of transmissions, what we observe may only be the transient winners of the race to detectability. This may help explain the difficulty in making effective HIV vaccines beyond the problem of the increasing the global diversity of HIV^46,47^, since a diverse transmitting population has a greater likelihood of harboring variants capable of escaping blockades the vaccines produce. This is consistent with the findings from large scale HIV vaccine trials in which infection with viruses more dissimilar to the vaccine are most likely to result in virus infection^48-50^.

Second, sites of strong selection, in particular purifying selection, have been a focus for vaccine antigen designs for the major viral proteins Gag, Pol and Env^51-53^. The strong conservation of p24, in particular (and reinforced in this study), has made this protein a target for research and development of a highly potent antiretroviral drug^54,55^ with particularly conserved regions of p24 used as vaccine candidates^56-61^. Interventions that effectively target conserved regions of the viral proteome may enhance vaccine efficacy due to their strict structural requirements for viral infectivity^62^. This study identified additional regions of Gag proteins to be undergoing negative selection, e.g., the central region of p17 and C-terminus of p7, and thus potential vaccine targets.

Third, the intracellular region C-terminal to the transmembrane sequence, p2, the N-terminal region of p7 and much of p6 were relatively strong targets of positive selection in the current study. As immunologically dominant epitopes can act as decoys that prevent immune recognition of vulnerable features of the viral proteome^63-66^, vaccines that target subdominant, conserved features of the viral proteome^52,53,65,67-69^ critical to viral fitness^62,70,71^ may benefit from the omission of these coding regions.

## Online Methods

### Study Subjects and specimens

Individuals included in this study were derived from 4 prospective cohorts of HIV acquisition (Table 1). The RV217 study included a cohort of males and transgender females (TGF) from Thailand and a cohort of females from Kenya^16^. The Females Rising through Education, Support, and Health (FRESH)^17^ cohort included females from KwaZulu-Natal, South Africa. Individuals were prospectively identified in acute HIV infection by twice-a-week plasma RNA testing with the date of detectable acquisition taken to be the center of bounds (COB) between the last negative and first positive HIV RNA test^20^. A subset of individuals from these cohorts were chosen for study based on plasma specimen availability and samples were chosen to match the predicted estimated date of detectable infection (EDDI) in individuals in the AMP trials. Plasma samples used for viral genome sequencing were taken from multiple time points prior to initiation of antiretroviral therapy (ART).

Two cohorts corresponded to the control arms receiving placebo only in the Antibody Mediated Prevention (AMP) clinical trials HVTN 703/HPTN081 (females from southern Africa, abbreviated HVTN 703) and HVTN 704/HPTN085 (MSM and transgender females from the Americas and Switzerland, abbreviated HVTN 704)^1^. AMP trial participants were monitored monthly and often retested 1-2 weeks following an initial viral RNA positive finding. At least the first 2 RNA positive visits were chosen for sequencing. Estimates of their EDDI^37^ were derived using both clinical diagnostic and viral sequencing data corresponding to a preliminary dataset of the sequences reported hered^20^ (see below).

### SMRT-UMI sequencing

The Pacific Biosciences single molecule real-time (SMRT) platform was used to sequence 2.5kb PCR amplicons encompassing the HIV *gag* and part of the *pol* gene (GP region), and 3kb amplicons from *rev* through *env* and a portion of the *nef* gene (REN region) (S1 Fig), with samples split between the University of Washington and University of Cape Town laboratories. Each sequence was derived from individual cDNA templates amplified by single genome amplification (SGA) and tagged with unique index adaptors for sequencing or, in most cases, tagged with unique molecular identifiers (UMI) during cDNA synthesis and amplified in bulk (SMRT-UMI)^15^. Both protocols resulted in accurate single-molecule sequencing and viral RNA template quantitation^15^. Sequencing was performed using the Pacific BioSciences Sequel and Sequel IIe instruments followed by demultiplexing and processing into consensus sequences for each viral template using the PORPIDpipeline^15^ for amplicons amplified using the SMRT-UMI approach or with a simplified pipeline (https://github.com/MullinsLab/sga_index_consensus) to demultiplex and generate a consensus from SGA. For samples with viral loads above 20,000 HIV RNA copies/ml of plasma, the concentration of amplifiable target amplicons were initially estimated using end-point dilution (EPD) nested PCR and the Quality tool (https://quality.fredhutch.org)72 with 3 replicates each at 5-6 dilutions estimated to reach an endpoint based on clinical viral loads. Westfall et al^15^ estimated PCR and sequencing error rates using this method paired with the PORPIDpipeline software to be less than 8.6 × 10^-8^ per base, or less than one error in every ∼3700 REN sequences. To minimize recombination during PCR, no more than 25 amplifiable copies of cDNA was added to each reaction. PCR recombination was observed if the number of cDNA molecules in a PCR was above 100.

Given an inherently large standard error in the quantitation measurements and with a target of obtaining at least 100 sequences per sample, 200, and in later experiments 250 templates, were targeted for amplification. Amplifiable templates in samples with viral loads between 1,000 & 20,000 copies/ml were not quantified by EPD PCR but rather the cDNA (derived from up to 1mL of plasma) was divided into 20 nested PCR reactions and only those PCR that were positive following a gel screen were processed using the SMRT-UMI protocol. SGA was performed for any sample with a viral load below 1,000 copies/ml of plasma.

The SMRT-UMI laboratory protocol resulted in carryover of a small amount of UMI-containing cDNA primers. This had the effect of a small number of cDNA primers being used as primers in subsequent PCR steps, thus artificially inflating the number of templates recovered. Sequences derived from these carryover cDNA primers typically had very small family sizes (a “family” corresponds to a group of sequences, each containing the same UMI). Following a series of experiments to quantify cDNA carryover we conservatively “cleaned” the datasets by removal of sequences with small family sizes below a threshold corresponding to 15% of the total number of reads (or 20% for the HVTN 703 REN datasets). This approach removed some of the “real” sequences but removed nearly all of the sequences derived from cDNA carryover. Sequences were checked for contamination against a laboratory database before nucleotide and amino acid alignment using BLAST^73^. Final sequences were deposited in GenBank after collapsing identical sequences, with Accession numbers as follows: FRESH GP (PV963961 - PV968620), FRESH REN (PV968621 – PV972048), RV217 GP (PX001706 – PX010774), RV217 REN (PX010775 - PX020890), HVTN 703 GP (PX030312 – PX031976, PX038076 – PX042058), HVTN 703 REN (PX072574 – PX077815), HVTN 704 GP (PX154055 – PX161466), and HVTN 704 REN (PX172337-PX183870).

Viral subtypes were determined using the Recombinant Identification Program (RIP,^74^, https://www.hiv.lanl.gov/content/sequence/RIP/RIP.html). When subtype assignments were unclear, the REGA HIV subtyping tool (https://www.genomedetective.com/app/typingtool/hiv) and the National Center for Biotechnology Information (NCBI) genotyping tool (https://www.ncbi.nlm.nih.gov/projects/genotyping/formpage.cgi) were also employed.

### Sequence alignments

Nucleotide alignments were initially generated using the MUSCLE algorithm^75^ version 3.8.31. To reduce the number of taxa in alignments, identical sequences were collapsed using a python script https://github.com/MullinsLab/sequence_collapsing. Three rounds of manual review and refinement were then conducted by three different experienced scientists. In the first round of review, codon position was used to assist placement of gaps, but otherwise alignments were not codon-optimized but rather optimized for sequence homology. Subsequently, a second scientist reviewed the refined alignments and any changes noted and discussed with the primary reviewer. Finally, a third scientist assessed the alignments and any further edits noted, with each alignment finalized in consultation with the primary and secondary reviewers.

### Lineage assignments

Lineages were assigned to each sequence within an individual using Poisson Fitter^21^ with sequences from the first available time point, and an iterative diversity/phylogenetic approach using sequences pooled from all time points. In the iterative approach, maximum likelihood phylogenetic trees and highlighter and match plots were generated for GP and REN regions using an in-house pipeline (https://github.com/MullinsLab/phylobook_pipeline) and displayed within the Phylobook tool^76^. Within Phylobook, clustering algorithms were used to help assign lineages along with manual review and selection. After several iterations and reviews, variants with a cluster of 4 or more shared nucleotide positions were found to categorize sequences into distinct lineages, judged to have been derived by the transmission and outgrowth of distinct variants. Variants with scattered unique changes were not characterized as being transmitted. The origin of lineages detected at the first time point and differing by 1-3 nucleotides were judged to be uncertain (i.e., uncertain origin lineages, UOL). For the comparison of distance distributions, UOL sequences were included within the most closely related lineage(s). Not counted in the identification of UOL were positions at which more than 2 prevalent nucleotide states were found, as this is likely indicative of diversifying selection.

Recombinant sequences were identified by manual inspection as having at least 2 nucleotide changes matching another lineage. As the structures of many recombinants were quite complex, no effort was made to separate them into distinct lineages. To assess lineage assignments, hypermutated sequences were removed using an R-script based on the LANL Hypermut tool^77^, and then maximum likelihood pairwise distance distributions generated within and between all lineages using PhyML v3.3^78^ within the DIVEIN suite (https://divein.fredhutch.org)^79^. Distance distributions from within- and between-lineages were compared and any values that overlapped the two distributions re-evaluated in Phylobook for proper lineage assignment. Following initial lineage assignment, a second experienced reviewer assessed assignments and any discrepancies discussed and assignments amended if needed. PhyML within DIVEIN was also used for phylogenetic tree generation using the GTR substitution model, an estimated gamma distribution and optimization for both topology and branch lengths.

### Statistical Analyses

Differences between genes and time points in the frequency of intact genes were assessed using both a parametric method, which allowed us to combine all variable into a single model, and a non-parametric method, by running a Wilcoxon test comparing each variable within each subgroup (gene/time point/subtype) and then applying an FDR threshold of 0.2 to correct for multiple testing (Supplemental Table 4). For the parametric method, we ran a random effects generalized linear model (GLM) with the logit transformed frequency as dependent variable, gene (either *gag* or *env*) and subtype (categorized as C, B or “other” for all other subtypes and including all intersubtype recombinants) as independent variables, and participant ID as a random effect. To make days since COB comparable across cohorts, we chose times since COB/EDDI to be as uniform as possible across participants, with each study participant represented only once for each model run. As such, we identified two time points (“windows”) with mean times since COB/EDDI of 30 and 45 days, respectively, selected as follows: **1)** Sequence datasets from approximately 30 days post COB/EDDI, including all first time point samples from V703 and V704, the second time point for FRESH, and the third time point for RV217, as for both the FRESH and RV217 the first time point was on average sampled around two weeks earlier than the first time point in the AMP cohorts (Supplementary Table 3). **2)** Sequence datasets from approximately 45 days post COB/EDDI, including the second time point for V703 and V704, the third for FRESH and the fourth for RV217. Associations between viral loads and fraction of intact sequences and hypermutated sequences were assessed using the Kendall correlation coefficient and test. These analyses were computed in R (version 4.2.1) using packages lme4, LaplacesDemon, and pbkrtest. Confidence intervals were calculated using the binomial (Wilson score interval) method. The power to detect minor variants by frequency and number of sampled sequences was calculated using the binomial function (pbinom) in R Studio.

### Measurement of entropy and selection

The entropy found in each lineage, with all timepoints combined as well as during each of the i-iv stages, were calculated using the R script https://github.com/MiguelMSandin/DNA-alignment-entropy modified to perform calculations in batch. Selection was similarly calculated using the FUBAR algorithm modified to perform calculations in batch (https://github.com/MullinsLab/FUBAR_in_batch). For both entropy and selection analyses, data was then summed at each nucleotide or amino acid position, with sequences placed in register from an inter-participant alignment. Python-and R-scripts were used for data processing and rendered in Prism (GraphPad Software, Inc).

## Acknowledgements

We gratefully acknowledge Kim Wong for technical assistance and the dedicated participation of the many individuals in the FRESH, RV217 and AMP trials cohorts, and the UKZN HIV Pathogenesis Programme laboratory staff without whom none of this work would be possible. This work was supported by the Bill and Melinda Gates Foundation (Investment Record ID INV-016189 to JIM; We thank Dr. Thandi Onami for critical guidance) and by Public Health Service Grants (UM1 AI068614, to the HIV Vaccine Trials Network [HVTN]; UM1 AI068635, to the HVTN Statistical Data Management Center, and; UM1 AI068618, to the HVTN Laboratory Center) from the National Institute of Allergy and Infectious Diseases (NIAID) of the National Institutes of Health (NIH). The content is solely the responsibility of the authors and does not necessarily represent the official views of the NIH. The FRESH program was supported in part by grants from the Bill & Melinda Gates Foundation, the International AIDS Vaccine Initiative (UKZNRSA1001), the Harvard University CFAR grant (P30 AI060354), the Witten Family Foundation, Dan and Marjorie Sullivan, the Mark and Lisa Schwartz Foundation, Ursula Brunner, AIDS Healthcare Foundation, and the Howard Hughes Medical Institute. TN was further supported through the Sub-Saharan African Network for TB/HIV Research Excellence (SANTHE) which is funded by the Science for Africa Foundation to the Developing Excellence in Leadership, Training and Science in Africa (DELTAS Africa) program [Del-22-007] with support from Wellcome Trust and the UK Foreign, Commonwealth & Development Office and is part of the EDCPT2 program supported by the European Union; the Bill & Melinda Gates Foundation [INV-033558]; and Gilead Sciences Inc. [19275]. MR, MLR, FS, and SN received support from a cooperative agreement (W81XWH-11-2-0174) between the Henry M. Jackson Foundation for the Advancement of Military Medicine and the US Department of Defense.

## Ethical statement

The work described here complied with all relevant ethical regulations. The Institutional Review Boards/Ethic Committees of participating clinical research sites (CRS) approved the studies, which were conducted under the oversight of the NIAID Data Safety Monitoring Board^1^. Viral genome sequencing at the University of Cape Town was approved by the UCT Human Research Ethics Committee (HREC reference no. 176/2017) and was considered exempt at the University of Washington.

