## Supplement for "Long-Read Deep Sequencing Reveals High Rates of Multilineage Transmission and Rapid Viral Population Changes in Acute HIV Infection"

**Supplementary Discussion**

1. Interestingly, in the second time window evaluated (See Online Methods), the frequencies of hypermutated sequences became higher in *gag* compared to *env* for Subtype C, whereas for the other subtypes it remained higher in *env*. Comparisons of the frequencies of hypermutated sequences in the second time window across clades were not significant in *gag* and they were significant in *env* only when comparing subtype B to subtype C (Lower in Subtype C; Supplementary Table 4C).

2. There was a significant correlation between the proportion of hypermutated sequences and viral load in *gag* (p=0.03) but not in *env* (p=0.9) in the first time window, and no significant associations were found in the second time window.

3. This method derives from the understanding that prior to the onset of detectable immune-driven selection in the viral proteome (~5-7 weeks post-acquisition; Goonetilleke N, et al. J. Exp. Med. 2009;206(6):1253–72; Liu, Y. et al. J. Virol. 2006;80(19):9519–29; Liu Y, et al. Virology 2006;347(1):140–6), an infection initiated by a single variant is expected to grow exponentially, randomly accumulating mutations, with mutational distances fitting a Poisson distribution. Failure to fit a Poisson indicates evidence of selection or acquisition of multiple lineages. Subsequent partitioning into groupings that each fit a Poisson resulted in the definition of distinct lineages from within these multilineage infections.

| Supplementary Table 1. HIV subtypes and discordance between GP and REN subtypes | | |
|---|---|---|
| **Cohort** | **REN Subtypes** | **GP-REN Subtype discordance[2]** |
| **RV217 (Thailand)** | 15x CRF01AE, 1x B | 1x B-AE |
| **RV217 (Kenya)** | 3x A1, 3x A1C, 5x A1D, 1x C, 1x D | 1x D-A1D, 1x A1-A1C, 2x A1-A1D, 1x A1C-A1D, 1x A1C-D |
| **FRESH** | 13x C | |
| **HVTN 703[1]** | 31x C, 1x G | 1x A1G-G |
| **HVTN 704[1]** | 37x B, 4x F1, 1x F2, 5x BF1, 1x A1B | 1x BF1A1-BF1, 1x B-BF1, 2x BF1-F1 |
| [1]From the trial control arms (placebo recipients) only. [2]Numbers and identities of discordant subtype determinations are listed for GP-REN. | | |

**Supplementary Table 2. Putative inactivating mutations**

#### Gag

| Cohort | Proportion intact | Putative Inactivating mutations (Proportions)[1] | | | |
|---|---|---|---|---|---|
| | | Premature stop | Large deletion[2] | Missing 5' end[3] | No ATG[4] |
| FRESH | 0.905 | 1 | 0 | 0 | 0 |
| RV217 Kenya | 0.950 | 0.99 | 0.003 | 0.0001 | 0.005 |
| RV217 Thailand | 0.964 | 1 | 0 | 0 | 0 |
| HVTN 703 (C) | 0.879 | 0.997 | 0.001 | 0 | 0.002 |
| HVTN 704 (C) | 0.977 | 0.987 | 0 | 0 | 0.013 |
| *Overall* | *0.943* | | | | |

#### Env

| Cohort | Proportion intact | Putative Inactivating mutations (Proportions)[1] | | | |
|---|---|---|---|---|---|
| | | Premature stop | Large deletion[2] | Missing 5' end[3] | No ATG[4] |
| FRESH | 0.821 | 0.986 | 0.003 | 0.01 | 0.0002 |
| RV217 Kenya | 0.847 | 0.989 | 0.003 | 0.004 | 0.003 |
| RV217 Thailand | 0.843 | 0.972 | 0.017 | 0.004 | 0.007 |
| HVTN 703 (C) | 0.789 | 0.968 | 0.008 | 0.019 | 0.004 |
| HVTN 704 (C) | 0.903 | 0.982 | 0.007 | 0.0006 | 0.004 |
| *Overall* | *0.856* | | | | |

[1] These proportions refer to the composition of the sequences that do not appear to be intact, e.g., the 0.905 proportion of sequences from the FRESH cohort. [2] A large deletion was defined as being >20% of the median length of the genes in a given individual. [3] A missing 5’ end was defined as a deletion encompassing at least a portion of the ATG. [4] An alternative start codon, i.e., not an ATG, may be present.

| Supplementary Table 3. Data used to assess intact genes[1] | | | | | | | | | | |
|---|---|---|---|---|---|---|---|---|---|---|
| Trial | Gene | Time Pt. | N | Window | COB/EDDI (Days) | | | | | |
| | | | | | Min. | 1st Q | Median | Mean | 3rd Q | Max. |
| FRESH | Gag | 1 | 13 | NA | 2 | 5 | 9 | 11.85 | 19 | 24 |
| FRESH | Gag | 2 | 13 | 1 | 12 | 16 | 26 | 27.00 | 33 | 59 |
| FRESH | Gag | 3 | 10 | 2 | 19 | 34.25 | 55.5 | 47.10 | 57.75 | 61 |
| FRESH | Env | 1 | 13 | NA | 2 | 5 | 9 | 10.77 | 19 | 24 |
| FRESH | Env | 2 | 13 | 1 | 12 | 16 | 23 | 24.23 | 30 | 44 |
| FRESH | Env | 3 | 10 | 2 | 19 | 34.25 | 51.5 | 46.20 | 57.5 | 61 |
| RV217 | Gag | 1 | 29 | NA | 2 | 4 | 5 | 5.34 | 7 | 14 |
| RV217 | Gag | 2 | 29 | NA | 5 | 10 | 15 | 16.52 | 18 | 51 |
| RV217 | Gag | 3 | 29 | 1 | 11 | 15 | 24 | 29.21 | 45 | 61 |
| RV217 | Gag | 4 | 16 | 2 | 15 | 19 | 27 | 30.25 | 39.25 | 61 |
| RV217 | Gag | 5 | 7 | NA | 26 | 30 | 34 | 34.86 | 39 | 46 |
| RV217 | Env | 1 | 29 | NA | 2 | 4 | 5 | 5.72 | 7 | 14 |
| RV217 | Env | 2 | 29 | 1 | 5 | 11 | 15 | 16.90 | 18 | 51 |
| RV217 | Env | 3 | 29 | 2 | 11 | 17 | 24 | 29.41 | 45 | 61 |
| RV217 | Env | 4 | 16 | NA | 15 | 23 | 31 | 32.19 | 39.25 | 61 |
| RV217 | Env | 5 | 5 | NA | 26 | 29 | 35 | 35.80 | 43 | 46 |
| V703 | Gag | 1 | 32 | 1 | 7 | 18 | 31 | 29.72 | 35.25 | 89 |
| V703 | Gag | 2 | 28 | 2 | 26 | 32.75 | 46 | 49.18 | 57 | 109 |
| V703 | Env | 1 | 32 | 1 | 7 | 18 | 31 | 28.81 | 35.25 | 89 |
| V703 | Env | 2 | 28 | 2 | 26 | 32.75 | 46 | 47.04 | 55.5 | 108 |
| V704 | Gag | 1 | 47 | 1 | 8 | 18 | 31 | 32.62 | 37.5 | 135 |
| V704 | Gag | 2 | 40 | 2 | 13 | 33.5 | 38.5 | 46.35 | 47 | 187 |
| V704 | Gag | 3 | 13 | NA | 20 | 46 | 56 | 58.54 | 60 | 151 |
| V704 | Env | 1 | 47 | 1 | 8 | 18 | 28 | 32.02 | 37 | 135 |
| V704 | Env | 2 | 41 | 2 | 13 | 34 | 38 | 46.10 | 46 | 187 |
| V704 | Env | 3 | 13 | NA | 20 | 46 | 56 | 58.54 | 60 | 151 |

[1] Samples in the blue font were included in analysis using the 1st time window, and red font if included in analysis using the 2nd time window. Samples in black font were not used in these analyses. NA = Not applicable, i.e., outside of windows 1 or 2.

### Supplementary Table 4

**Supplementary Table 2A. Proportion of genes that were intact (Generalized Linear Model; GLM)**

| Comparison | Estimate | Std.Error | t.Value | p.Value |
|---|---|---|---|---|
| gag vs. env | 0.9405822 | 0.5129923 | 5.38391 | 1.20E-07 |
| B vs. C | 0.8396171 | 0.6310992 | 2.62301 | 9.30E-03 |
| Other* vs. B | 0.1350254 | 0.6555201 | -2.8332 | 5.30E-03 |
| Other* vs C | 0.4497069 | 0.6310992 | -0.3198 | 0.75 |

**Supplementary Table 2B. Proportion of genes that were intact (Wilcoxon tests)**

| | | Time Window 1[#] | | | Time Window 2 | | |
|---|---|---|---|---|---|---|---|
| **Comparison** | **Subset** | **Medians** | **P-Value** | **FDR Q** | **Medians** | **P-Value** | **FDR Q** |
| gag vs. env^ | Clade B | 98% vs. 93% | 2.90E-11 | 1.74E-10 | 98% vs. 91% | 4.70E-10 | 1.41E-09 |
| gag vs. env^ | Clade C | 92% vs 82% | 1.14E-13 | 2.05E-12 | 88% vs. 79% | 5.80E-11 | 2.61E-10 |
| gag vs. env^ | Other* | 97% vs. 83% | 2.90E-11 | 1.74E-10 | 98% vs. 90% | 1.90E-06 | 3.11E-06 |
| C vs. B | gag | 92% vs. 98% | 8.64E-07 | 1.56E-06 | 88% vs. 98% | 2.74E-10 | 9.86E-10 |
| B vs. Other* | gag | 98% vs. 97% | 0.006 | 7.20E-03 | 98% vs. 98% | 0.546 | 0.546 |
| C vs. Other* | gag | 92% vs. 97% | 0.0003 | 3.86E-04 | 88% vs. 98% | 1.51E-08 | 3.02E-08 |
| C vs. B | env | 82% vs. 93% | 7.11E-09 | 1.60E-08 | 79% vs. 91% | 3.50E-09 | 9.00E-09 |
| B vs. Other* | env | 93% vs. 83% | 2.19E-05 | 3.29E-05 | 91% vs. 90% | 0.163 | 0.173 |
| C vs. Other* | env | 82% vs. 83% | 0.141 | 0.159 | 79% vs. 90% | 5.08E-05 | 7.03E-05 |

**Supplementary Table 2C. Proportion of sequences that were hypermutated (Wilcoxon tests)**

| | | Time Window 1 | | | Time Window 2 | | |
|---|---|---|---|---|---|---|---|
| **Comparison** | **Subset** | **Medians** | **P-Value** | **FDR Q** | **Medians** | **P-Value** | **FDR Q** |
| gag vs. env | Clade B | 0% vs. 0.78% | 0.665 | 0.856 | 0.34% VS. 1.6% | 0.223 | 0.574 |
| gag vs. env | Clade C | 0.22% vs. 0.17% | 0.620 | 0.856 | 0.21% VS. 0% | 0.075 | 0.345 |
| gag vs. env | Other* | 0.65% vs. 0.64% | 0.124 | 0.372 | 0% VS. 0% | 0.295 | 0.589 |
| C vs. B | gag | 0.22% vs. 0% | 0.839 | 0.899 | 0.21% VS. 0.34% | 0.894 | 0.899 |
| B vs. Other* | gag | 0% vs. 0.65% | 0.096 | 0.345 | 0.34% VS. 0% | 0.866 | 0.899 |
| C vs. Other* | gag | 0.22% vs. 0.65% | 0.078 | 0.345 | 0.21% VS. 0% | 0.899 | 0.899 |
| C vs. B | env | 0.17% vs. 0.78% | 0.288 | 0.589 | 0% VS. 1.6% | 0.001 | 0.012 |
| B vs. Other* | env | 0.78% vs. 0.64% | 0.619 | 0.856 | 0% VS. 1.6% | 0.025 | 0.223 |
| C vs. Other* | env | 0.17% vs. 0.64% | 0.492 | 0.804 | 0% VS. 0% | 0.369 | 0.665 |

* "Other" refers to a combined dataset including all intersubtype recombinants and pure subtype sequences aside from subtype B and C. * Two time windows were examined, see Online Methods. ^ Normalized by gene length when comparing different gene regions. Red font indicates significant comparisons by both p-value <0.05 and FDR Q<0.2

**Table 5. Selection within Gag and Env at different stages of Acute and Early infection[1]**

| Protein | Stage | Number of Sites | α>β (negative) [2] | α<β (positive) [2] | Ratio (Neg/Pos.) |
|---|---|---|---|---|---|
| Gag | Overall | 238,779 | 2026 (0.85%) | 290 (0.12%) | 7.1 |
| | i | 70,338 | 342 (0.49%) | 52 (0.07%) | 7.0 |
| | ii | 92,550 | 867 (0.94%) | 103 (0.11%) | 8.5 |
| | iii | 54,296 | 579 (1.07%) | 76 (0.14%) | 7.6 |
| | iv | 21,595 | 238 (1.10%) | 59 (0.27%) | |
| Env | Overall | 481,100 | 1249 (0.26%) | 801 (0.17%) | 1.5 |
| | i | 158,480 | 276 (0.17%) | 137 (0.09%) | 1.9 |
| | ii | 186,780 | 497 (0.27%) | 288 (0.15%) | 1.8 |
| | iii | 101,880 | 346 (0.34%) | 286 (0.28%) | 1.2 |
| | iv | 33,960 | 130 (0.38%) | 90 (0.27%) | |

[1] Only lineages with at least 10 members were used in these calculations. [2] Sites with probabilities ≥0.9 (percent of total).

| **Supplementary Table 6. Acute and Early Infection Staging** | |
|---|---|
| Infection Stage definitions | Stage |
| Ascending viremia; 1-18 days from the COB or EDDI[1]; Fiebig I-III[2] | i |
| Rapidly descending viremia; 19-40 days from the COB or EDDI; Fiebig IV-V | ii |
| Slowly descending viremia; 41-61 days from the COB or EDDI; Fiebig V | iii |
| Entering steady state viremia; > 61 days from the COB or EDDI; Fiebig V | iv |

[1] COB = Center of bounds between the last negative and first positive HIV RNA test, used to date virus acquisitions for the RV217 and FRESH cohorts. EDDI = Estimated date of detectable infection assigned using COB and immune response data (Rossenkhan R, et al, mBio. 2025:e0188125). [2] Fiebig stages of infection (Fiebig EW, et al. AIDS. 2003;17(13):1871–9).

**Supplementary Table 7. Median gp120 Env and variable region lengths by subtype**

| Dataset: Subtype: | LANL B | RV217/ HVTN 704 B | LANL C | RV217/FRESH /HVTN 703 C | LANL CRF01AE | RV217 CRF01AE |
|---|---|---|---|---|---|---|
| V1 | 31 | 30 | 28 | 27 | 31 | 31 |
| V2 | 42 | 42 | 44 | 43 | 43 | 45 |
| V4 | 32 | 33 | 29 | 28 | 28 | 29 |
| V5 | 13 | 14 | 13 | 13 | 12 | 12 |
| V regions total | 118 | 119 | 114 | 111 | 114 | 117 |
| gp120 | 512 | 514 | 505 | 506 | 507 | 511 |

**Supplementary Fig. 1**

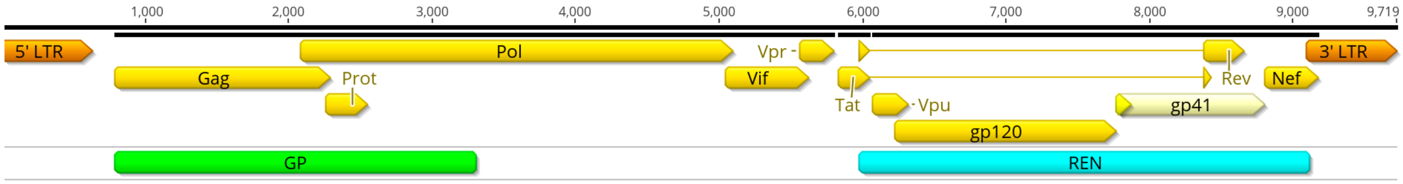

**S1 Fig. HIV-1 genome representation corresponding to the reference sequence HXB2 (GenBank accession number K03455.1) with amplicons used in this study shown below the open reading frame map.** The GP amplicon extends from HXB2 position 790 to 3297 and the REN amplicon from HXB2 position 5970 to 9012. Image generated in Geneious Prime version 2025.2.1 (Dotmatics).

#### Supplementary Fig. 2

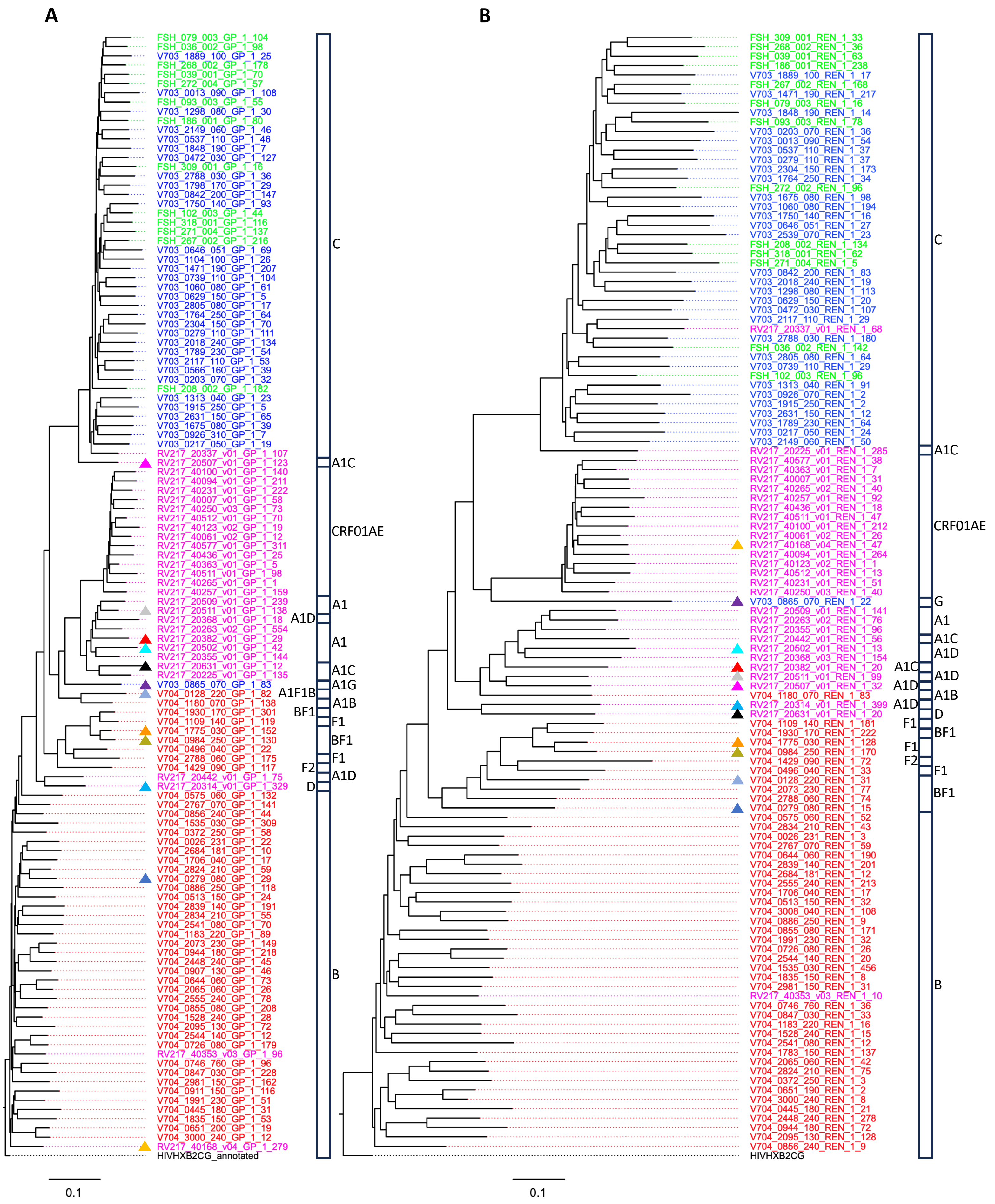

**Supplementary Figure 2. Phylogenetic analysis of representative HIV viral genome sequences used in this study.** The most common sequence detected in the GP (**A**) and REN (**B**) region from each participant were aligned and maximum likelihood trees generated. Sequence labels were colored by cohort (FRESH, green; HVTN 703, blue; RV217, magenta; HVTN 704, red), with the Subtype B HXB2 reference sequence in black. The identity of subtypes and intersubtype recombinant designations are indicated adjacent to the sectors within the vertical bars. Viruses with discordant subtypes in GP versus REN are indicated with colored triangles to the left of the sequence labels. The scales at the bottom of each tree indicate 10% sequence divergence.

**Supplementary Fig. 3**

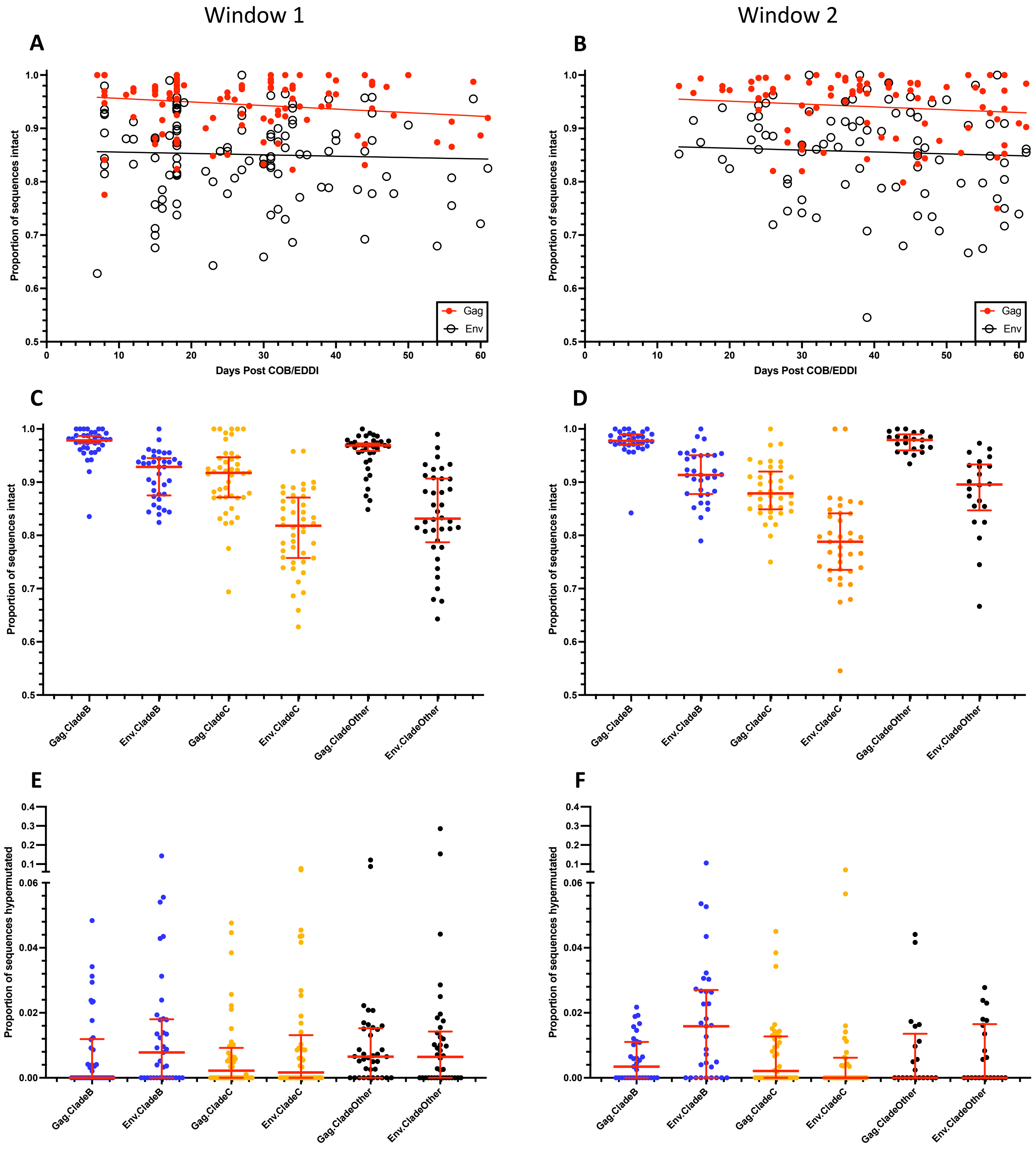

**Supplementary Fig 3. Intact and hypermutated *gag* and *env* genes as a function of time and viral subtype.** The proportion of intact *gag* and *env* genes (i.e., lacking putative inactivating mutations, S1 Table) as a function of days post COB (RV217 and FRESH cohorts) or EDDI (HVTN 703 and HVTN 704 cohorts) are shown for the first (**A**) and second (**B**) time windows. Red symbols refer to *gag* genes and open black symbols refer to *env* genes. The proportion of intact genes as a function of viral subtype are shown in panels (**C**) and (**D**) in windows 1 and 2, respectively. Panels (**E**) and (**F**) show proportion of *gag* and *env* genes that were hypermutated (25) in windows 1 and 2, respectively.

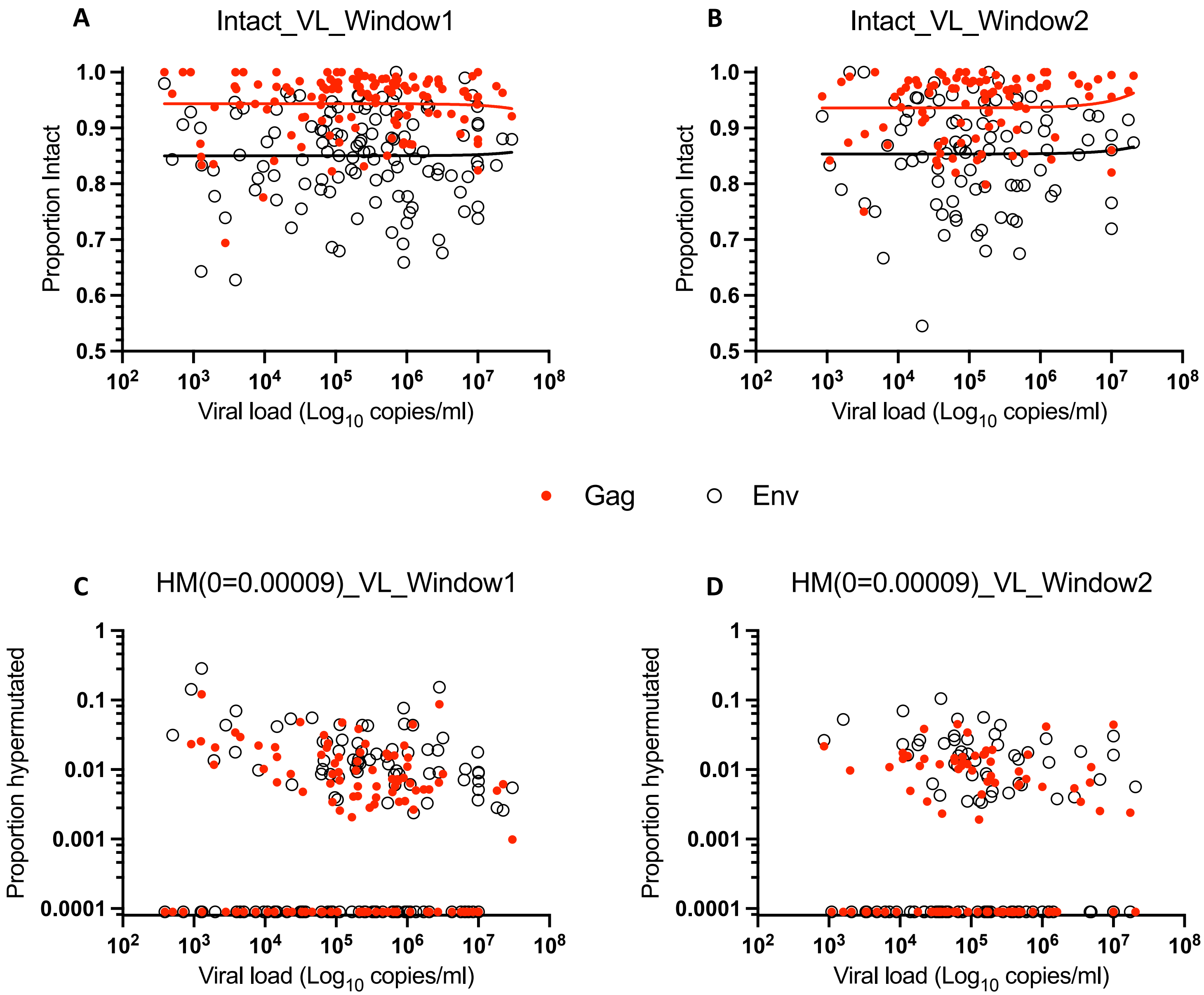

**Supplementary Fig. 4. Viral load correlated with intact genes and hypermutation.** Proportion of intact genes as a function of viral load in the first (**A**) and second (**B**) time windows. Proportion of sequences that were hypermutated as a function of plasma viral load for time windows 1 (**C**) and 2 (**D**). For illustration on a log plot, those datasets with no hypermutated sequences were set to a value of 0.00009. Gag gene values are shown with red dots and Env genes with open black circles.

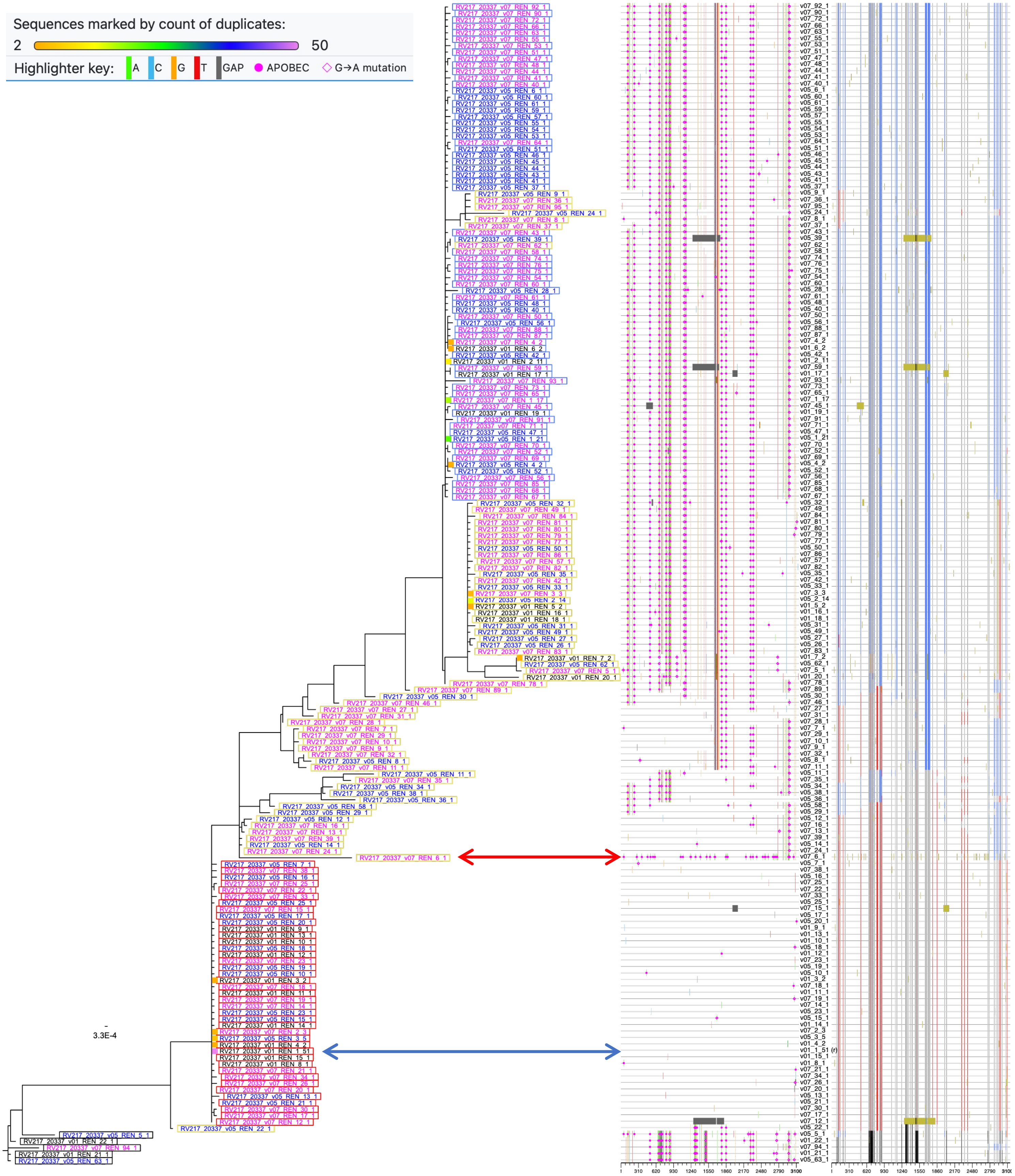

**Supplementary Fig. 5. Phylogeny-based lineage definition method.** Phylobook (Furlong, J.C., et al. *Biotechniques* **76**, 263–274 (2024) output from one RV217 participant. **Left panel:** Identical sequences were collapsed into one, with the number of collapsed sequences roughly indicated by a colored box to the left of the taxon name (see color key at top) and the precise number shown at the end of the taxon name. Taxon names were color coded based on visit (1st, black; 2nd, blue; 3rd, magenta). Taxon names were enclosed with colored boxes corresponding to different lineages: most common at the 1st time point, red; 2nd most common, blue; 3rd, black; recombinants, yellow). **Middle panel:** Highlighter plot showing differences relative to the most common sequence at the 1st timepoint (blue arrow). Dark gray areas show the position of deletions. The red arrow indicates a hypermutated sequence. **Right panel:** Match plot showing identities to each of the 3 identified lineages (gray if found in multiple lineages).

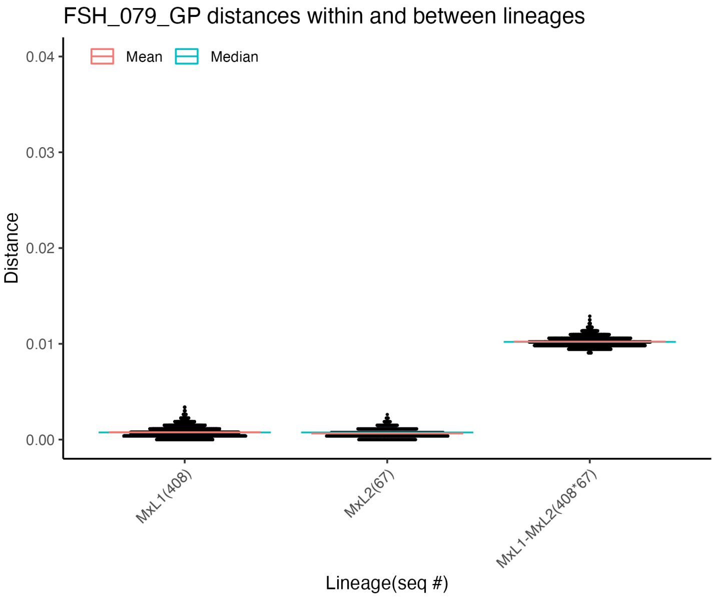

**Supplementary Figure 6. Maximum likelihood pairwise differences within participants infected with multiple lineages.** Pairwise differences were determined for all time points combined. Genetic distance distributions are shown for individual lineages as well as for inter-lineage comparisons. The numbers in parentheses below each distribution indicate the number of sequences within that lineage. In some cases (data points circled in red), deletions within variable regions resulted in lower pairwise distances. MxL(n) refers to the lineages observed within multilineage infections, numbered in order of abundance at the first time point sampled. "x" refers to the stage of infection, not specified here since all timepoints were pooled for these analyses.

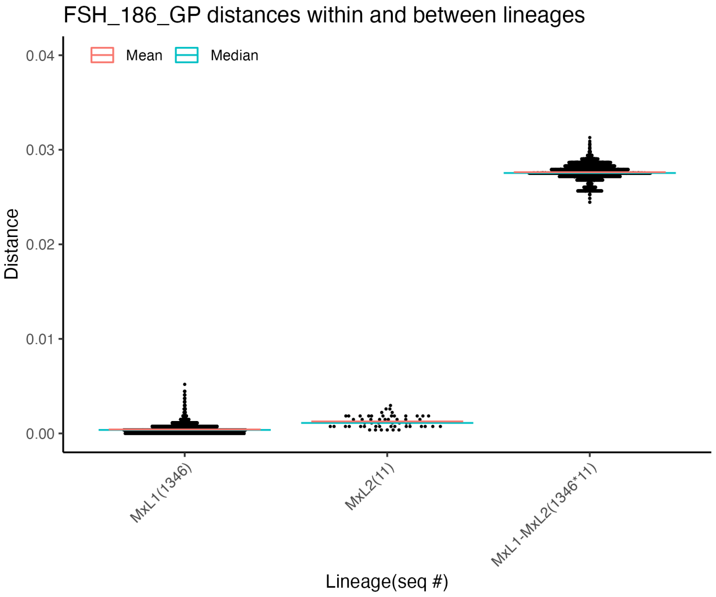

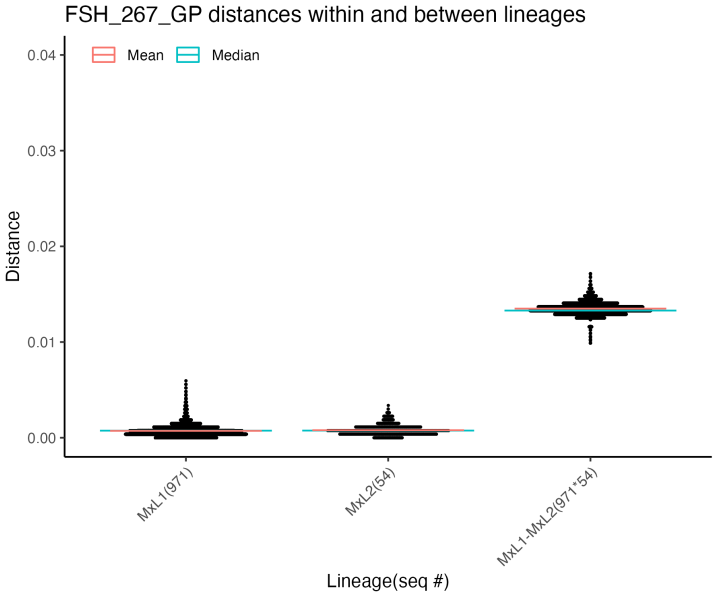

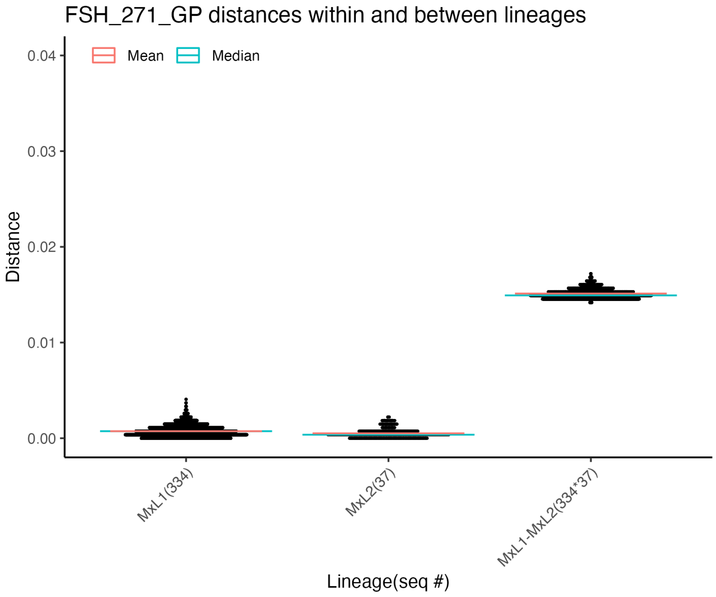

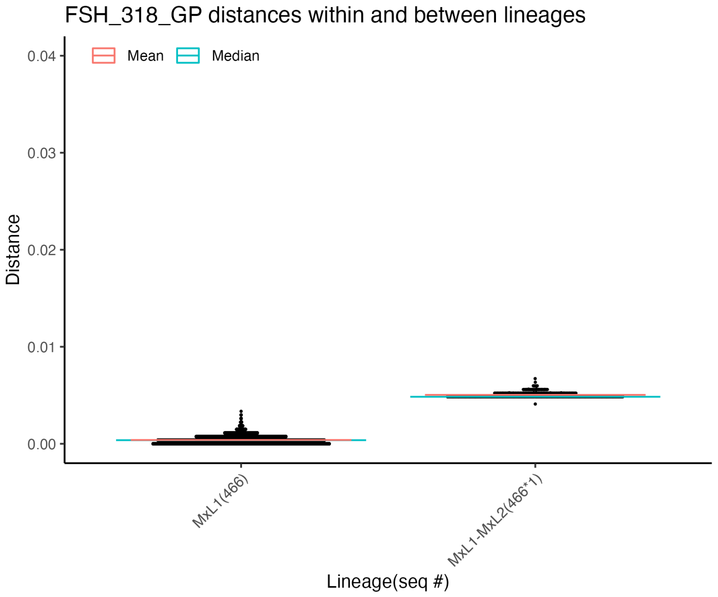

**S3.2 Fig.**

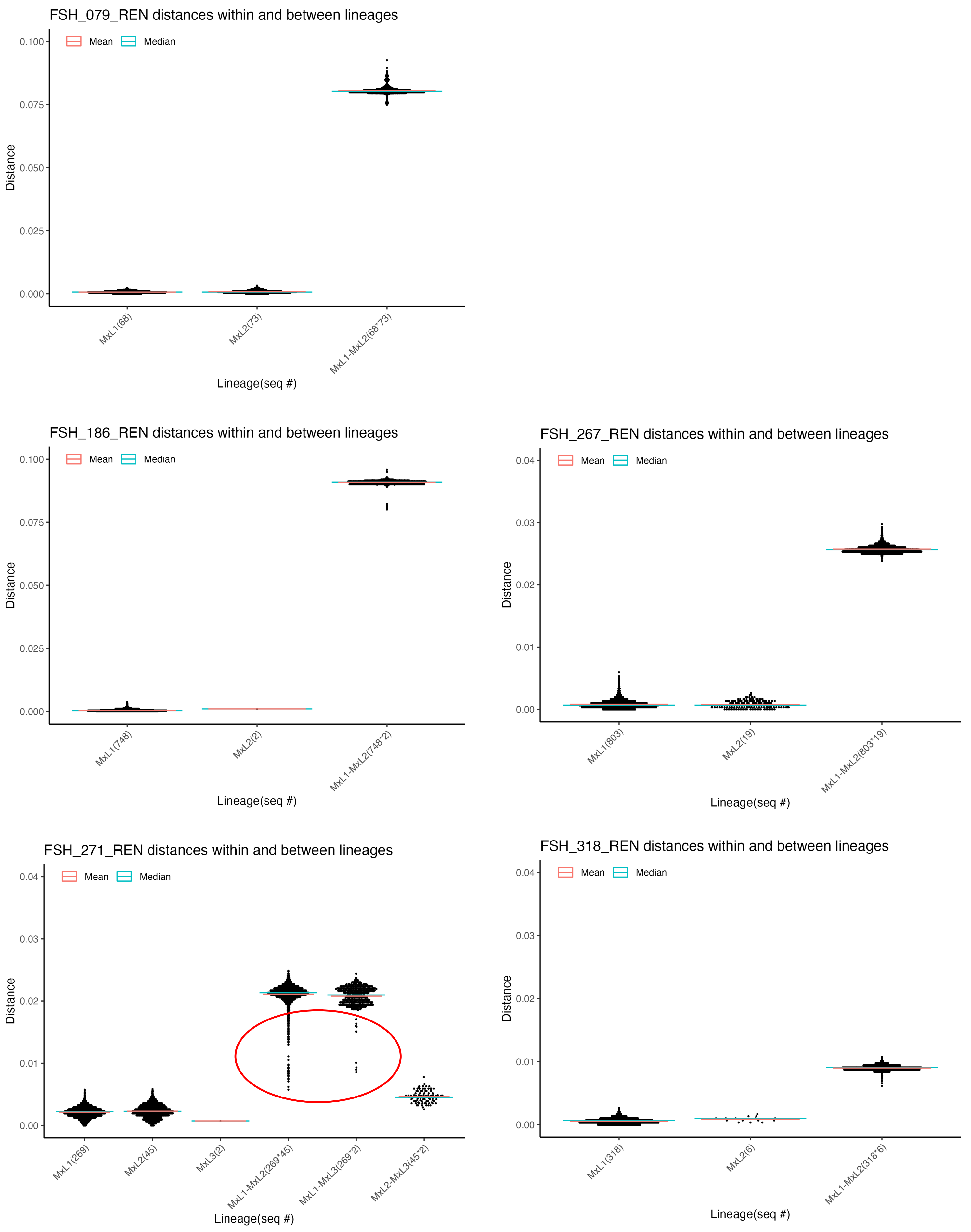
FSH_079_REN distances within and between lineages
Mean
Median
Distance
0.100
0.075
0.050
0.025
0.000
MxL1(68)
MxL2(73)
MxL1-MxL2(68*73)
Lineage(seq #)
FSH_186_REN distances within and between lineages
Mean
Median
Distance
0.100
0.075
0.050
0.025
0.000
MxL1(748)
MxL2(2)
MxL1-MxL2(748*2)
Lineage(seq #)
FSH_267_REN distances within and between lineages
Mean
Median
Distance
0.04
0.03
0.02
0.01
0.00
MxL1(803)
MxL2(19)
MxL1-MxL2(803*19)
Lineage(seq #)
FSH_271_REN distances within and between lineages
Mean
Median
Distance
0.04
0.03
0.02
0.01
0.00
MxL1(269)
MxL2(45)
MxL3(2)
MxL1-MxL2(269*45)
MxL1-MxL3(269*2)
MxL2-MxL3(45*2)
Lineage(seq #)
FSH_318_REN distances within and between lineages
Mean
Median
Distance
0.04
0.03
0.02
0.01
0.00
MxL1(318)
MxL2(6)
MxL1-MxL2(318*6)
Lineage(seq #)

**S6.3 Fig.**

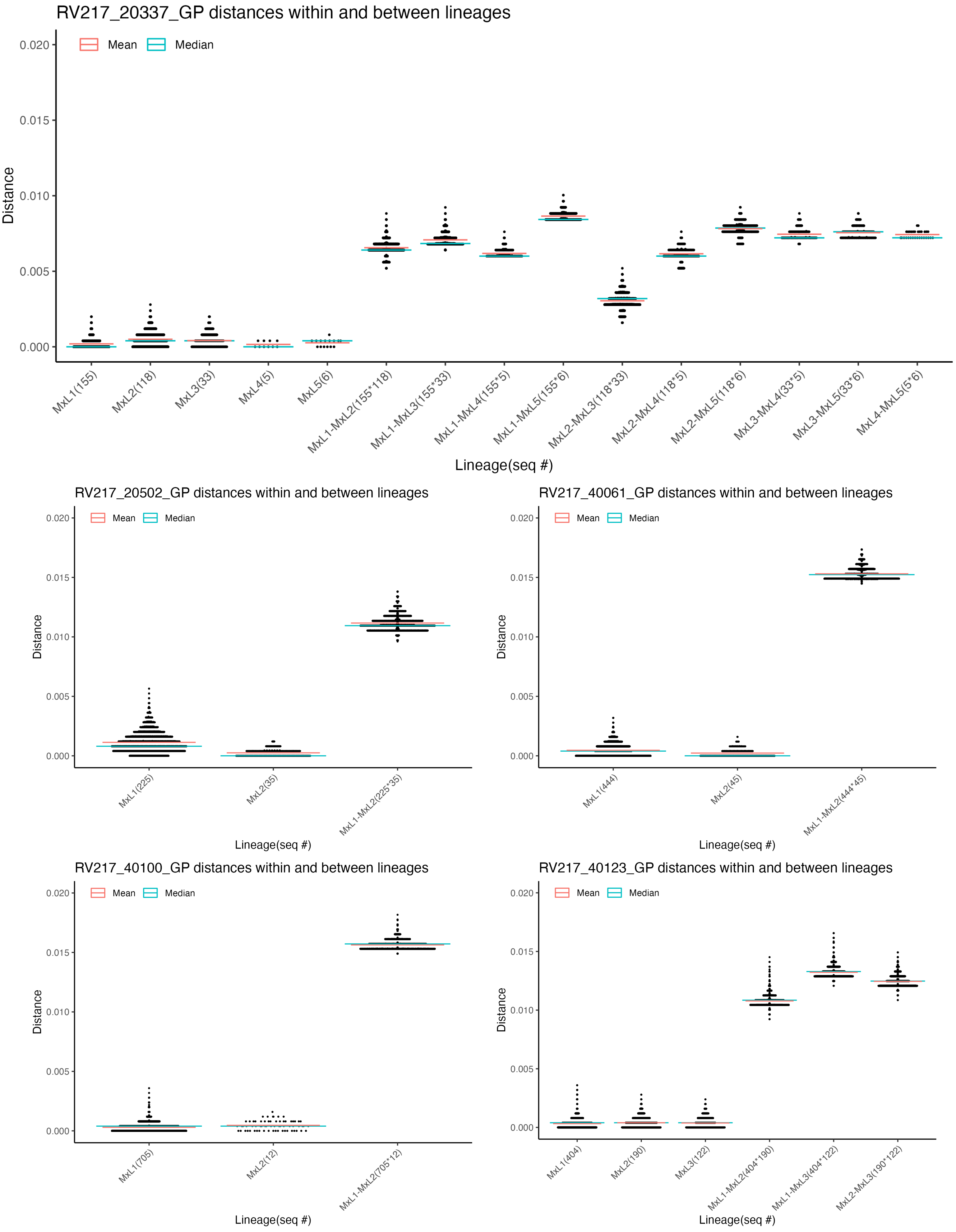
RV217_20337_GP distances within and between lineages
Mean
Median
Distance
0.020
0.015
0.010
0.005
0.000
MxL1(155)
MxL2(118)
MxL3(33)
MxL4(5)
MxL5(6)
MxL1-MxL2(155*118)
MxL1-MxL3(155*33)
MxL1-MxL4(155*5)
MxL1-MxL5(155*6)
MxL2-MxL3(118*33)
MxL2-MxL4(118*5)
MxL2-MxL5(118*6)
MxL3-MxL4(33*5)
MxL3-MxL5(33*6)
MxL4-MxL5(5*6)
Lineage(seq #)
RV217_20502_GP distances within and between lineages
Mean
Median
Distance
0.020
0.015
0.010
0.005
0.000
MxL1(225)
MxL2(35)
MxL1-MxL2(225*35)
Lineage(seq #)
RV217_40061_GP distances within and between lineages
Mean
Median
Distance
0.020
0.015
0.010
0.005
0.000
MxL1(444)
MxL2(45)
MxL1-MxL2(444*45)
Lineage(seq #)
RV217_40100_GP distances within and between lineages
Mean
Median
Distance
0.020
0.015
0.010
0.005
0.000
MxL1(705)
MxL2(12)
MxL1-MxL2(705*12)
Lineage(seq #)
RV217_40123_GP distances within and between lineages
Mean
Median
Distance
0.020
0.015
0.010
0.005
0.000
MxL1(404)
MxL2(190)
MxL3(122)
MxL1-MxL2(404*190)
MxL1-MxL3(404*122)
MxL2-MxL3(190*122)
Lineage(seq #)

**S6.4 Fig.**

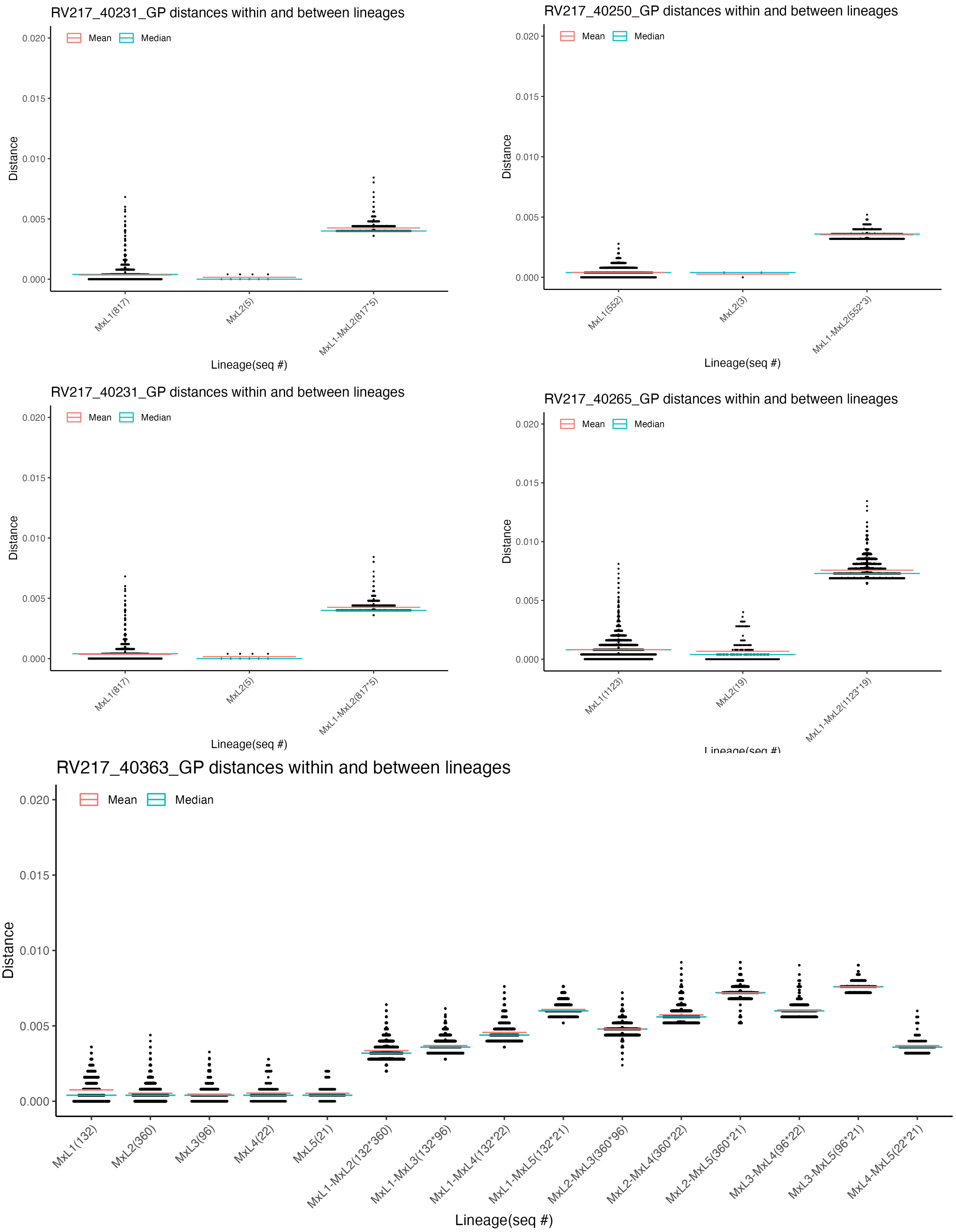
RV217_40231_GP distances within and between lineages
Mean
Median
Distance
0.020
0.015
0.010
0.005
0.000
MxL1(817)
MxL2(5)
MxL1-MxL2(817*5)
Lineage(seq #)
RV217_40250_GP distances within and between lineages
Mean
Median
Distance
0.020
0.015
0.010
0.005
0.000
MxL1(552)
MxL2(3)
MxL1-MxL2(552*3)
Lineage(seq #)
RV217_40231_GP distances within and between lineages
Mean
Median
Distance
0.020
0.015
0.010
0.005
0.000
MxL1(817)
MxL2(5)
MxL1-MxL2(817*5)
Lineage(seq #)
RV217_40265_GP distances within and between lineages
Mean
Median
Distance
0.020
0.015
0.010
0.005
0.000
MxL1(1123)
MxL2(19)
MxL1-MxL2(1123*19)
RV217_40363_GP distances within and between lineages
Mean
Median
Distance
0.020
0.015
0.010
0.005
0.000
MxL1(132)
MxL2(360)
MxL3(96)
MxL4(22)
MxL5(21)
MxL1-MxL2(132*360)
MxL1-MxL3(132*96)
MxL1-MxL4(132*22)
MxL1-MxL5(132*21)
MxL2-MxL3(360*96)
MxL2-MxL4(360*22)
MxL2-MxL5(360*21)
MxL3-MxL4(96*22)
MxL3-MxL5(96*21)
MxL4-MxL5(22*21)
Lineage(seq #)

**S6.5 Fig.**

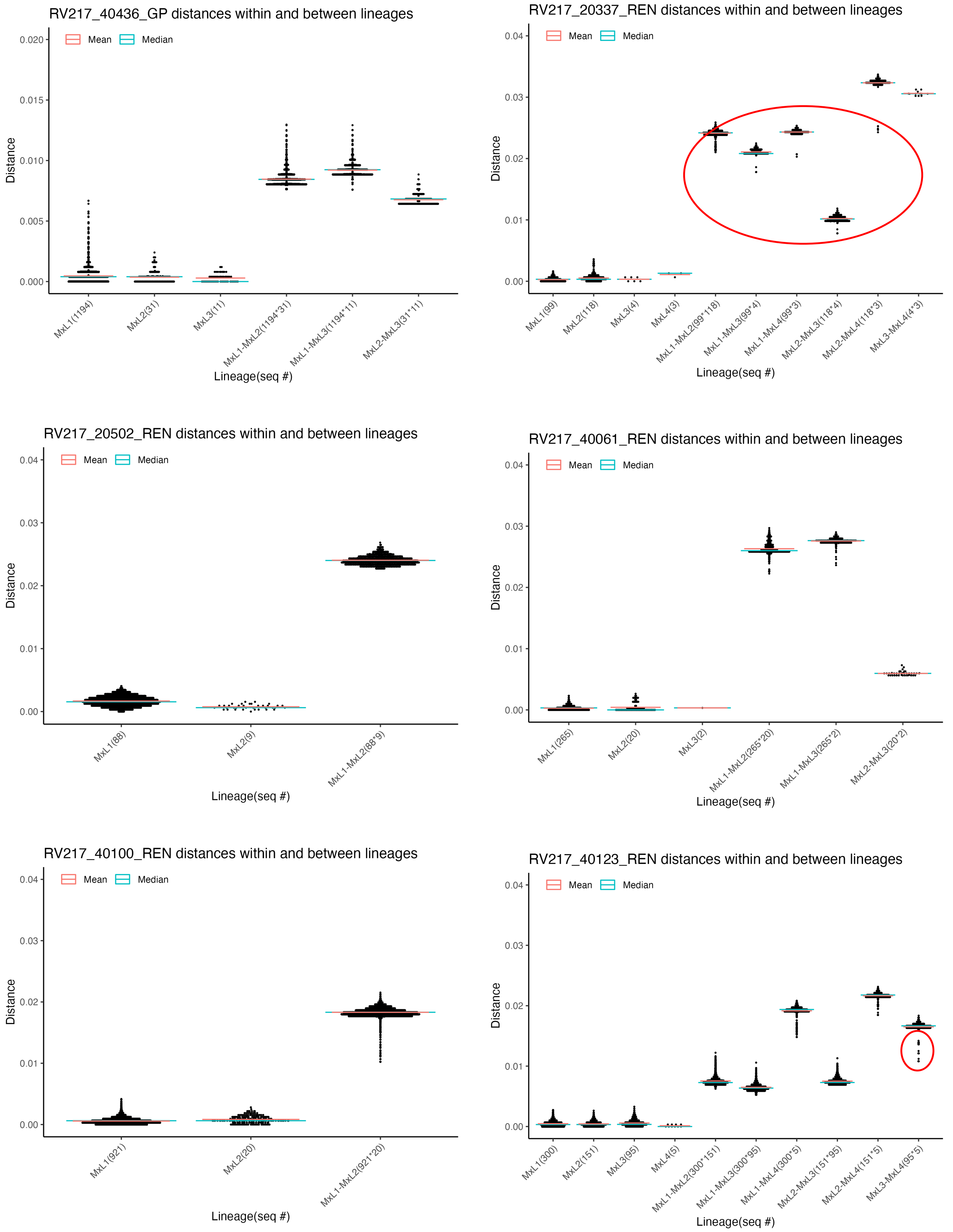
RV217_40436_GP distances within and between lineages
Mean
Median
Distance
0.020
0.015
0.010
0.005
0.000
MxL1(1194)
MxL2(31)
MxL3(11)
MxL1-MxL2(1194*31)
MxL1-MxL3(1194*11)
MxL2-MxL3(31*11)
Lineage(seq #)
RV217_20337_REN distances within and between lineages
Mean
Median
Distance
0.04
0.03
0.02
0.01
0.00
MxL1(99)
MxL2(118)
MxL3(4)
MxL4(3)
MxL1-MxL2(99*118)
MxL1-MxL3(99*4)
MxL1-MxL4(99*3)
MxL2-MxL3(118*4)
MxL2-MxL4(118*3)
MxL3-MxL4(4*3)
Lineage(seq #)
RV217_20502_REN distances within and between lineages
Mean
Median
Distance
0.04
0.03
0.02
0.01
0.00
MxL1(88)
MxL2(9)
MxL1-MxL2(88*9)
Lineage(seq #)
RV217_40061_REN distances within and between lineages
Mean
Median
Distance
0.04
0.03
0.02
0.01
0.00
MxL1(265)
MxL2(20)
MxL3(2)
MxL1-MxL2(265*20)
MxL1-MxL3(265*2)
MxL2-MxL3(20*2)
Lineage(seq #)
RV217_40100_REN distances within and between lineages
Mean
Median
Distance
0.04
0.03
0.02
0.01
0.00
MxL1(921)
MxL2(20)
MxL1-MxL2(921*20)
Lineage(seq #)
RV217_40123_REN distances within and between lineages
Mean
Median
Distance
0.04
0.03
0.02
0.01
0.00
MxL1(300)
MxL2(151)
MxL3(95)
MxL4(5)
MxL1-MxL2(300*151)
MxL1-MxL3(300*95)
MxL1-MxL4(300*5)
MxL2-MxL3(151*95)
MxL2-MxL4(151*5)
MxL3-MxL4(95*5)
Lineage(seq #)

**S3.6 Fig.**

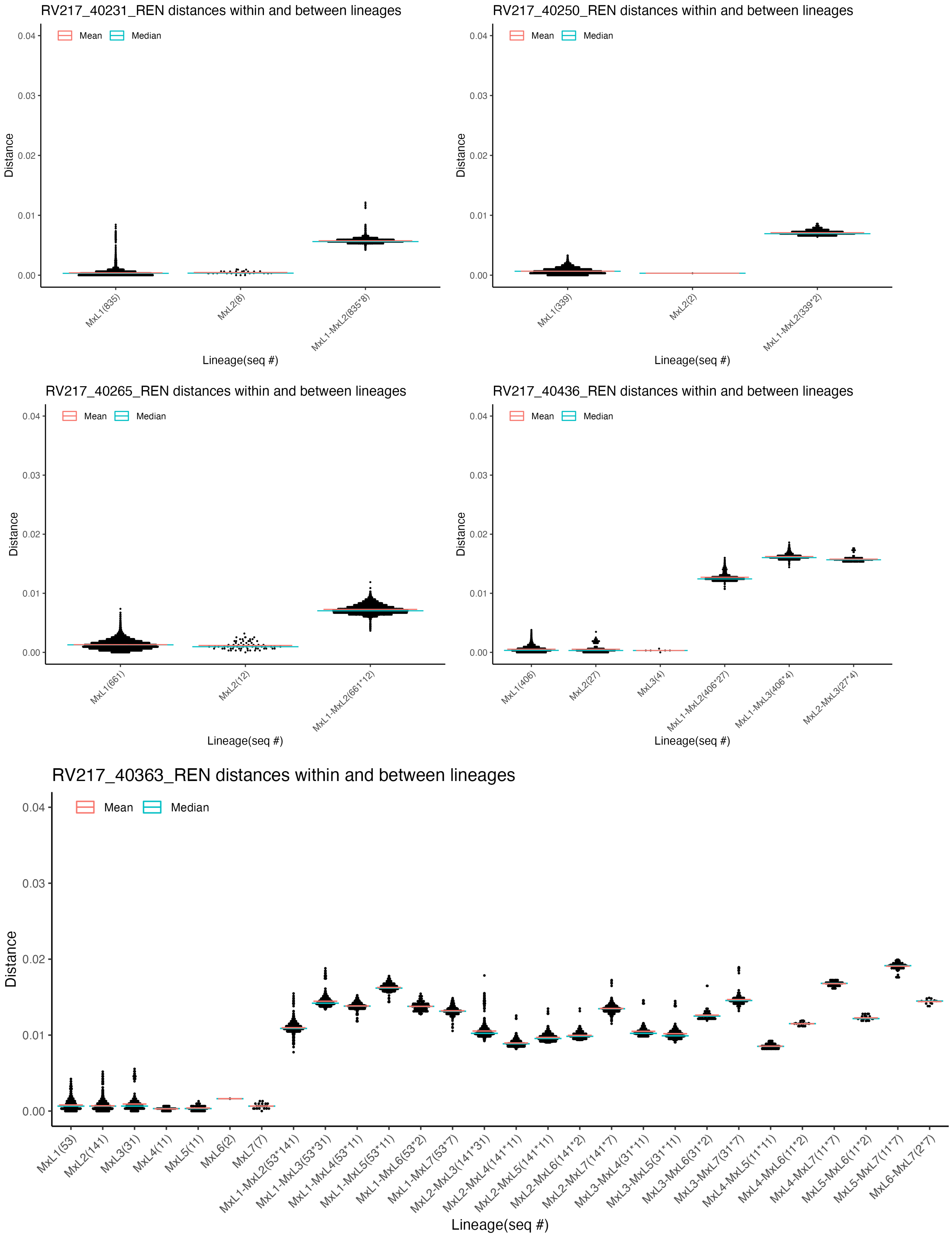
RV217_40231_REN distances within and between lineages
Mean
Median
Distance
0.04
0.03
0.02
0.01
0.00
MxL1(835)
MxL2(8)
MxL1-MxL2(835*8)
Lineage(seq #)
RV217_40250_REN distances within and between lineages
Mean
Median
Distance
0.04
0.03
0.02
0.01
0.00
MxL1(339)
MxL2(2)
MxL1-MxL2(339*2)
Lineage(seq #)
RV217_40265_REN distances within and between lineages
Mean
Median
Distance
0.04
0.03
0.02
0.01
0.00
MxL1(661)
MxL2(12)
MxL1-MxL2(661*12)
Lineage(seq #)
RV217_40436_REN distances within and between lineages
Mean
Median
Distance
0.04
0.03
0.02
0.01
0.00
MxL1(406)
MxL2(27)
MxL3(4)
MxL1-MxL2(406*27)
MxL1-MxL3(406*4)
MxL2-MxL3(27*4)
Lineage(seq #)
RV217_40363_REN distances within and between lineages
Mean
Median
Distance
0.04
0.03
0.02
0.01
0.00
MxL1(53)
MxL2(141)
MxL3(31)
MxL4(11)
MxL5(11)
MxL6(2)
MxL7(7)
MxL1-MxL2(53*141)
MxL1-MxL3(53*31)
MxL1-MxL4(53*11)
MxL1-MxL5(53*11)
MxL1-MxL6(53*2)
MxL1-MxL7(53*7)
MxL2-MxL3(141*31)
MxL2-MxL4(141*11)
MxL2-MxL5(141*11)
MxL2-MxL6(141*2)
MxL2-MxL7(141*7)
MxL3-MxL4(31*11)
MxL3-MxL5(31*11)
MxL3-MxL6(31*2)
MxL3-MxL7(31*7)
MxL4-MxL5(11*11)
MxL4-MxL6(11*2)
MxL4-MxL7(11*7)
MxL5-MxL6(11*2)
MxL5-MxL7(11*7)
MxL6-MxL7(2*7)
Lineage(seq #)

**S3.7 Fig.**

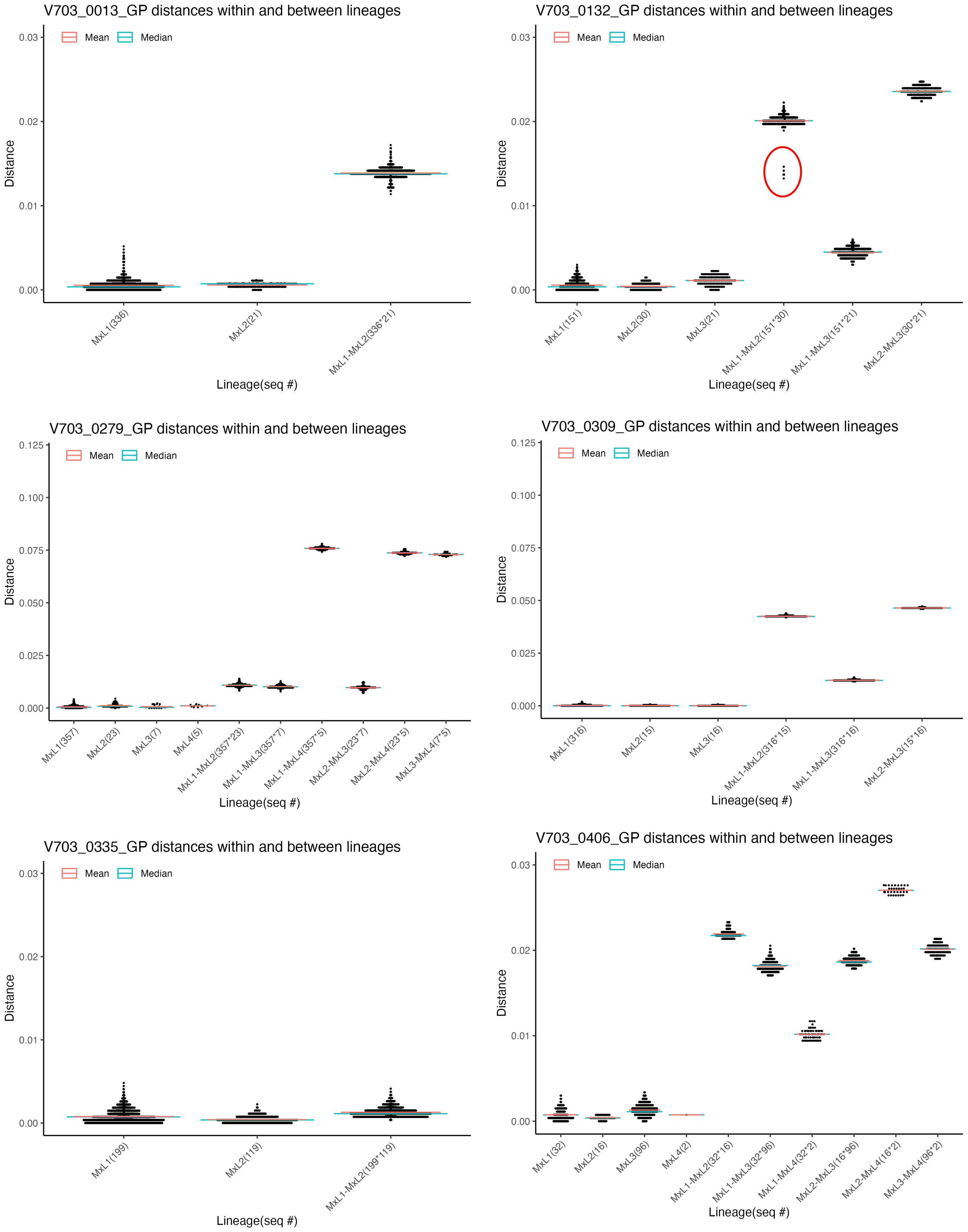
V703_0013_GP distances within and between lineages
Mean
Median
Distance
Lineage(seq #)
MxL1(336)
MxL2(21)
MxL1-MxL2(336*21)
V703_0132_GP distances within and between lineages
MxL1(151)
MxL2(30)
MxL3(21)
MxL1-MxL2(151*30)
MxL1-MxL3(151*21)
MxL2-MxL3(30*21)
V703_0279_GP distances within and between lineages
MxL1(357)
MxL2(23)
MxL3(7)
MxL4(5)
MxL1-MxL2(357*23)
MxL1-MxL3(357*7)
MxL1-MxL4(357*5)
MxL2-MxL3(23*7)
MxL2-MxL4(23*5)
MxL3-MxL4(7*5)
V703_0309_GP distances within and between lineages
MxL1(316)
MxL2(15)
MxL3(16)
MxL1-MxL2(316*15)
MxL1-MxL3(316*16)
MxL2-MxL3(15*16)
V703_0335_GP distances within and between lineages
MxL1(199)
MxL2(119)
MxL1-MxL2(199*119)
V703_0406_GP distances within and between lineages
MxL1(32)
MxL2(16)
MxL3(96)
MxL4(2)
MxL1-MxL2(32*16)
MxL1-MxL3(32*96)
MxL1-MxL4(32*2)
MxL2-MxL3(16*96)
MxL2-MxL4(16*2)
MxL3-MxL4(96*2)

**S3.8 Fig.**

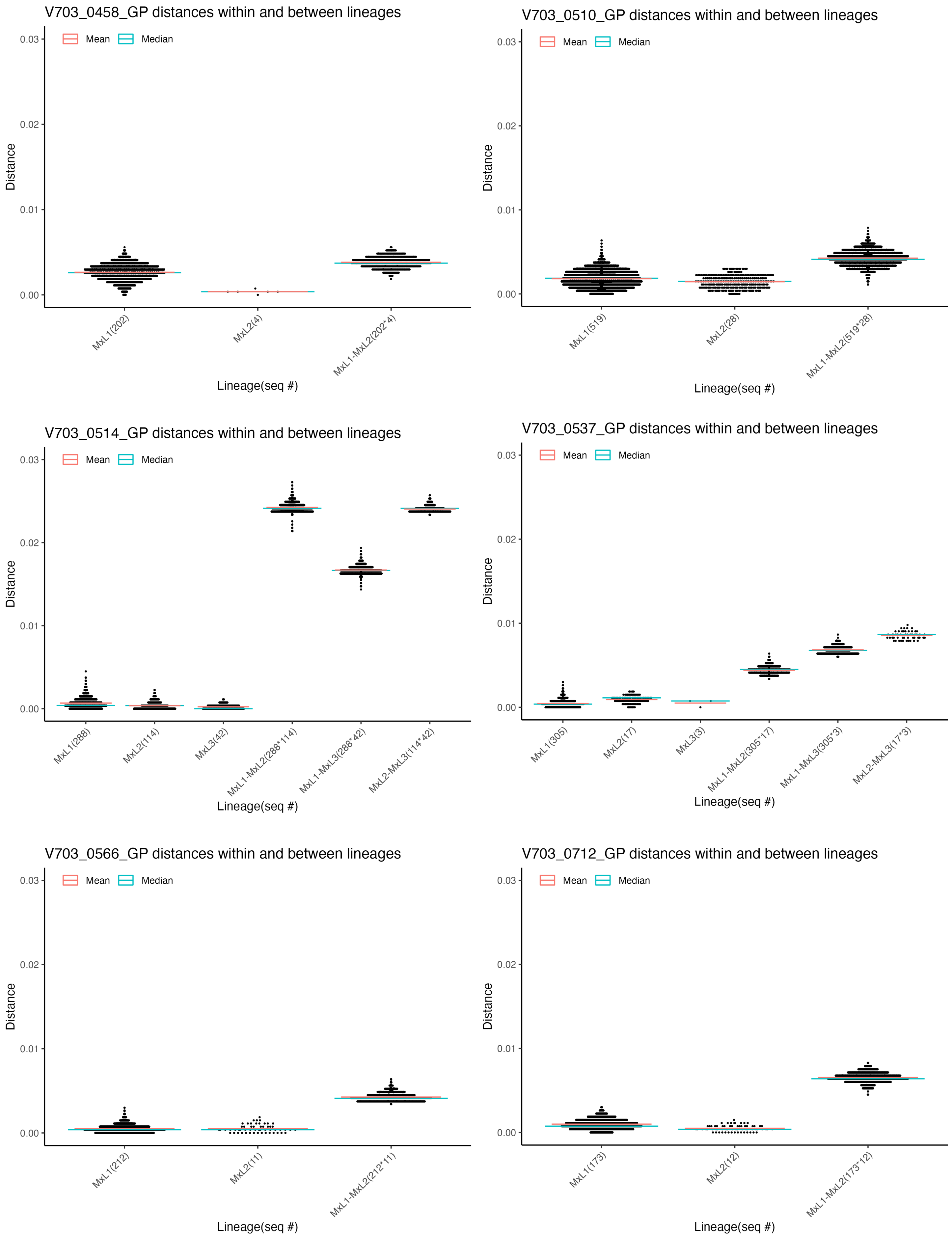
V703_0458_GP distances within and between lineages
Mean
Median
Distance
0.03
0.02
0.01
0.00
MxL1(202)
MxL2(4)
MxL1-MxL2(202*4)
Lineage(seq #)
V703_0510_GP distances within and between lineages
Mean
Median
Distance
0.03
0.02
0.01
0.00
MxL1(519)
MxL2(28)
MxL1-MxL2(519*28)
Lineage(seq #)
V703_0514_GP distances within and between lineages
Mean
Median
Distance
0.03
0.02
0.01
0.00
MxL1(288)
MxL2(114)
MxL3(42)
MxL1-MxL2(288*114)
MxL1-MxL3(288*42)
MxL2-MxL3(114*42)
Lineage(seq #)
V703_0537_GP distances within and between lineages
Mean
Median
Distance
0.03
0.02
0.01
0.00
MxL1(305)
MxL2(17)
MxL3(3)
MxL1-MxL2(305*17)
MxL1-MxL3(305*3)
MxL2-MxL3(17*3)
Lineage(seq #)
V703_0566_GP distances within and between lineages
Mean
Median
Distance
0.03
0.02
0.01
0.00
MxL1(212)
MxL2(11)
MxL1-MxL2(212*11)
Lineage(seq #)
V703_0712_GP distances within and between lineages
Mean
Median
Distance
0.03
0.02
0.01
0.00
MxL1(173)
MxL2(12)
MxL1-MxL2(173*12)
Lineage(seq #)

**S3.9 Fig.**

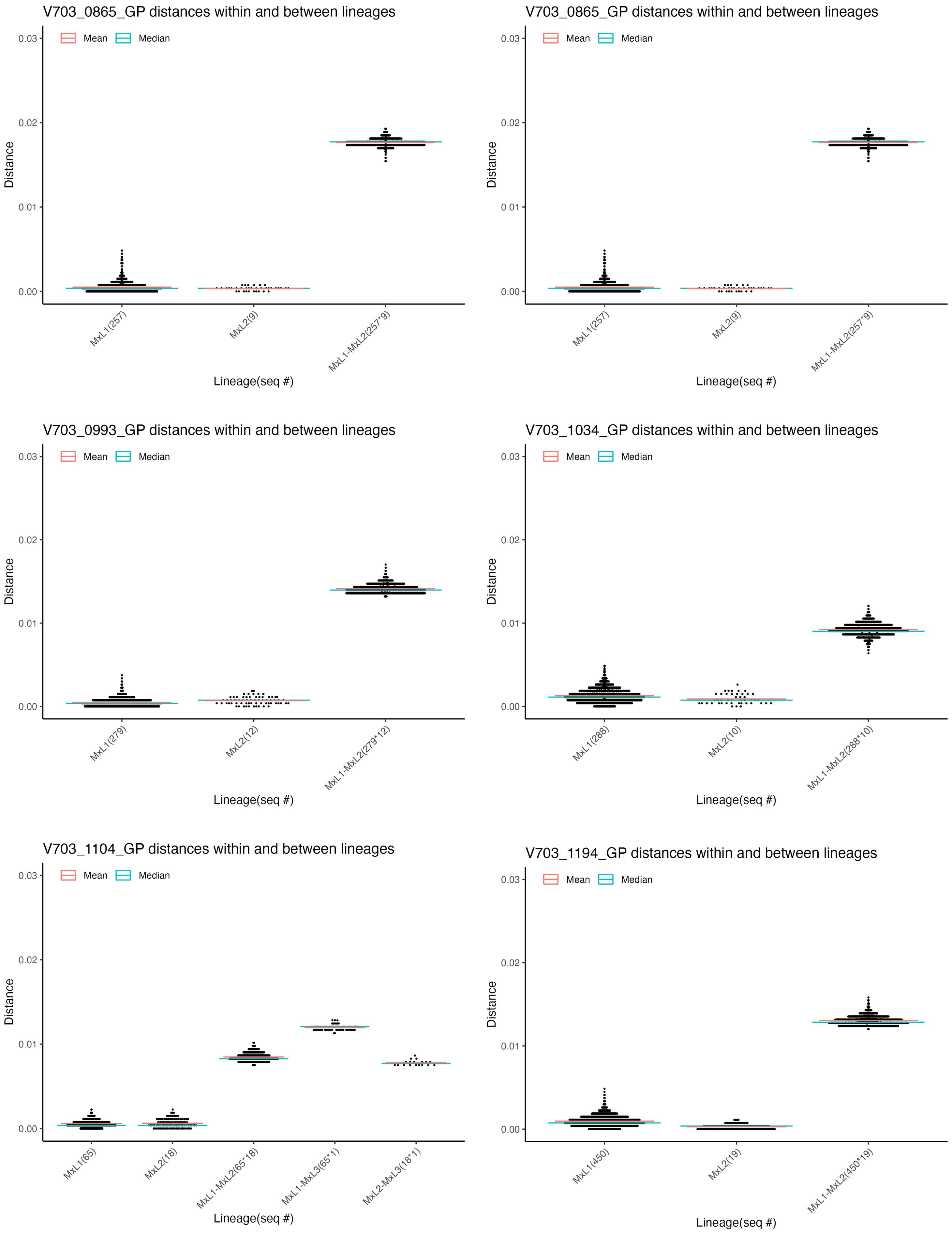
V703_0865_GP distances within and between lineages
Mean
Median
Distance
0.03
0.02
0.01
0.00
MxL1(257)
MxL2(9)
MxL1-MxL2(257*9)
Lineage(seq #)
V703_0865_GP distances within and between lineages
Mean
Median
Distance
MxL1(257)
MxL2(9)
MxL1-MxL2(257*9)
Lineage(seq #)
V703_0993_GP distances within and between lineages
Mean
Median
Distance
MxL1(279)
MxL2(12)
MxL1-MxL2(279*12)
Lineage(seq #)
V703_1034_GP distances within and between lineages
Mean
Median
Distance
MxL1(288)
MxL2(10)
MxL1-MxL2(288*10)
Lineage(seq #)
V703_1104_GP distances within and between lineages
Mean
Median
Distance
MxL1(65)
MxL2(18)
MxL1-MxL2(65*18)
MxL1-MxL3(65*1)
MxL2-MxL3(18*1)
Lineage(seq #)
V703_1194_GP distances within and between lineages
Mean
Median
Distance
MxL1(450)
MxL2(19)
MxL1-MxL2(450*19)
Lineage(seq #)

**S3.10 Fig.**

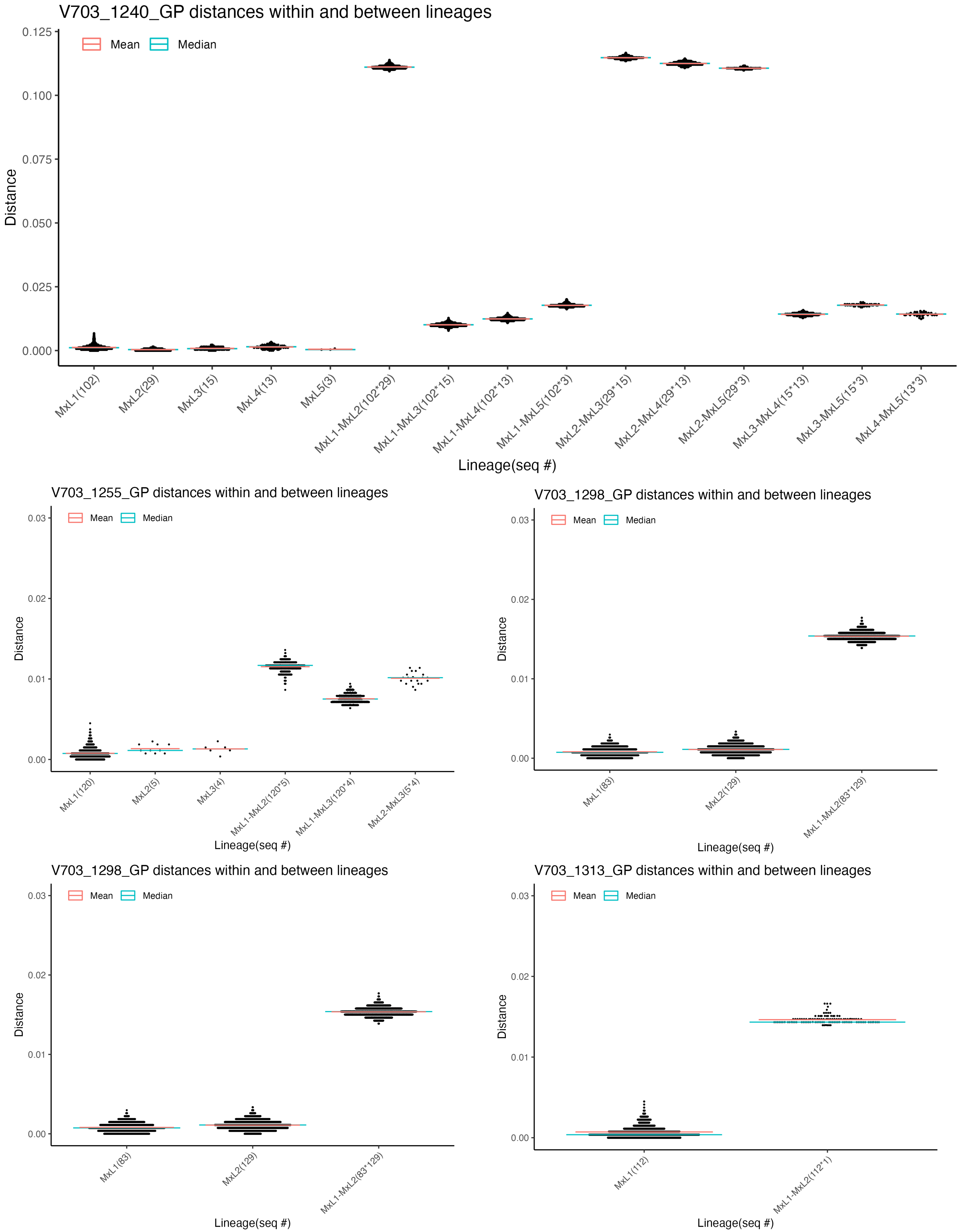
V703_1240_GP distances within and between lineages
Mean
Median
Distance
0.125
0.100
0.075
0.050
0.025
0.000
MxL1(102)
MxL2(29)
MxL3(15)
MxL4(13)
MxL5(3)
MxL1-MxL2(102*29)
MxL1-MxL3(102*15)
MxL1-MxL4(102*13)
MxL1-MxL5(102*3)
MxL2-MxL3(29*15)
MxL2-MxL4(29*13)
MxL2-MxL5(29*3)
MxL3-MxL4(15*13)
MxL3-MxL5(15*3)
MxL4-MxL5(13*3)
Lineage(seq #)
V703_1255_GP distances within and between lineages
Mean
Median
Distance
0.03
0.02
0.01
0.00
MxL1(120)
MxL2(5)
MxL3(4)
MxL1-MxL2(120*5)
MxL1-MxL3(120*4)
MxL2-MxL3(5*4)
Lineage(seq #)
V703_1298_GP distances within and between lineages
Mean
Median
Distance
0.03
0.02
0.01
0.00
MxL1(83)
MxL2(129)
MxL1-MxL2(83*129)
Lineage(seq #)
V703_1298_GP distances within and between lineages
Mean
Median
Distance
0.03
0.02
0.01
0.00
MxL1(83)
MxL2(129)
MxL1-MxL2(83*129)
Lineage(seq #)
V703_1313_GP distances within and between lineages
Mean
Median
Distance
0.03
0.02
0.01
0.00
MxL1(112)
MxL1-MxL2(112*1)
Lineage(seq #)

**S3.11 Fig.**

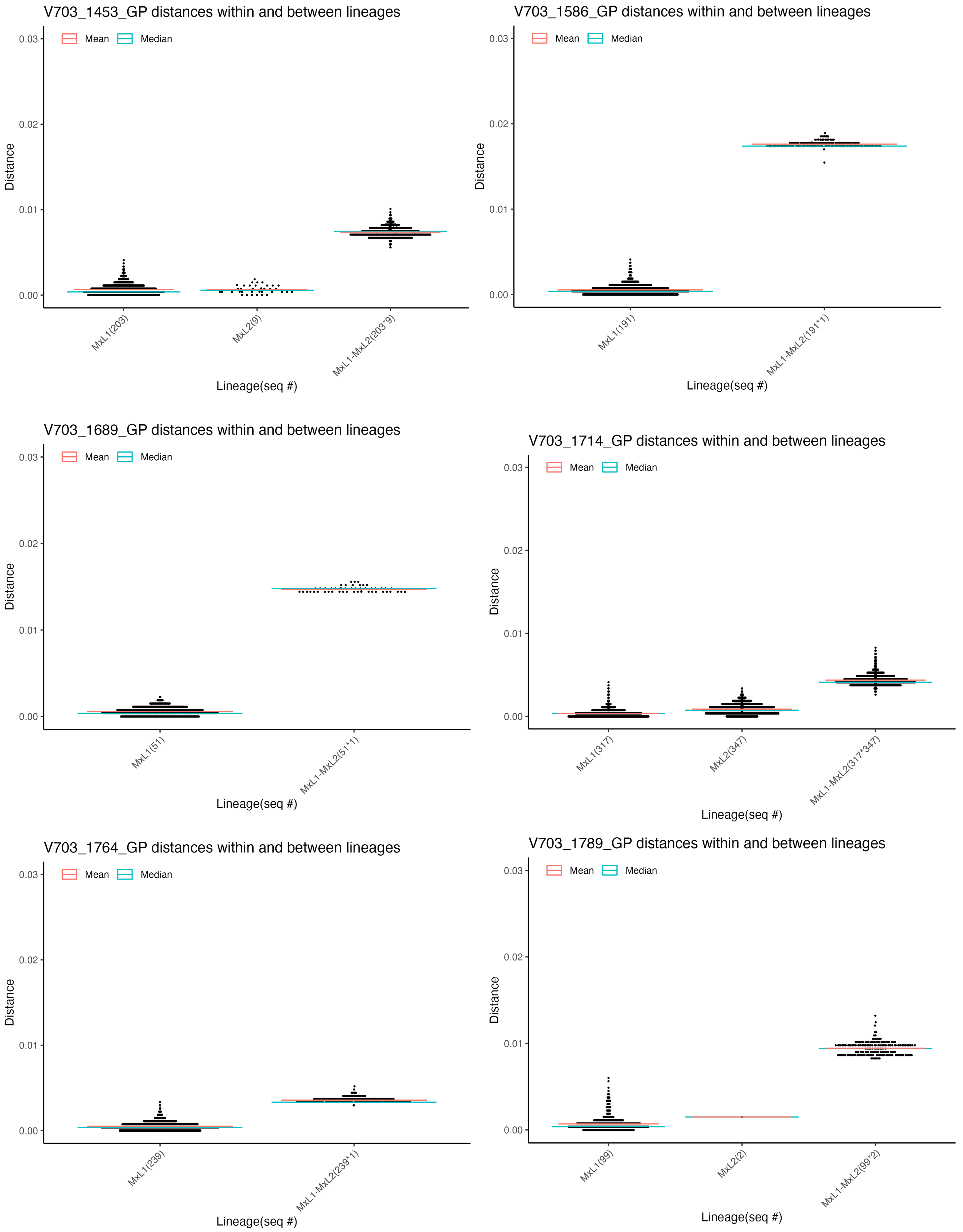
V703_1453_GP distances within and between lineages
Mean
Median
Distance
0.03
0.02
0.01
0.00
MxL1(203)
MxL2(9)
MxL1-MxL2(203*9)
Lineage(seq #)
V703_1586_GP distances within and between lineages
Mean
Median
Distance
0.03
0.02
0.01
0.00
MxL1(191)
MxL1-MxL2(191*1)
Lineage(seq #)
V703_1689_GP distances within and between lineages
Mean
Median
Distance
0.03
0.02
0.01
0.00
MxL1(51)
MxL1-MxL2(51*1)
Lineage(seq #)
V703_1714_GP distances within and between lineages
Mean
Median
Distance
0.03
0.02
0.01
0.00
MxL1(317)
MxL2(347)
MxL1-MxL2(317*347)
Lineage(seq #)
V703_1764_GP distances within and between lineages
Mean
Median
Distance
0.03
0.02
0.01
0.00
MxL1(239)
MxL1-MxL2(239*1)
Lineage(seq #)
V703_1789_GP distances within and between lineages
Mean
Median
Distance
0.03
0.02
0.01
0.00
MxL1(99)
MxL2(2)
MxL1-MxL2(99*2)
Lineage(seq #)

**S3.12 Fig.**

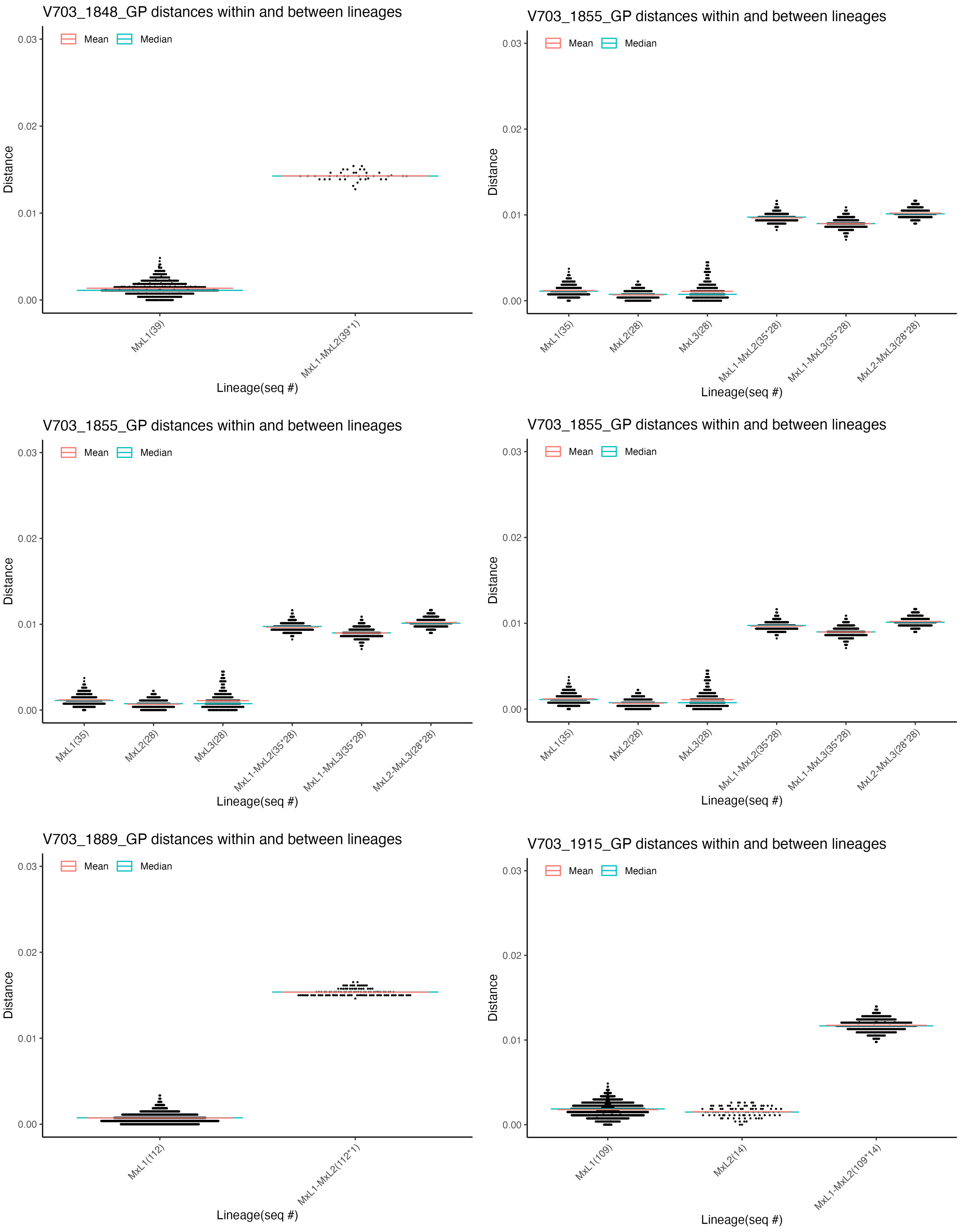
V703_1848_GP distances within and between lineages
Mean
Median
Distance
0.03
0.02
0.01
0.00
MxL1(39)
MxL1-MxL2(39*1)
Lineage(seq #)
V703_1855_GP distances within and between lineages
MxL1(35)
MxL2(28)
MxL3(28)
MxL1-MxL2(35*28)
MxL1-MxL3(35*28)
MxL2-MxL3(28*28)
V703_1889_GP distances within and between lineages
MxL1(112)
MxL1-MxL2(112*1)
V703_1915_GP distances within and between lineages
MxL1(109)
MxL2(14)
MxL1-MxL2(109*14)

**S3.13 Fig.**

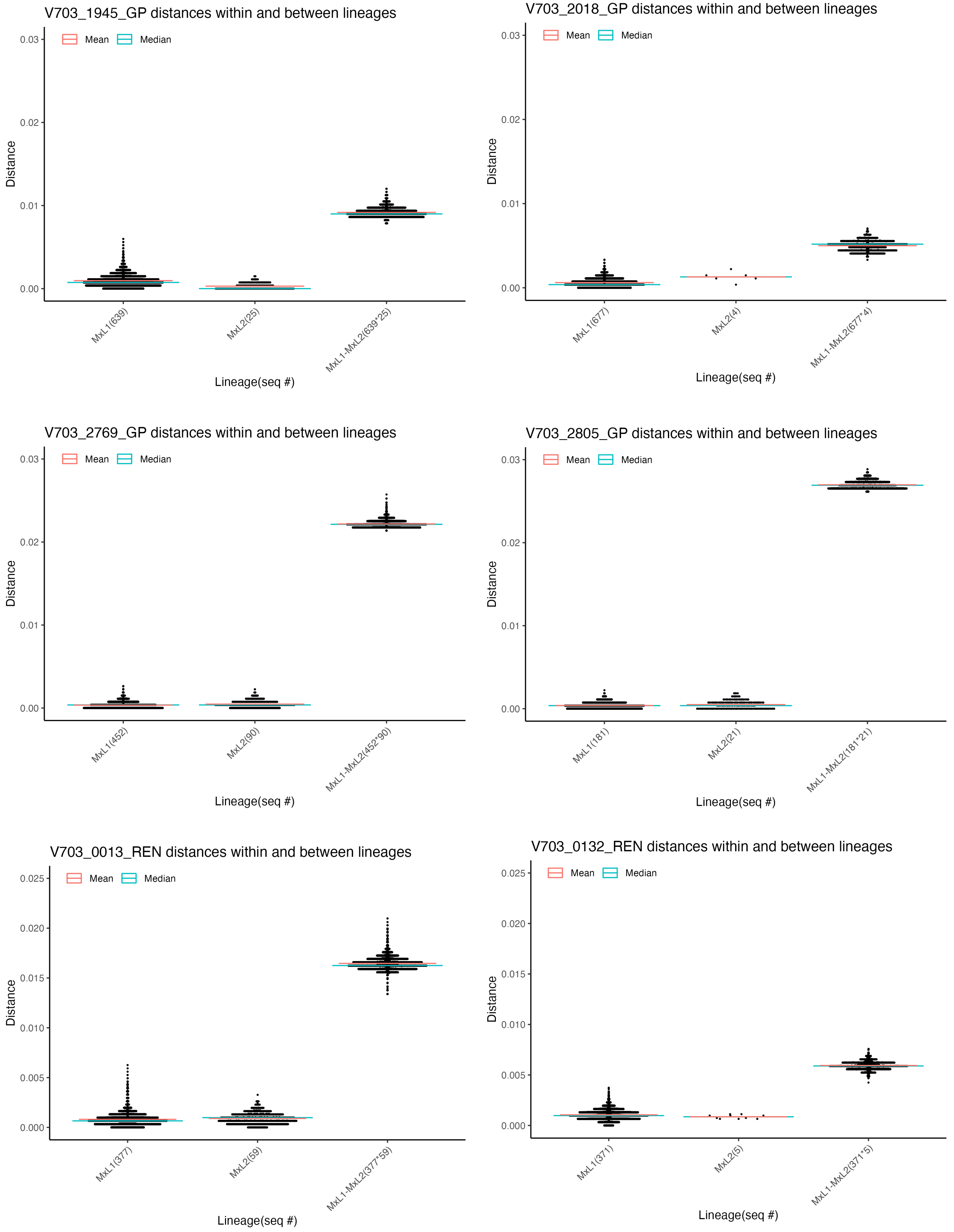
V703_1945_GP distances within and between lineages
Mean
Median
Distance
0.03
0.02
0.01
0.00
MxL1(639)
MxL2(25)
MxL1-MxL2(639*25)
Lineage(seq #)
V703_2018_GP distances within and between lineages
Mean
Median
Distance
0.03
0.02
0.01
0.00
MxL1(677)
MxL2(4)
MxL1-MxL2(677*4)
Lineage(seq #)
V703_2769_GP distances within and between lineages
Mean
Median
Distance
0.03
0.02
0.01
0.00
MxL1(452)
MxL2(90)
MxL1-MxL2(452*90)
Lineage(seq #)
V703_2805_GP distances within and between lineages
Mean
Median
Distance
0.03
0.02
0.01
0.00
MxL1(181)
MxL2(21)
MxL1-MxL2(181*21)
Lineage(seq #)
V703_0013_REN distances within and between lineages
Mean
Median
Distance
0.025
0.020
0.015
0.010
0.005
0.000
MxL1(377)
MxL2(59)
MxL1-MxL2(377*59)
Lineage(seq #)
V703_0132_REN distances within and between lineages
Mean
Median
Distance
0.025
0.020
0.015
0.010
0.005
0.000
MxL1(371)
MxL2(5)
MxL1-MxL2(371*5)
Lineage(seq #)

**S3.14 Fig.**

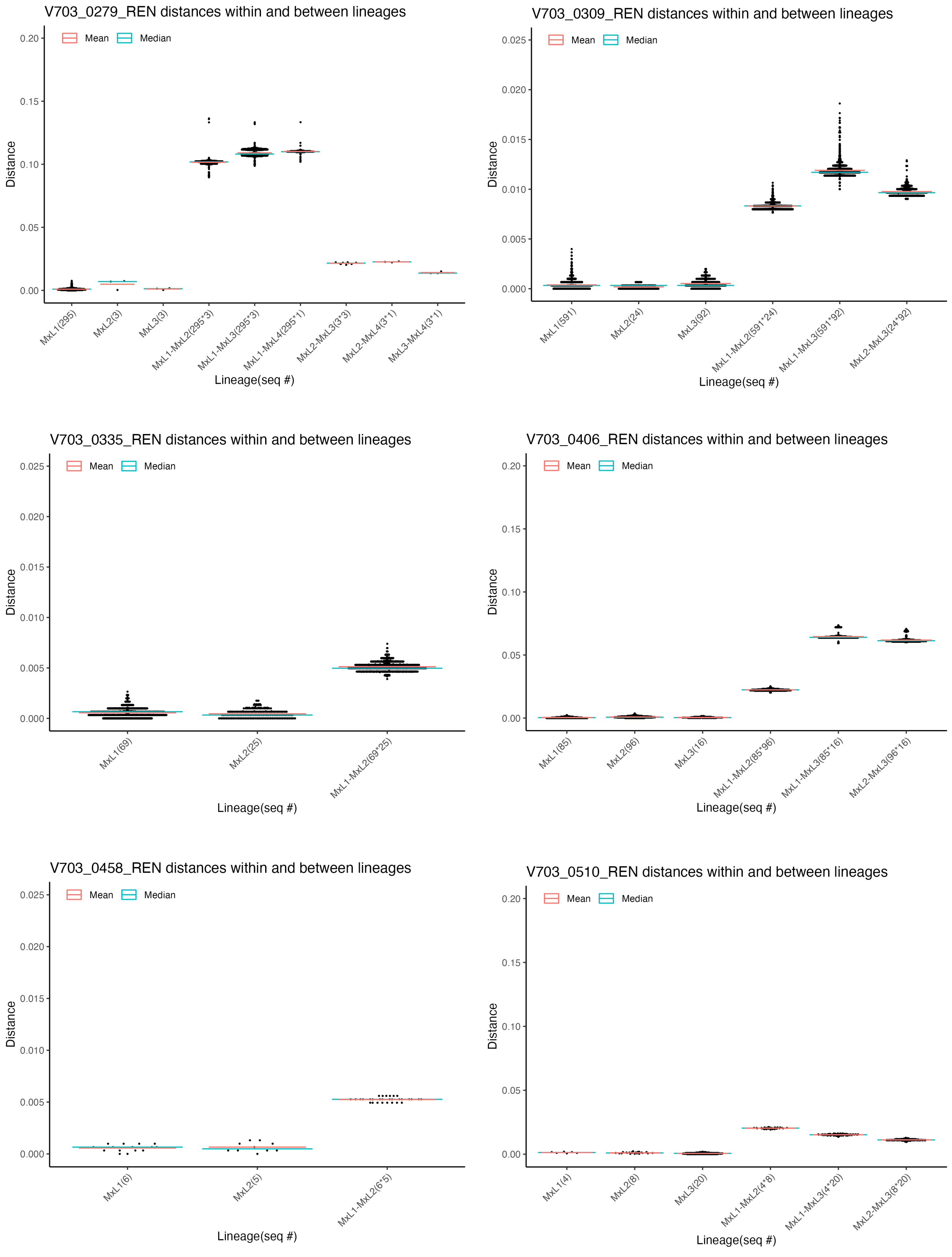
V703_0279_REN distances within and between lineages
Mean
Median
Distance
0.20
0.15
0.10
0.05
0.00
MxL1(295)
MxL2(3)
MxL3(3)
MxL1-MxL2(295*3)
MxL1-MxL3(295*3)
MxL1-MxL4(295*1)
MxL2-MxL3(3*3)
MxL2-MxL4(3*1)
MxL3-MxL4(3*1)
Lineage(seq #)
V703_0309_REN distances within and between lineages
Mean
Median
Distance
0.025
0.020
0.015
0.010
0.005
0.000
MxL1(591)
MxL2(24)
MxL3(92)
MxL1-MxL2(591*24)
MxL1-MxL3(591*92)
MxL2-MxL3(24*92)
Lineage(seq #)
V703_0335_REN distances within and between lineages
Mean
Median
Distance
0.025
0.020
0.015
0.010
0.005
0.000
MxL1(69)
MxL2(25)
MxL1-MxL2(69*25)
Lineage(seq #)
V703_0406_REN distances within and between lineages
Mean
Median
Distance
0.20
0.15
0.10
0.05
0.00
MxL1(85)
MxL2(96)
MxL3(16)
MxL1-MxL2(85*96)
MxL1-MxL3(85*16)
MxL2-MxL3(96*16)
Lineage(seq #)
V703_0458_REN distances within and between lineages
Mean
Median
Distance
0.025
0.020
0.015
0.010
0.005
0.000
MxL1(6)
MxL2(5)
MxL1-MxL2(6*5)
Lineage(seq #)
V703_0510_REN distances within and between lineages
Mean
Median
Distance
0.20
0.15
0.10
0.05
0.00
MxL1(4)
MxL2(8)
MxL3(20)
MxL1-MxL2(4*8)
MxL1-MxL3(4*20)
MxL2-MxL3(8*20)
Lineage(seq #)

**S3.15 Fig.**

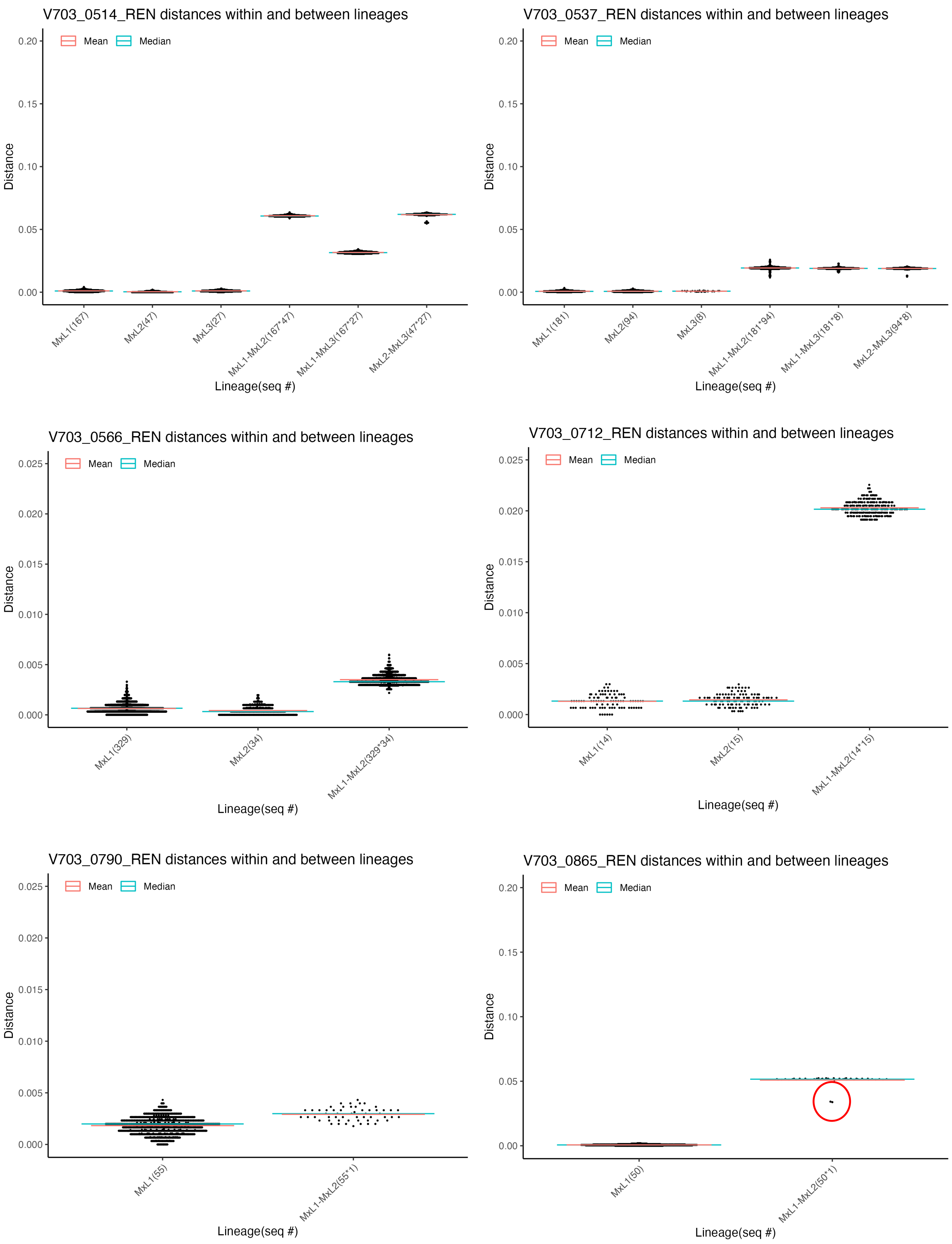

V703_0514_REN distances within and between lineages
Mean
Median
Distance
0.20
0.15
0.10
0.05
0.00
MxL1(167)
MxL2(47)
MxL3(27)
MxL1-MxL2(167*47)
MxL1-MxL3(167*27)
MxL2-MxL3(47*27)
Lineage(seq #)
V703_0537_REN distances within and between lineages
Mean
Median
Distance
0.20
0.15
0.10
0.05
0.00
MxL1(181)
MxL2(94)
MxL3(8)
MxL1-MxL2(181*94)
MxL1-MxL3(181*8)
MxL2-MxL3(94*8)
Lineage(seq #)
V703_0566_REN distances within and between lineages
Mean
Median
Distance
0.025
0.020
0.015
0.010
0.005
0.000
MxL1(329)
MxL2(34)
MxL1-MxL2(329*34)
Lineage(seq #)
V703_0712_REN distances within and between lineages
Mean
Median
Distance
0.025
0.020
0.015
0.010
0.005
0.000
MxL1(14)
MxL2(15)
MxL1-MxL2(14*15)
Lineage(seq #)
V703_0790_REN distances within and between lineages
Mean
Median
Distance
0.025
0.020
0.015
0.010
0.005
0.000
MxL1(55)
MxL1-MxL2(55*1)
Lineage(seq #)
V703_0865_REN distances within and between lineages
Mean
Median
Distance
0.20
0.15
0.10
0.05
0.00
MxL1(50)
MxL1-MxL2(50*1)
Lineage(seq #)

**S3.16 Fig.**

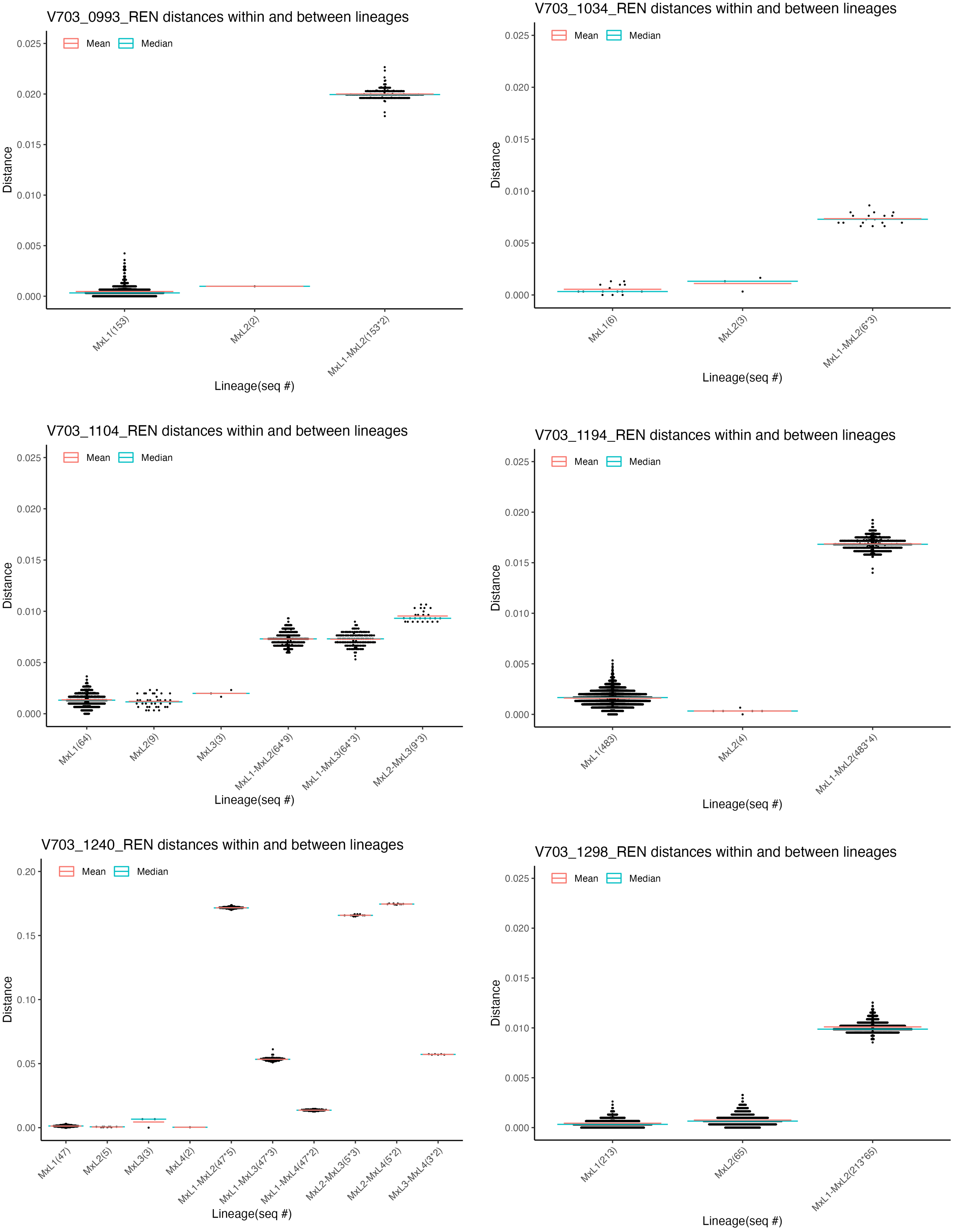
V703_0993_REN distances within and between lineages
Mean
Median
Distance
0.025
0.020
0.015
0.010
0.005
0.000
MxL1(153)
MxL2(2)
MxL1-MxL2(153*2)
Lineage(seq #)
V703_1034_REN distances within and between lineages
Mean
Median
Distance
0.025
0.020
0.015
0.010
0.005
0.000
MxL1(6)
MxL2(3)
MxL1-MxL2(6*3)
Lineage(seq #)
V703_1104_REN distances within and between lineages
Mean
Median
Distance
0.025
0.020
0.015
0.010
0.005
0.000
MxL1(64)
MxL2(9)
MxL3(3)
MxL1-MxL2(64*9)
MxL1-MxL3(64*3)
MxL2-MxL3(9*3)
Lineage(seq #)
V703_1194_REN distances within and between lineages
Mean
Median
Distance
0.025
0.020
0.015
0.010
0.005
0.000
MxL1(483)
MxL2(4)
MxL1-MxL2(483*4)
Lineage(seq #)
V703_1240_REN distances within and between lineages
Mean
Median
Distance
0.20
0.15
0.10
0.05
0.00
MxL1(47)
MxL2(5)
MxL3(3)
MxL4(2)
MxL1-MxL2(47*5)
MxL1-MxL3(47*3)
MxL1-MxL4(47*2)
MxL2-MxL3(5*3)
MxL2-MxL4(5*2)
MxL3-MxL4(3*2)
Lineage(seq #)
V703_1298_REN distances within and between lineages
Mean
Median
Distance
0.025
0.020
0.015
0.010
0.005
0.000
MxL1(213)
MxL2(65)
MxL1-MxL2(213*65)
Lineage(seq #)

**S3.17 Fig.**

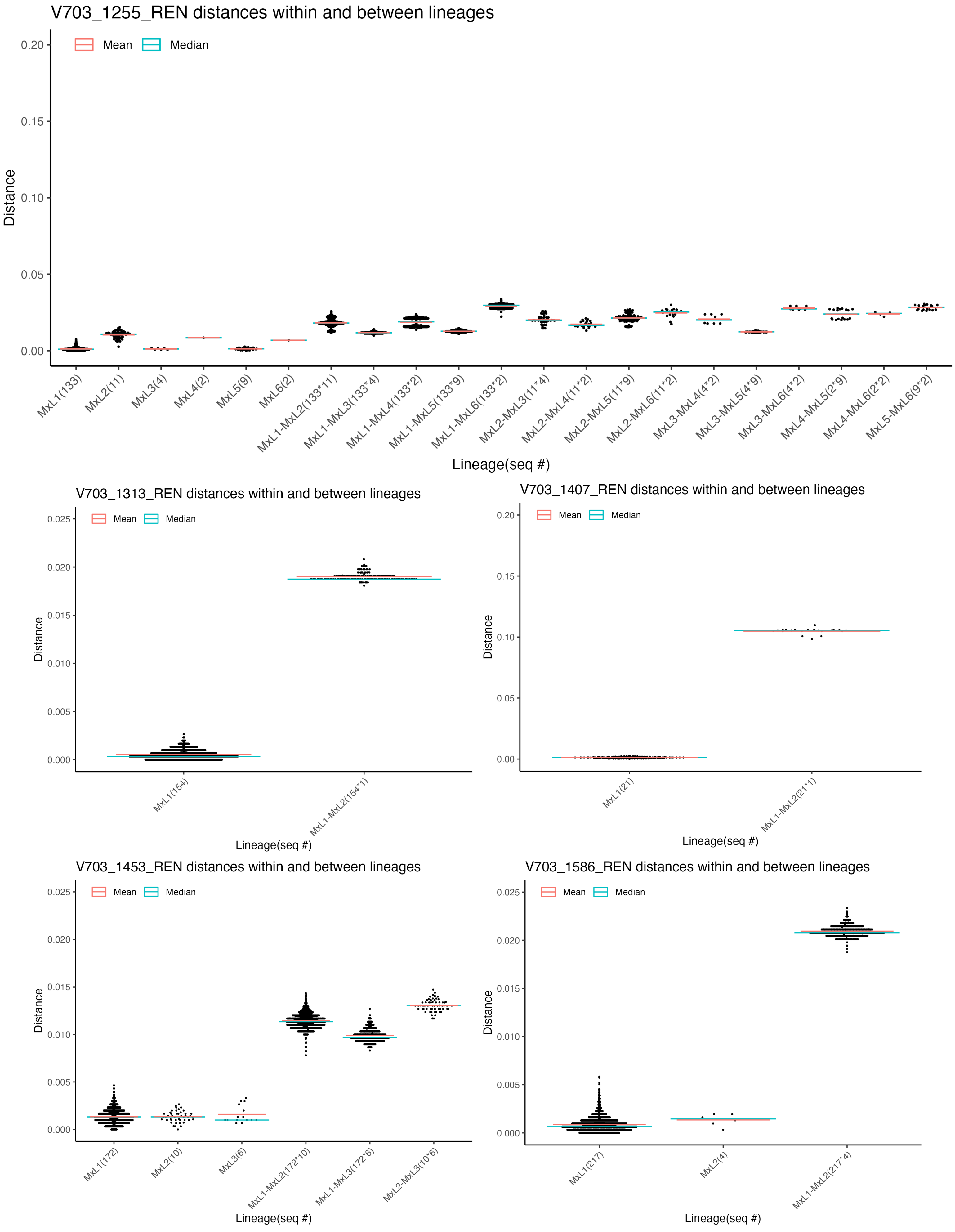
V703_1255_REN distances within and between lineages
Mean
Median
Distance
0.20
0.15
0.10
0.05
0.00
MxL1(133)
MxL2(11)
MxL3(4)
MxL4(2)
MxL5(9)
MxL6(2)
MxL1-MxL2(133*11)
MxL1-MxL3(133*4)
MxL1-MxL4(133*2)
MxL1-MxL5(133*9)
MxL1-MxL6(133*2)
MxL2-MxL3(11*4)
MxL2-MxL4(11*2)
MxL2-MxL5(11*9)
MxL2-MxL6(11*2)
MxL3-MxL4(4*2)
MxL3-MxL5(4*9)
MxL3-MxL6(4*2)
MxL4-MxL5(2*9)
MxL4-MxL6(2*2)
MxL5-MxL6(9*2)
Lineage(seq #)
V703_1313_REN distances within and between lineages
Mean
Median
Distance
0.025
0.020
0.015
0.010
0.005
0.000
MxL1(154)
MxL1-MxL2(154*1)
Lineage(seq #)
V703_1407_REN distances within and between lineages
Mean
Median
Distance
0.20
0.15
0.10
0.05
0.00
MxL1(21)
MxL1-MxL2(21*1)
Lineage(seq #)
V703_1453_REN distances within and between lineages
Mean
Median
Distance
0.025
0.020
0.015
0.010
0.005
0.000
MxL1(172)
MxL2(10)
MxL3(6)
MxL1-MxL2(172*10)
MxL1-MxL3(172*6)
MxL2-MxL3(10*6)
Lineage(seq #)
V703_1586_REN distances within and between lineages
Mean
Median
Distance
0.025
0.020
0.015
0.010
0.005
0.000
MxL1(217)
MxL2(4)
MxL1-MxL2(217*4)
Lineage(seq #)

**S3.18 Fig.**

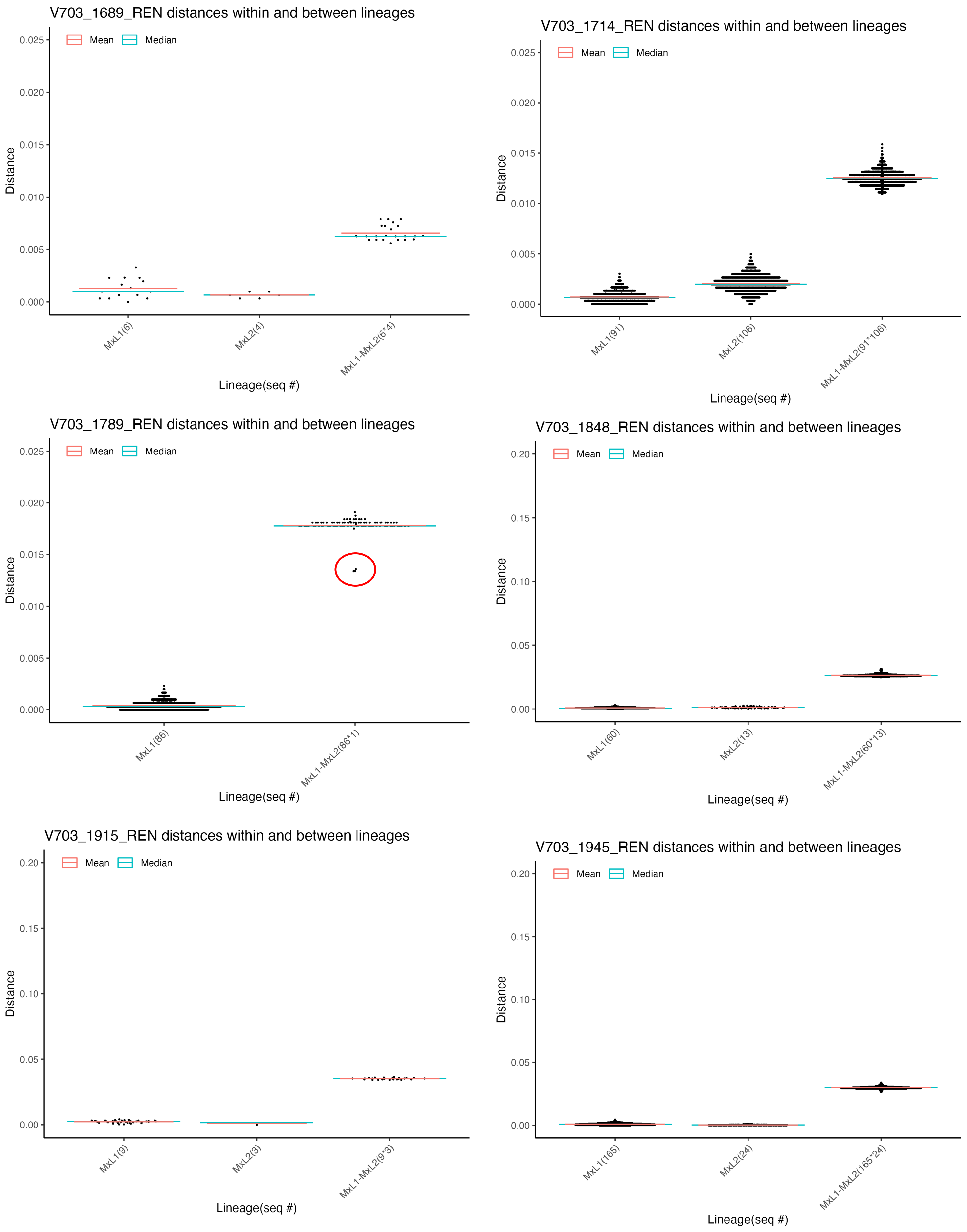
V703_1689_REN distances within and between lineages
Mean
Median
Distance
Lineage(seq #)
MxL1(6)
MxL2(4)
MxL1-MxL2(6*4)
V703_1714_REN distances within and between lineages
MxL1(91)
MxL2(106)
MxL1-MxL2(91*106)
V703_1789_REN distances within and between lineages
MxL1(86)
MxL1-MxL2(86*1)
V703_1848_REN distances within and between lineages
MxL1(60)
MxL2(13)
MxL1-MxL2(60*13)
V703_1915_REN distances within and between lineages
MxL1(9)
MxL2(3)
MxL1-MxL2(9*3)
V703_1945_REN distances within and between lineages
MxL1(165)
MxL2(24)
MxL1-MxL2(165*24)

**S3.19 Fig.**

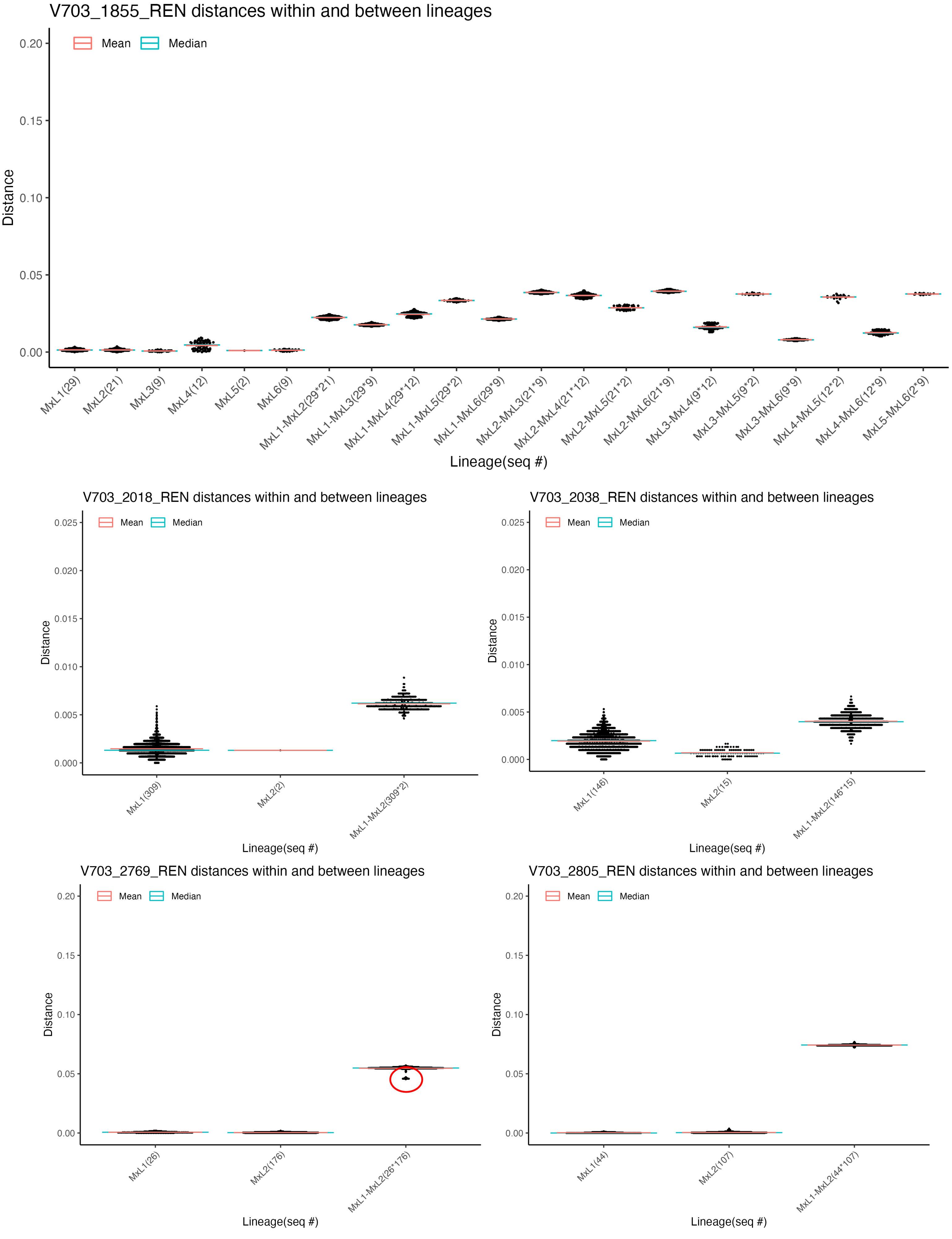
V703_1855_REN distances within and between lineages
Mean
Median
Distance
0.20
0.15
0.10
0.05
0.00
MxL1(29)
MxL2(21)
MxL3(9)
MxL4(12)
MxL5(2)
MxL6(9)
MxL1-MxL2(29*21)
MxL1-MxL3(29*9)
MxL1-MxL4(29*12)
MxL1-MxL5(29*2)
MxL1-MxL6(29*9)
MxL2-MxL3(21*9)
MxL2-MxL4(21*12)
MxL2-MxL5(21*2)
MxL2-MxL6(21*9)
MxL3-MxL4(9*12)
MxL3-MxL5(9*2)
MxL3-MxL6(9*9)
MxL4-MxL5(12*2)
MxL4-MxL6(12*9)
MxL5-MxL6(2*9)
Lineage(seq #)
V703_2018_REN distances within and between lineages
Mean
Median
Distance
0.025
0.020
0.015
0.010
0.005
0.000
MxL1(309)
MxL2(2)
MxL1-MxL2(309*2)
Lineage(seq #)
V703_2038_REN distances within and between lineages
Mean
Median
Distance
0.025
0.020
0.015
0.010
0.005
0.000
MxL1(146)
MxL2(15)
MxL1-MxL2(146*15)
Lineage(seq #)
V703_2769_REN distances within and between lineages
Mean
Median
Distance
0.20
0.15
0.10
0.05
0.00
MxL1(26)
MxL2(176)
MxL1-MxL2(26*176)
Lineage(seq #)
V703_2805_REN distances within and between lineages
Mean
Median
Distance
0.20
0.15
0.10
0.05
0.00
MxL1(44)
MxL2(107)
MxL1-MxL2(44*107)
Lineage(seq #)

**S3.20 Fig.**

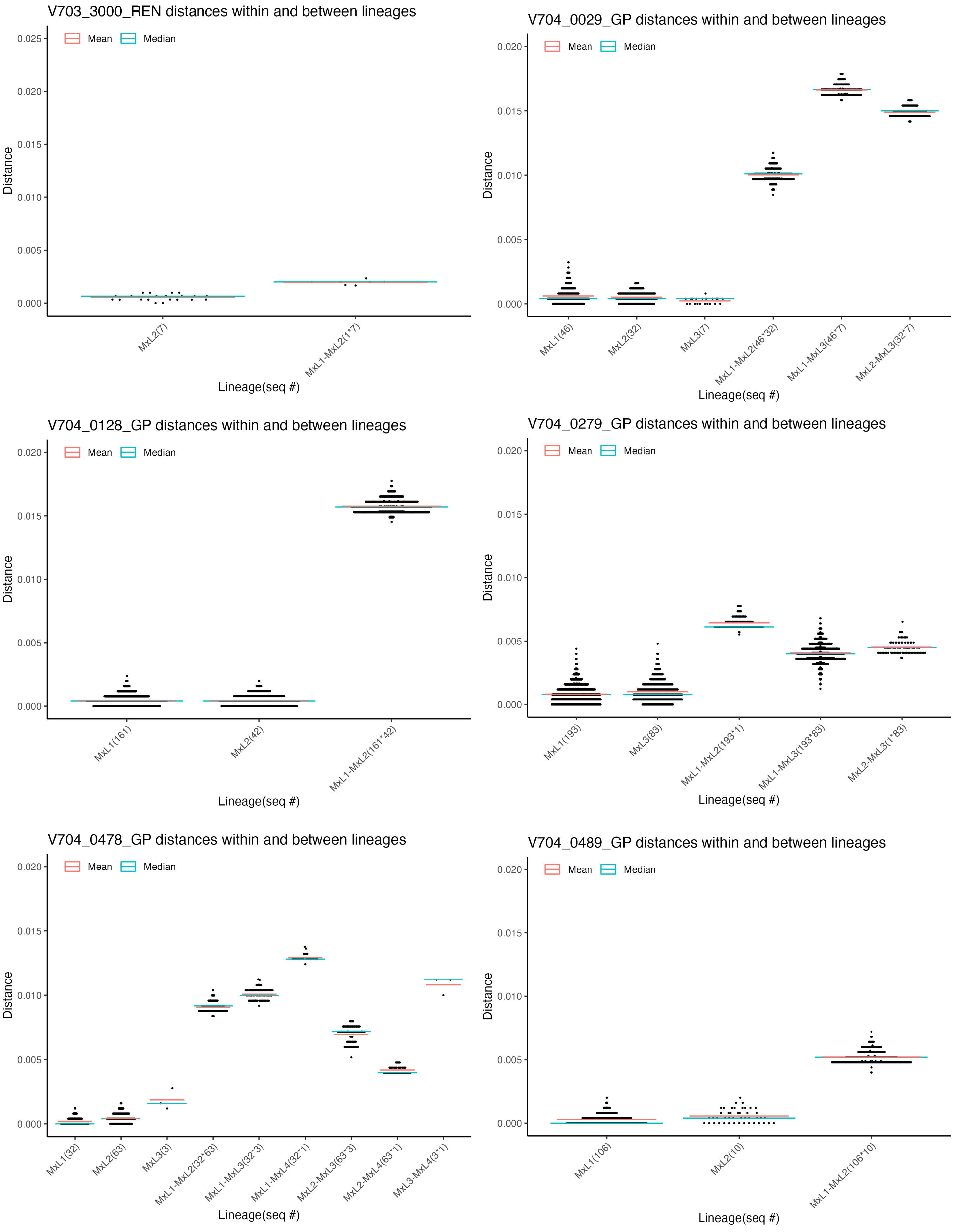

V703_3000_REN distances within and between lineages
Mean
Median
Distance
0.025
0.020
0.015
0.010
0.005
0.000
MxL2(7)
MxL1-MxL2(1*7)
Lineage(seq #)
V704_0029_GP distances within and between lineages
Mean
Median
Distance
0.020
0.015
0.010
0.005
0.000
MxL1(46)
MxL2(32)
MxL3(7)
MxL1-MxL2(46*32)
MxL1-MxL3(46*7)
MxL2-MxL3(32*7)
Lineage(seq #)
V704_0128_GP distances within and between lineages
Mean
Median
Distance
0.020
0.015
0.010
0.005
0.000
MxL1(161)
MxL2(42)
MxL1-MxL2(161*42)
Lineage(seq #)
V704_0279_GP distances within and between lineages
Mean
Median
Distance
0.020
0.015
0.010
0.005
0.000
MxL1(193)
MxL3(83)
MxL1-MxL2(193*1)
MxL1-MxL3(193*83)
MxL2-MxL3(1*83)
Lineage(seq #)
V704_0478_GP distances within and between lineages
Mean
Median
Distance
0.020
0.015
0.010
0.005
0.000
MxL1(32)
MxL2(63)
MxL3(3)
MxL1-MxL2(32*63)
MxL1-MxL3(32*3)
MxL1-MxL4(32*1)
MxL2-MxL3(63*3)
MxL2-MxL4(63*1)
MxL3-MxL4(3*1)
Lineage(seq #)
V704_0489_GP distances within and between lineages
Mean
Median
Distance
0.020
0.015
0.010
0.005
0.000
MxL1(106)
MxL2(10)
MxL1-MxL2(106*10)
Lineage(seq #)

**S3.21 Fig.**

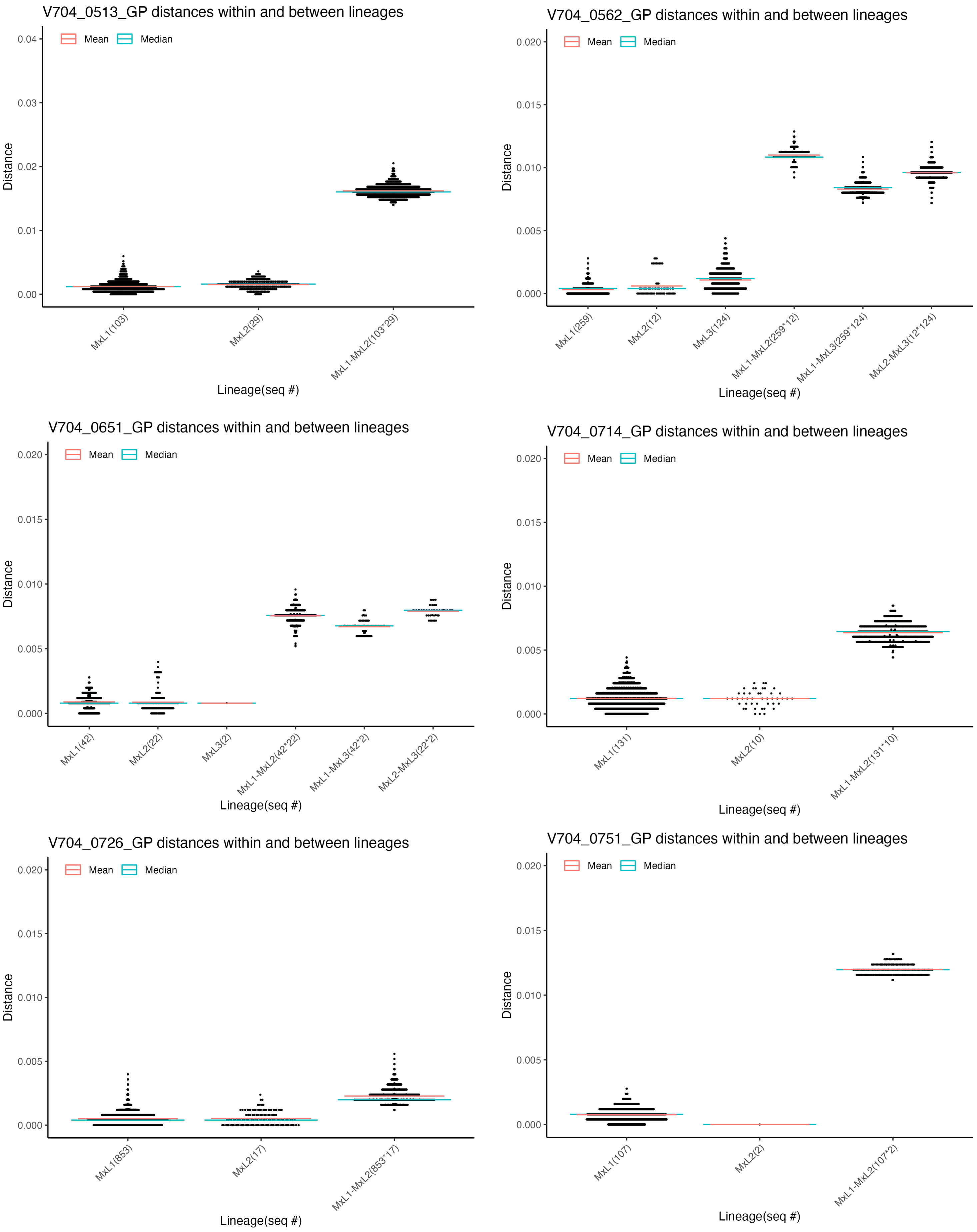
V704_0513_GP distances within and between lineages
Mean
Median
Distance
0.04
0.03
0.02
0.01
0.00
MxL1(103)
MxL2(29)
MxL1-MxL2(103*29)
Lineage(seq #)
V704_0562_GP distances within and between lineages
Mean
Median
Distance
0.020
0.015
0.010
0.005
0.000
MxL1(259)
MxL2(12)
MxL3(124)
MxL1-MxL2(259*12)
MxL1-MxL3(259*124)
MxL2-MxL3(12*124)
Lineage(seq #)
V704_0651_GP distances within and between lineages
Mean
Median
Distance
0.020
0.015
0.010
0.005
0.000
MxL1(42)
MxL2(22)
MxL3(2)
MxL1-MxL2(42*22)
MxL1-MxL3(42*2)
MxL2-MxL3(22*2)
Lineage(seq #)
V704_0714_GP distances within and between lineages
Mean
Median
Distance
0.020
0.015
0.010
0.005
0.000
MxL1(131)
MxL2(10)
MxL1-MxL2(131*10)
Lineage(seq #)
V704_0726_GP distances within and between lineages
Mean
Median
Distance
0.020
0.015
0.010
0.005
0.000
MxL1(853)
MxL2(17)
MxL1-MxL2(853*17)
Lineage(seq #)
V704_0751_GP distances within and between lineages
Mean
Median
Distance
0.020
0.015
0.010
0.005
0.000
MxL1(107)
MxL2(2)
MxL1-MxL2(107*2)
Lineage(seq #)

**S3.22 Fig.**

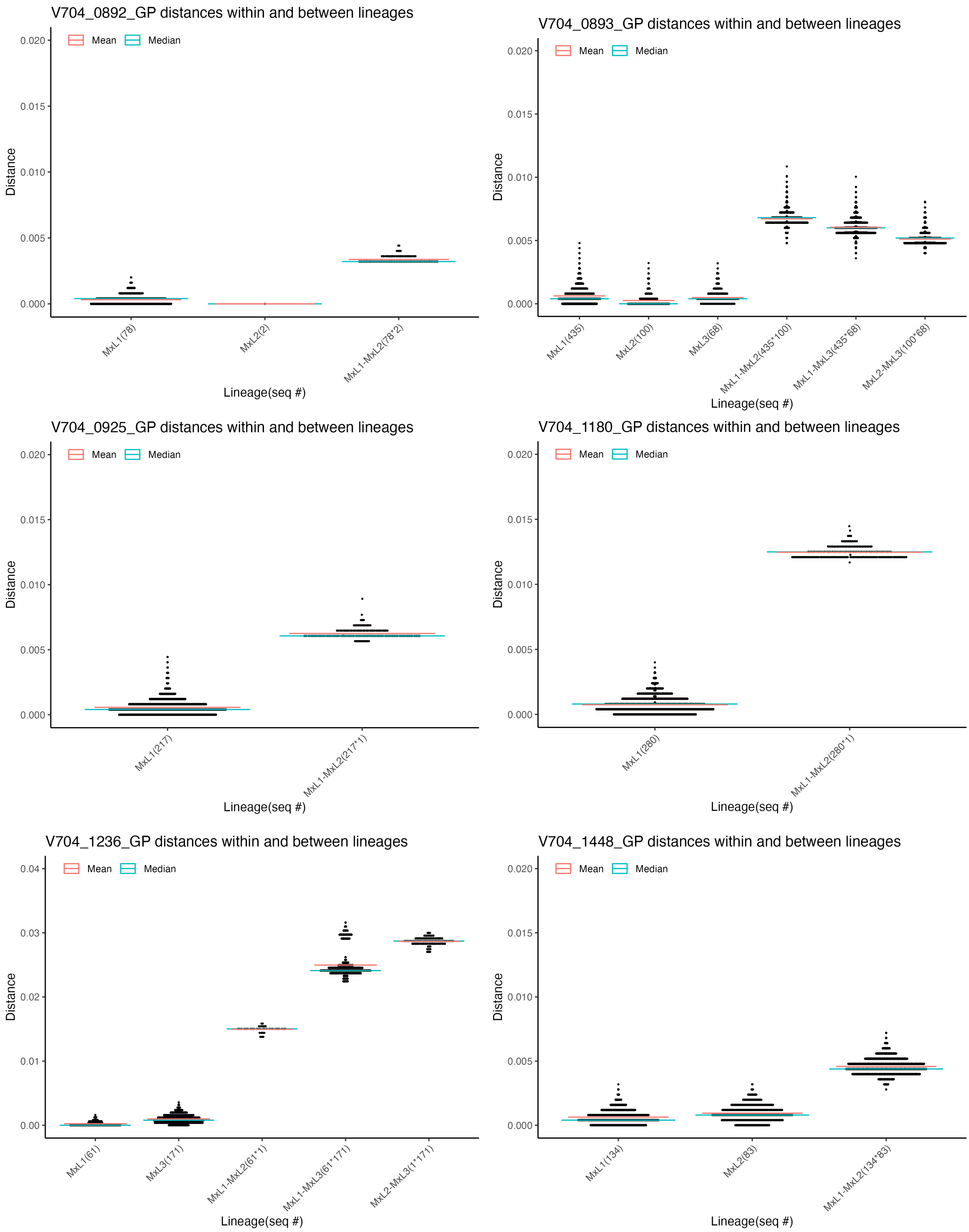
V704_0892_GP distances within and between lineages
Mean
Median
Distance
0.000
0.005
0.010
0.015
0.020
MxL1(78)
MxL2(2)
MxL1-MxL2(78*2)
Lineage(seq #)
V704_0893_GP distances within and between lineages
MxL1(435)
MxL2(100)
MxL3(68)
MxL1-MxL2(435*100)
MxL1-MxL3(435*68)
MxL2-MxL3(100*68)
V704_0925_GP distances within and between lineages
MxL1(217)
MxL1-MxL2(217*1)
V704_1180_GP distances within and between lineages
MxL1(280)
MxL1-MxL2(280*1)
V704_1236_GP distances within and between lineages
0.00
0.01
0.02
0.03
0.04
MxL1(61)
MxL3(171)
MxL1-MxL2(61*1)
MxL1-MxL3(61*171)
MxL2-MxL3(1*171)
V704_1448_GP distances within and between lineages
MxL1(134)
MxL2(83)
MxL1-MxL2(134*83)

**S3.23 Fig.**

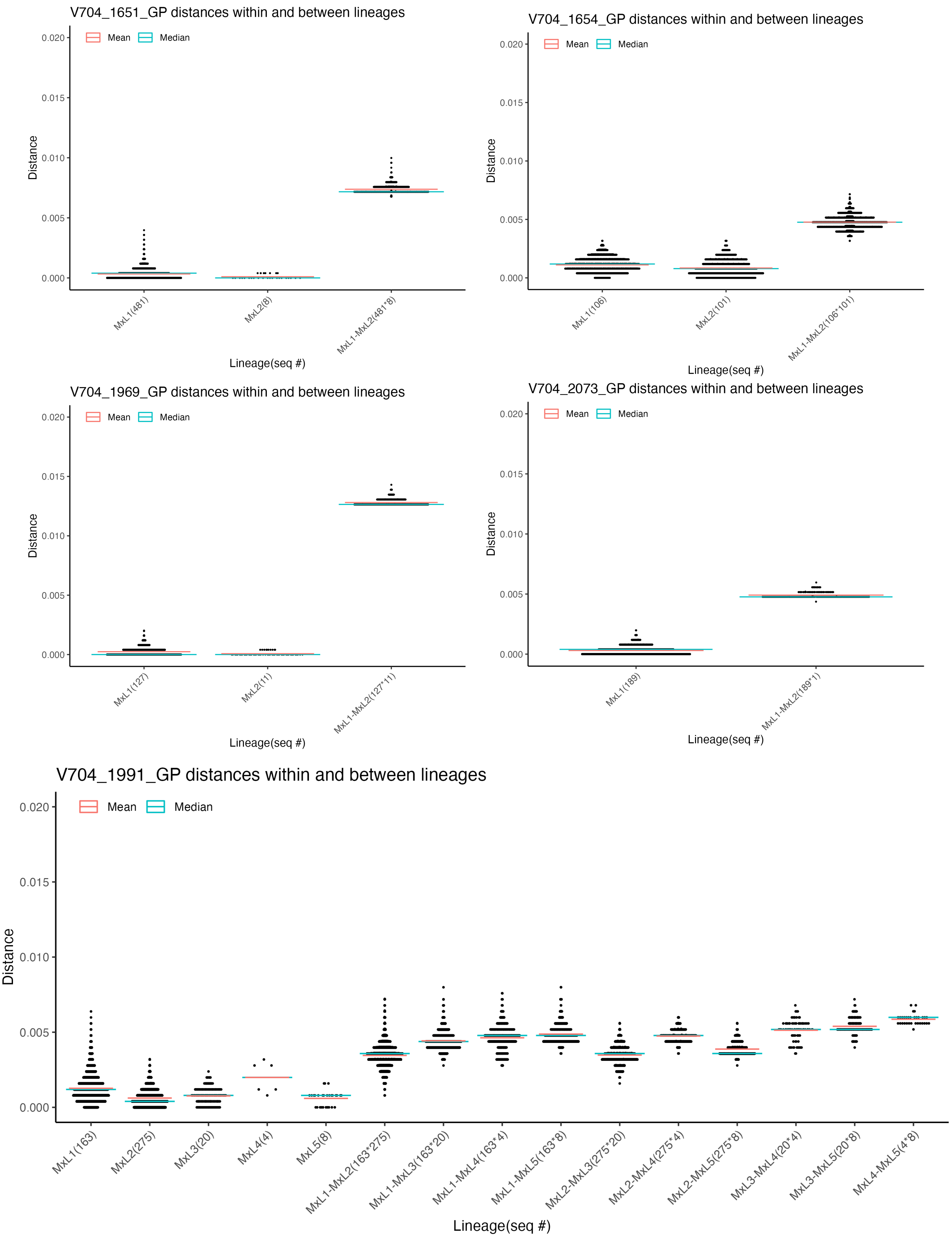
V704_1651_GP distances within and between lineages
Mean
Median
Distance
0.020
0.015
0.010
0.005
0.000
MxL1(481)
MxL2(8)
MxL1-MxL2(481*8)
Lineage(seq #)
V704_1654_GP distances within and between lineages
Mean
Median
Distance
0.020
0.015
0.010
0.005
0.000
MxL1(106)
MxL2(101)
MxL1-MxL2(106*101)
Lineage(seq #)
V704_1969_GP distances within and between lineages
Mean
Median
Distance
0.020
0.015
0.010
0.005
0.000
MxL1(127)
MxL2(11)
MxL1-MxL2(127*11)
Lineage(seq #)
V704_2073_GP distances within and between lineages
Mean
Median
Distance
0.020
0.015
0.010
0.005
0.000
MxL1(189)
MxL1-MxL2(189*1)
Lineage(seq #)
V704_1991_GP distances within and between lineages
Mean
Median
Distance
0.020
0.015
0.010
0.005
0.000
MxL1(163)
MxL2(275)
MxL3(20)
MxL4(4)
MxL5(8)
MxL1-MxL2(163*275)
MxL1-MxL3(163*20)
MxL1-MxL4(163*4)
MxL1-MxL5(163*8)
MxL2-MxL3(275*20)
MxL2-MxL4(275*4)
MxL2-MxL5(275*8)
MxL3-MxL4(20*4)
MxL3-MxL5(20*8)
MxL4-MxL5(4*8)
Lineage(seq #)

**S3.24 Fig.**

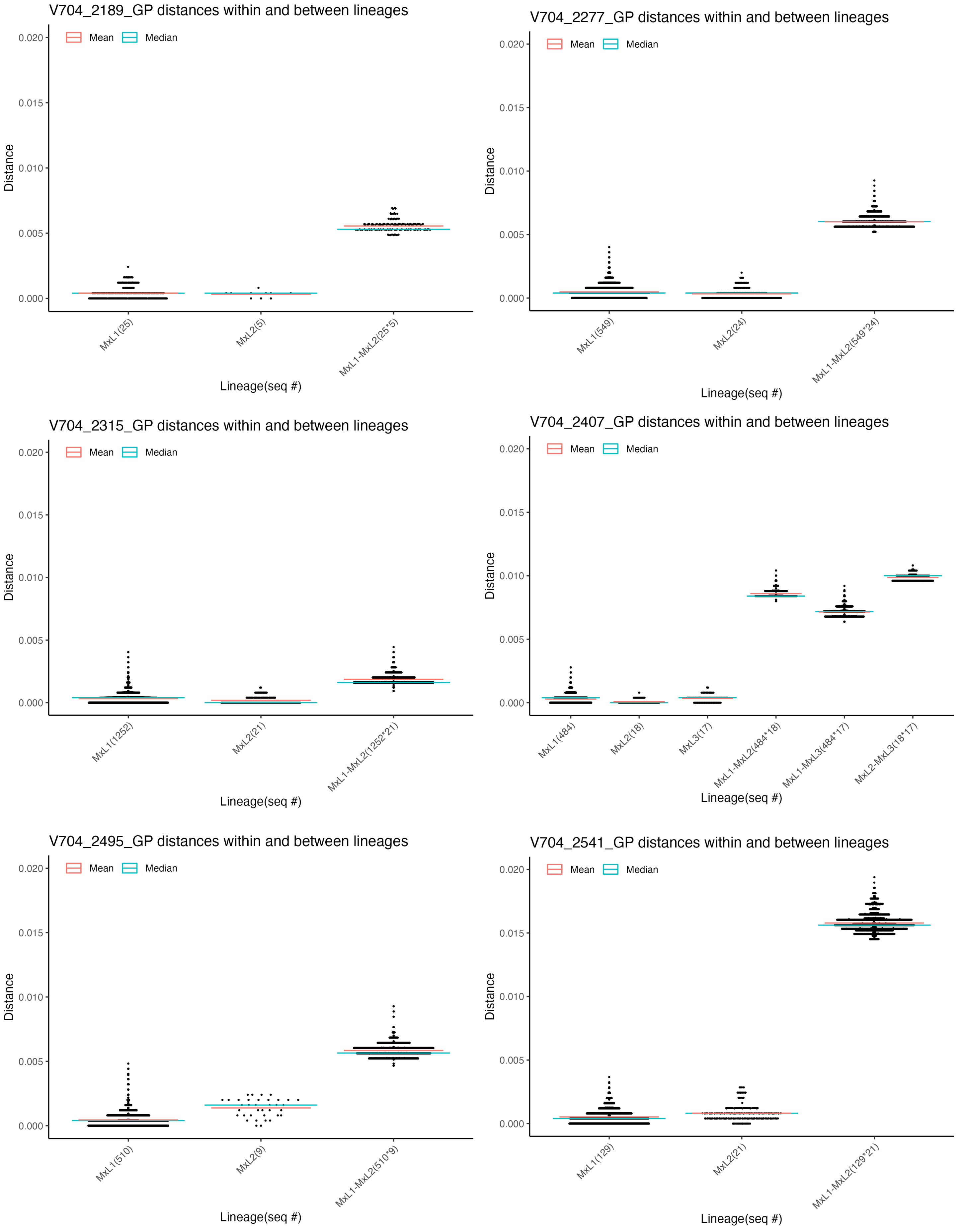

V704_2189_GP distances within and between lineages
Mean
Median
Distance
0.020
0.015
0.010
0.005
0.000
MxL1(25)
MxL2(5)
MxL1-MxL2(25*5)
Lineage(seq #)
V704_2277_GP distances within and between lineages
Mean
Median
Distance
0.020
0.015
0.010
0.005
0.000
MxL1(549)
MxL2(24)
MxL1-MxL2(549*24)
Lineage(seq #)
V704_2315_GP distances within and between lineages
Mean
Median
Distance
0.020
0.015
0.010
0.005
0.000
MxL1(1252)
MxL2(21)
MxL1-MxL2(1252*21)
Lineage(seq #)
V704_2407_GP distances within and between lineages
Mean
Median
Distance
0.020
0.015
0.010
0.005
0.000
MxL1(484)
MxL2(18)
MxL3(17)
MxL1-MxL2(484*18)
MxL1-MxL3(484*17)
MxL2-MxL3(18*17)
Lineage(seq #)
V704_2495_GP distances within and between lineages
Mean
Median
Distance
0.020
0.015
0.010
0.005
0.000
MxL1(510)
MxL2(9)
MxL1-MxL2(510*9)
Lineage(seq #)
V704_2541_GP distances within and between lineages
Mean
Median
Distance
0.020
0.015
0.010
0.005
0.000
MxL1(129)
MxL2(21)
MxL1-MxL2(129*21)
Lineage(seq #)

**S3.25 Fig.**

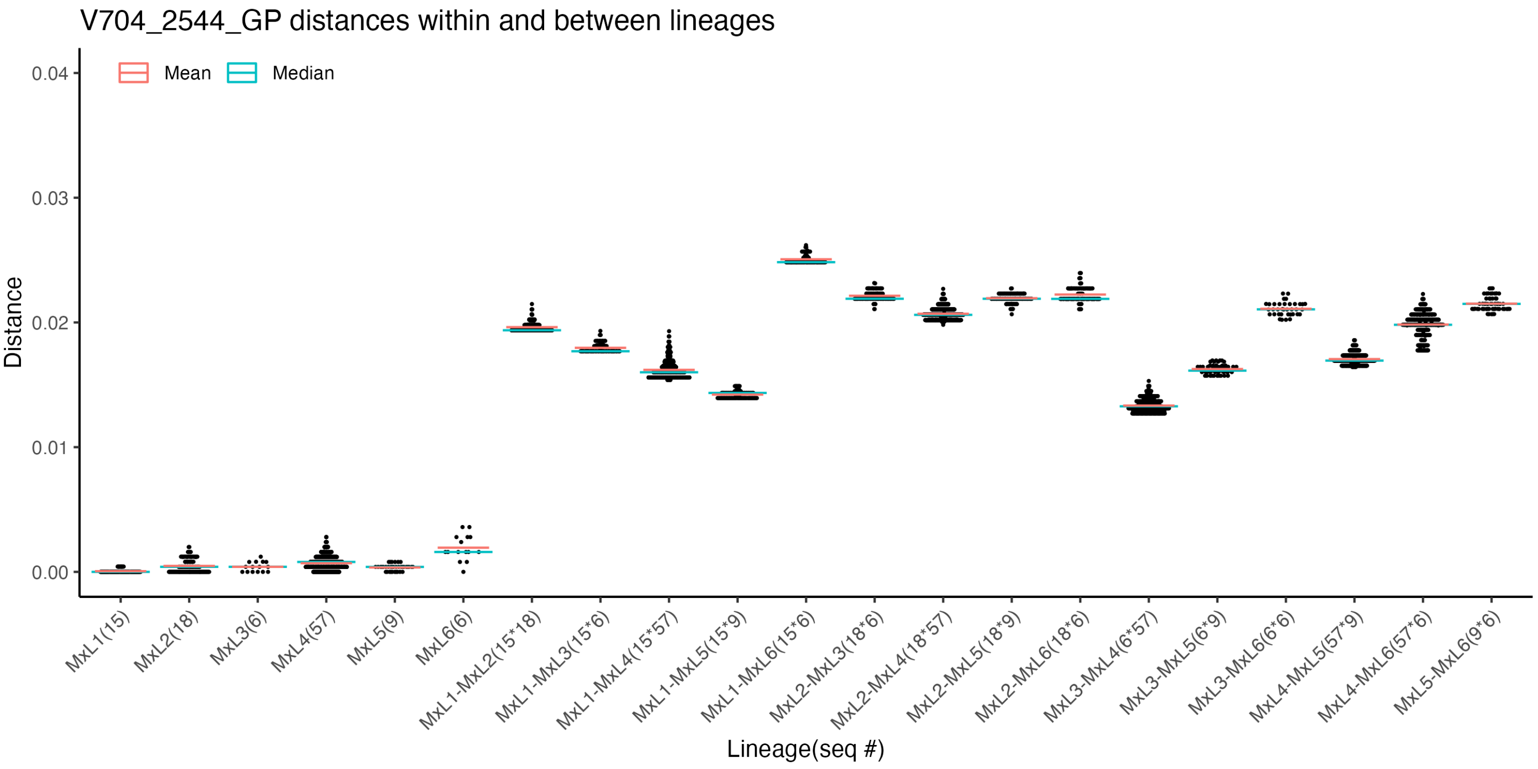

V704_2544_GP distances within and between lineages
Mean
Median
Distance
0.04
0.03
0.02
0.01
0.00
MxL1(15)
MxL2(18)
MxL3(6)
MxL4(57)
MxL5(9)
MxL6(6)
MxL1-MxL2(15*18)
MxL1-MxL3(15*6)
MxL1-MxL4(15*57)
MxL1-MxL5(15*9)
MxL1-MxL6(15*6)
MxL2-MxL3(18*6)
MxL2-MxL4(18*57)
MxL2-MxL5(18*9)
MxL2-MxL6(18*6)
MxL3-MxL4(6*57)
MxL3-MxL5(6*9)
MxL3-MxL6(6*6)
MxL4-MxL5(57*9)
MxL4-MxL6(57*6)
MxL5-MxL6(9*6)
Lineage(seq #)

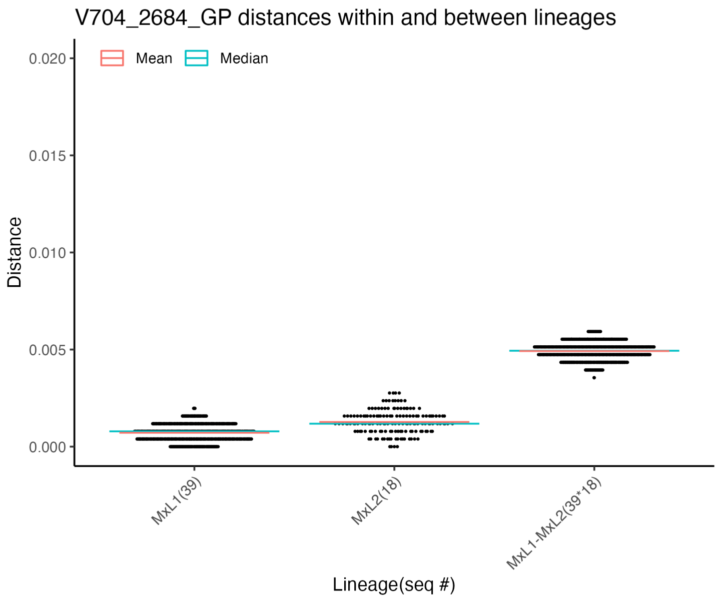

V704_2684_GP distances within and between lineages
Mean
Median
Distance
0.020
0.015
0.010
0.005
0.000
MxL1(39)
MxL2(18)
MxL1-MxL2(39*18)
Lineage(seq #)

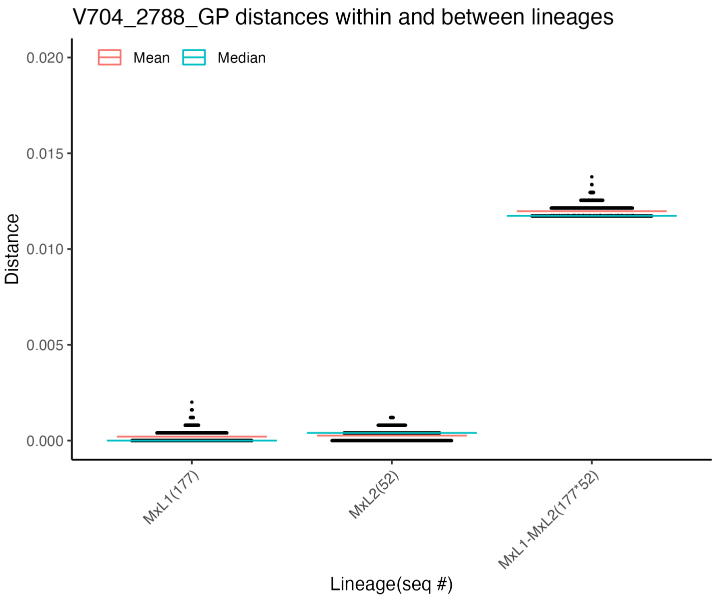

V704_2788_GP distances within and between lineages
Mean
Median
Distance
0.020
0.015
0.010
0.005
0.000
MxL1(177)
MxL2(52)
MxL1-MxL2(177*52)
Lineage(seq #)

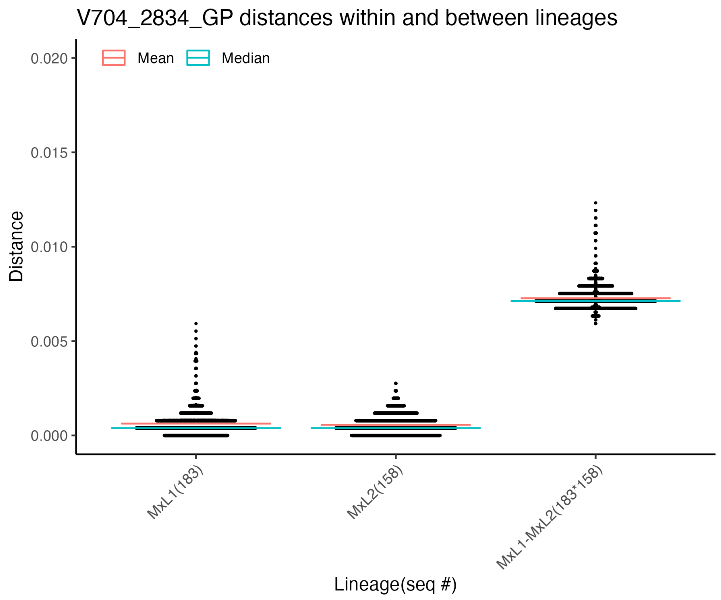

V704_2834_GP distances within and between lineages
Mean
Median
Distance
0.020
0.015
0.010
0.005
0.000
MxL1(183)
MxL2(158)
MxL1-MxL2(183*158)
Lineage(seq #)

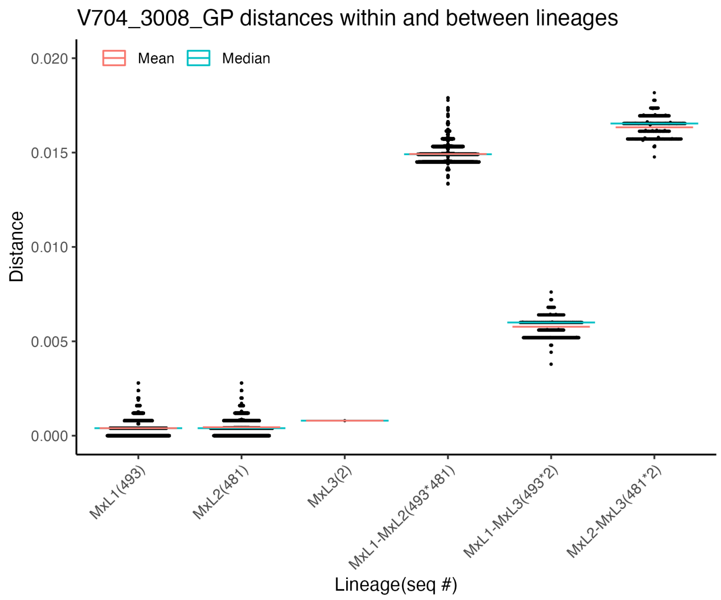

V704_3008_GP distances within and between lineages
Mean
Median
Distance
0.020
0.015
0.010
0.005
0.000
MxL1(493)
MxL2(481)
MxL3(2)
MxL1-MxL2(493*481)
MxL1-MxL3(493*2)
MxL2-MxL3(481*2)
Lineage(seq #)

**S3.26 Fig.**

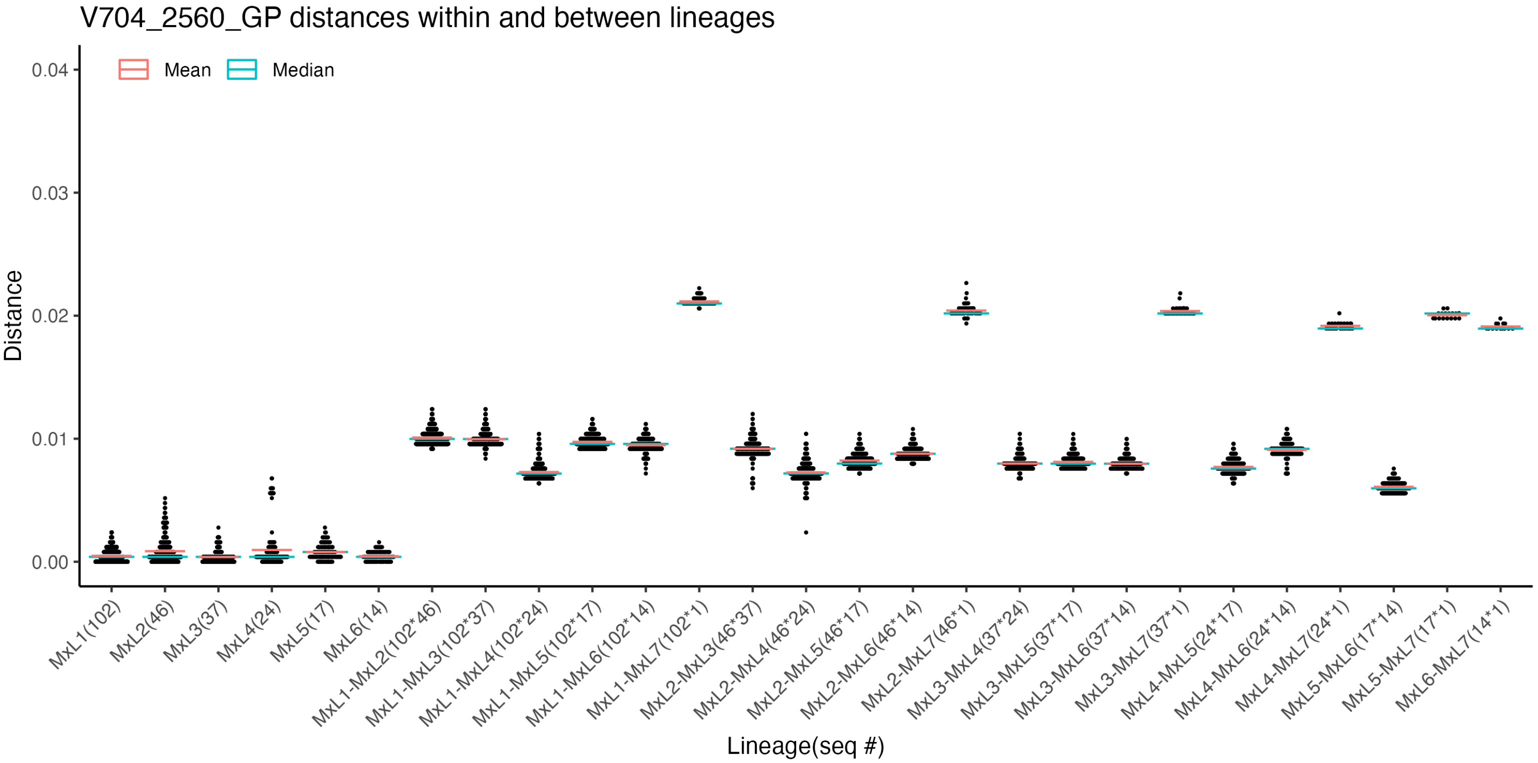
V704_2560_GP distances within and between lineages
Mean
Median
Distance
0.04
0.03
0.02
0.01
0.00
MxL1(102)
MxL2(46)
MxL3(37)
MxL4(24)
MxL5(17)
MxL6(14)
MxL1-MxL2(102*46)
MxL1-MxL3(102*37)
MxL1-MxL4(102*24)
MxL1-MxL5(102*17)
MxL1-MxL6(102*14)
MxL1-MxL7(102*1)
MxL2-MxL3(46*37)
MxL2-MxL4(46*24)
MxL2-MxL5(46*17)
MxL2-MxL6(46*14)
MxL2-MxL7(46*1)
MxL3-MxL4(37*24)
MxL3-MxL5(37*17)
MxL3-MxL6(37*14)
MxL3-MxL7(37*1)
MxL4-MxL5(24*17)
MxL4-MxL6(24*14)
MxL4-MxL7(24*1)
MxL5-MxL6(17*14)
MxL5-MxL7(17*1)
MxL6-MxL7(14*1)
Lineage(seq #)

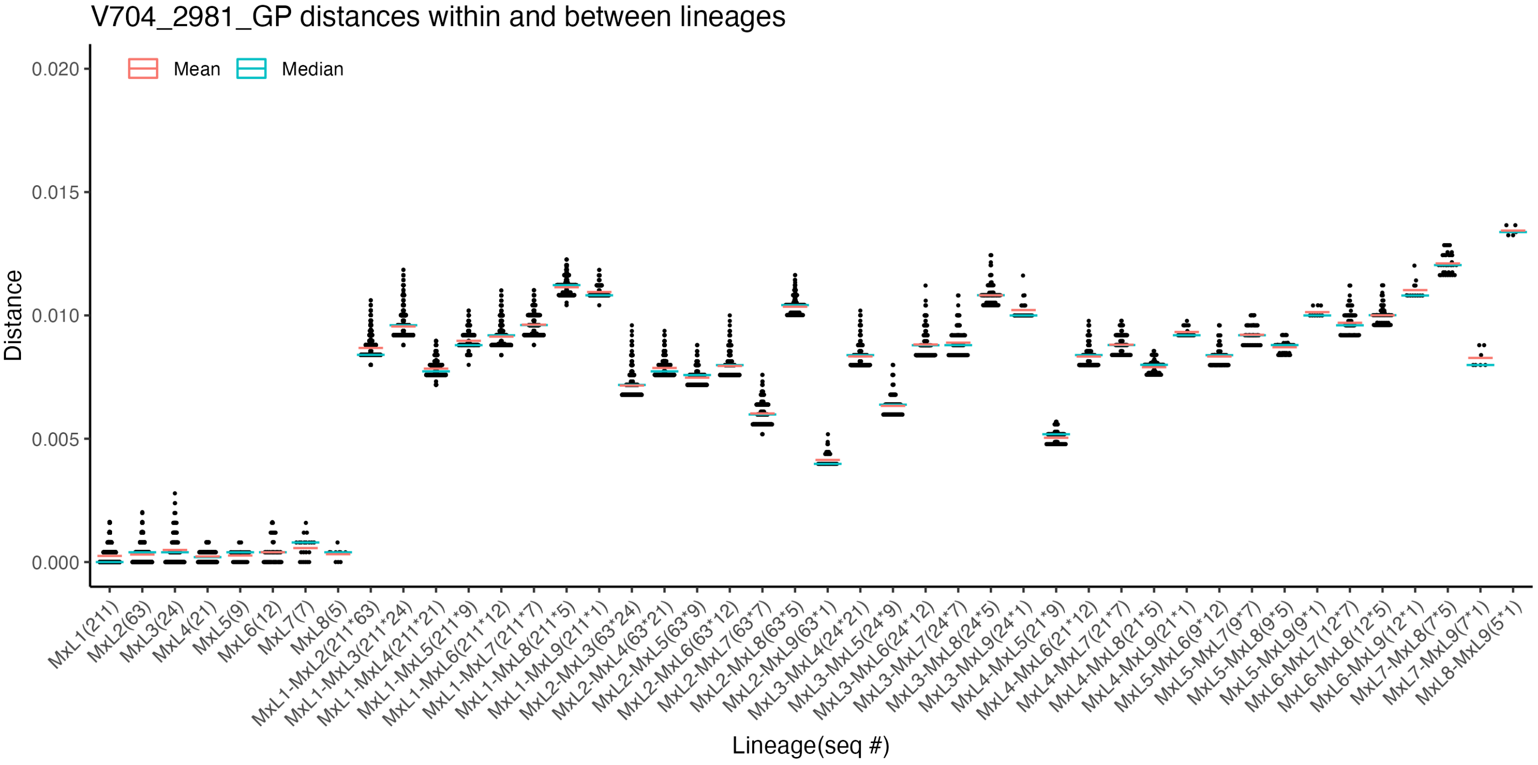
V704_2981_GP distances within and between lineages
Mean
Median
Distance
0.020
0.015
0.010
0.005
0.000
MxL1(211)
MxL2(63)
MxL3(24)
MxL4(21)
MxL5(9)
MxL6(12)
MxL7(7)
MxL8(5)
MxL1-MxL2(211*63)
MxL1-MxL3(211*24)
MxL1-MxL4(211*21)
MxL1-MxL5(211*9)
MxL1-MxL6(211*12)
MxL1-MxL7(211*7)
MxL1-MxL8(211*5)
MxL1-MxL9(211*1)
MxL2-MxL3(63*24)
MxL2-MxL4(63*21)
MxL2-MxL5(63*9)
MxL2-MxL6(63*12)
MxL2-MxL7(63*7)
MxL2-MxL8(63*5)
MxL2-MxL9(63*1)
MxL3-MxL4(24*21)
MxL3-MxL5(24*9)
MxL3-MxL6(24*12)
MxL3-MxL7(24*7)
MxL3-MxL8(24*5)
MxL3-MxL9(24*1)
MxL4-MxL5(21*9)
MxL4-MxL6(21*12)
MxL4-MxL7(21*7)
MxL4-MxL8(21*5)
MxL4-MxL9(21*1)
MxL5-MxL6(9*12)
MxL5-MxL7(9*7)
MxL5-MxL8(9*5)
MxL5-MxL9(9*1)
MxL6-MxL7(12*7)
MxL6-MxL8(12*5)
MxL6-MxL9(12*1)
MxL7-MxL8(7*5)
MxL7-MxL9(7*1)
MxL8-MxL9(5*1)
Lineage(seq #)

**S3.27 Fig.**

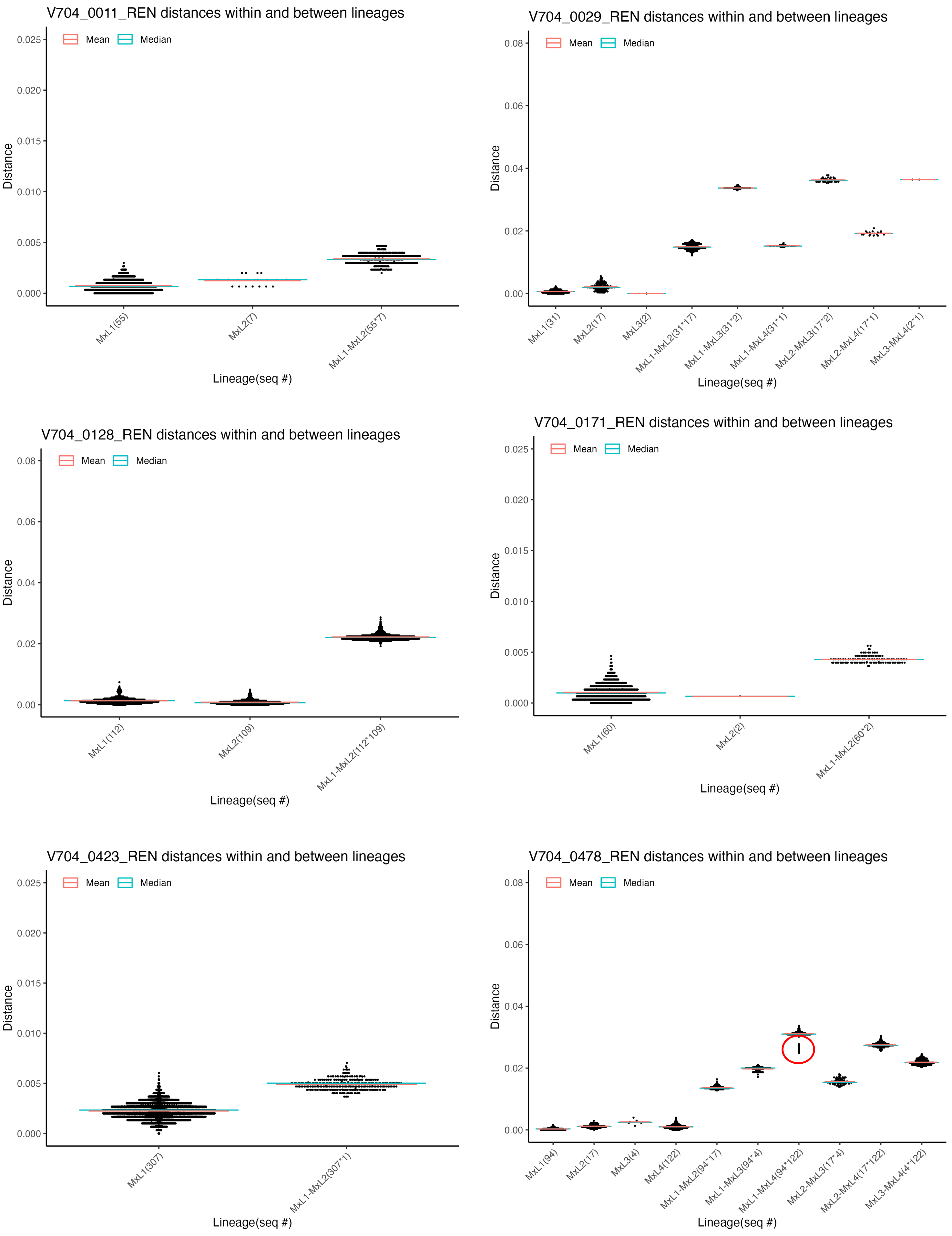
V704_0011_REN distances within and between lineages
Mean
Median
Distance
0.025
0.020
0.015
0.010
0.005
0.000
MxL1(55)
MxL2(7)
MxL1-MxL2(55*7)
Lineage(seq #)
V704_0029_REN distances within and between lineages
Mean
Median
Distance
0.08
0.06
0.04
0.02
0.00
MxL1(31)
MxL2(17)
MxL3(2)
MxL1-MxL2(31*17)
MxL1-MxL3(31*2)
MxL1-MxL4(31*1)
MxL2-MxL3(17*2)
MxL2-MxL4(17*1)
MxL3-MxL4(2*1)
Lineage(seq #)
V704_0128_REN distances within and between lineages
Mean
Median
Distance
0.08
0.06
0.04
0.02
0.00
MxL1(112)
MxL2(109)
MxL1-MxL2(112*109)
Lineage(seq #)
V704_0171_REN distances within and between lineages
Mean
Median
Distance
0.025
0.020
0.015
0.010
0.005
0.000
MxL1(60)
MxL2(2)
MxL1-MxL2(60*2)
Lineage(seq #)
V704_0423_REN distances within and between lineages
Mean
Median
Distance
0.025
0.020
0.015
0.010
0.005
0.000
MxL1(307)
MxL1-MxL2(307*1)
Lineage(seq #)
V704_0478_REN distances within and between lineages
Mean
Median
Distance
0.08
0.06
0.04
0.02
0.00
MxL1(94)
MxL2(17)
MxL3(4)
MxL4(122)
MxL1-MxL2(94*17)
MxL1-MxL3(94*4)
MxL1-MxL4(94*122)
MxL2-MxL3(17*4)
MxL2-MxL4(17*122)
MxL3-MxL4(4*122)
Lineage(seq #)

**S3.28 Fig.**

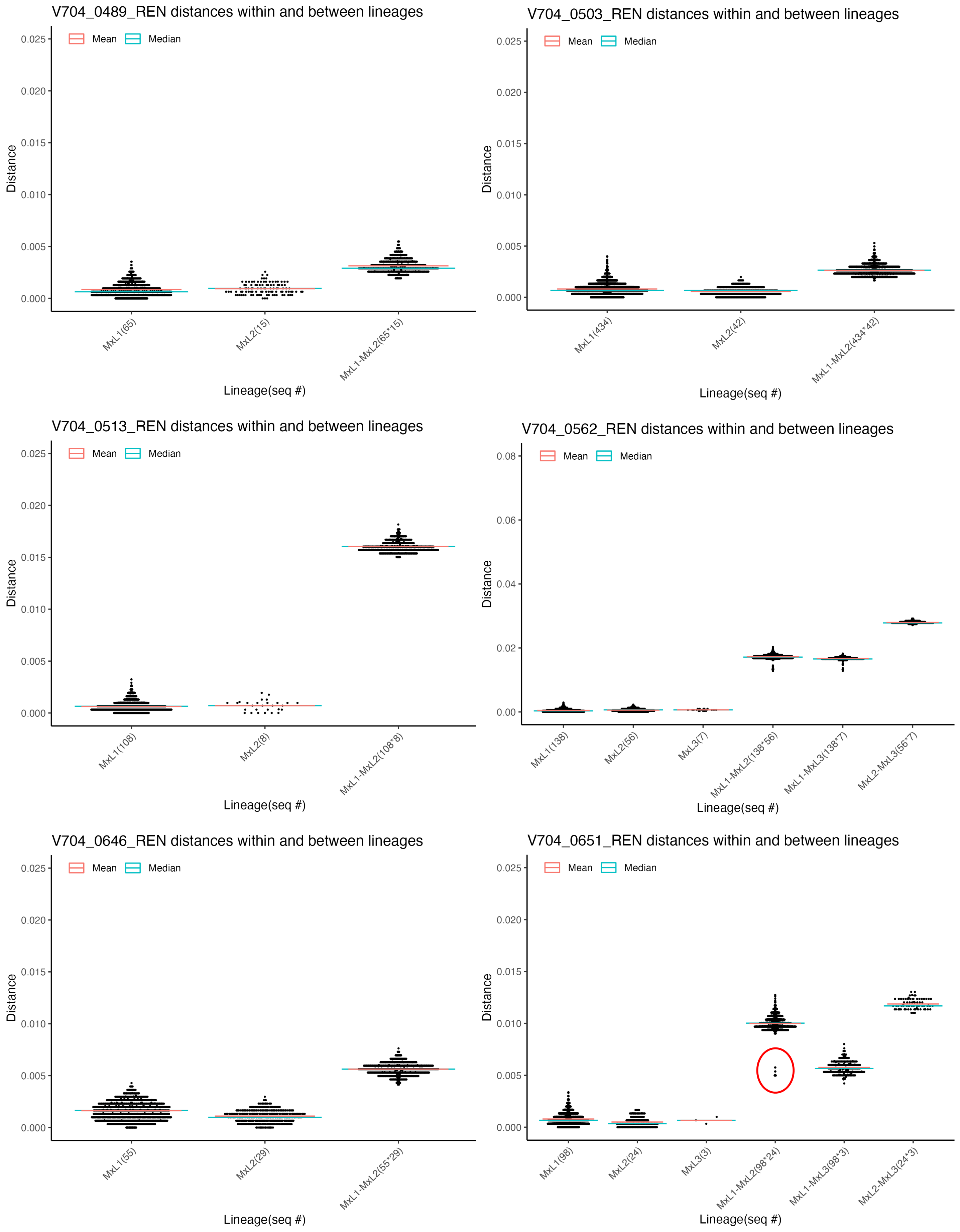
V704_0489_REN distances within and between lineages
Mean
Median
Distance
0.025
0.020
0.015
0.010
0.005
0.000
MxL1(65)
MxL2(15)
MxL1-MxL2(65*15)
Lineage(seq #)
V704_0503_REN distances within and between lineages
Mean
Median
Distance
0.025
0.020
0.015
0.010
0.005
0.000
MxL1(434)
MxL2(42)
MxL1-MxL2(434*42)
Lineage(seq #)
V704_0513_REN distances within and between lineages
Mean
Median
Distance
0.025
0.020
0.015
0.010
0.005
0.000
MxL1(108)
MxL2(8)
MxL1-MxL2(108*8)
Lineage(seq #)
V704_0562_REN distances within and between lineages
Mean
Median
Distance
0.08
0.06
0.04
0.02
0.00
MxL1(138)
MxL2(56)
MxL3(7)
MxL1-MxL2(138*56)
MxL1-MxL3(138*7)
MxL2-MxL3(56*7)
Lineage(seq #)
V704_0646_REN distances within and between lineages
Mean
Median
Distance
0.025
0.020
0.015
0.010
0.005
0.000
MxL1(55)
MxL2(29)
MxL1-MxL2(55*29)
Lineage(seq #)
V704_0651_REN distances within and between lineages
Mean
Median
Distance
0.025
0.020
0.015
0.010
0.005
0.000
MxL1(98)
MxL2(24)
MxL3(3)
MxL1-MxL2(98*24)
MxL1-MxL3(98*3)
MxL2-MxL3(24*3)
Lineage(seq #)

**S3.29 Fig.**

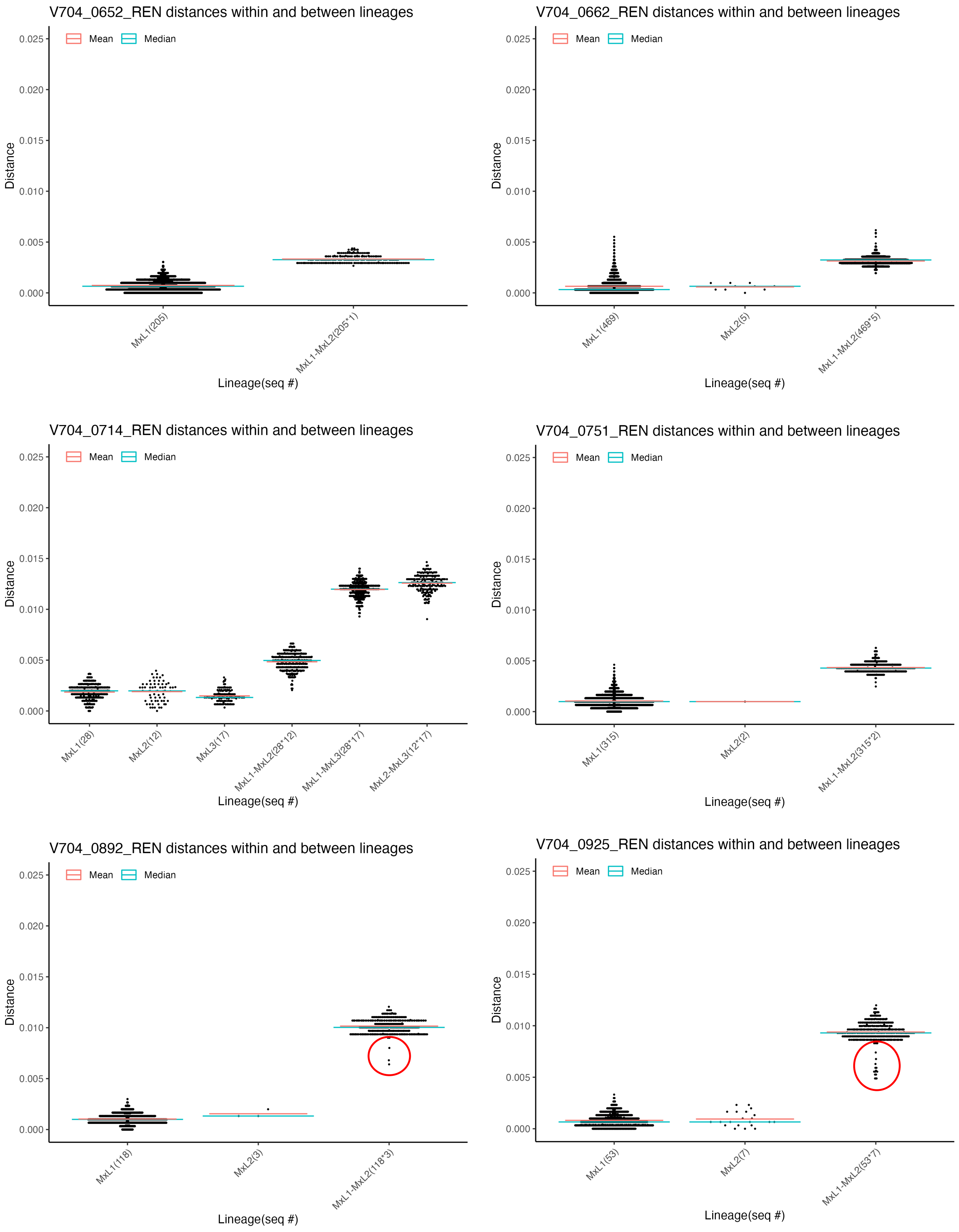

V704_0652_REN distances within and between lineages
Mean
Median
Distance
0.000
0.005
0.010
0.015
0.020
0.025
MxL1(205)
MxL1-MxL2(205*1)
Lineage(seq #)
V704_0662_REN distances within and between lineages
MxL1(469)
MxL2(5)
MxL1-MxL2(469*5)
V704_0714_REN distances within and between lineages
MxL1(28)
MxL2(12)
MxL3(17)
MxL1-MxL2(28*12)
MxL1-MxL3(28*17)
MxL2-MxL3(12*17)
V704_0751_REN distances within and between lineages
MxL1(315)
MxL2(2)
MxL1-MxL2(315*2)
V704_0892_REN distances within and between lineages
MxL1(118)
MxL2(3)
MxL1-MxL2(118*3)
V704_0925_REN distances within and between lineages
MxL1(53)
MxL2(7)
MxL1-MxL2(53*7)

**S3.30 Fig.**

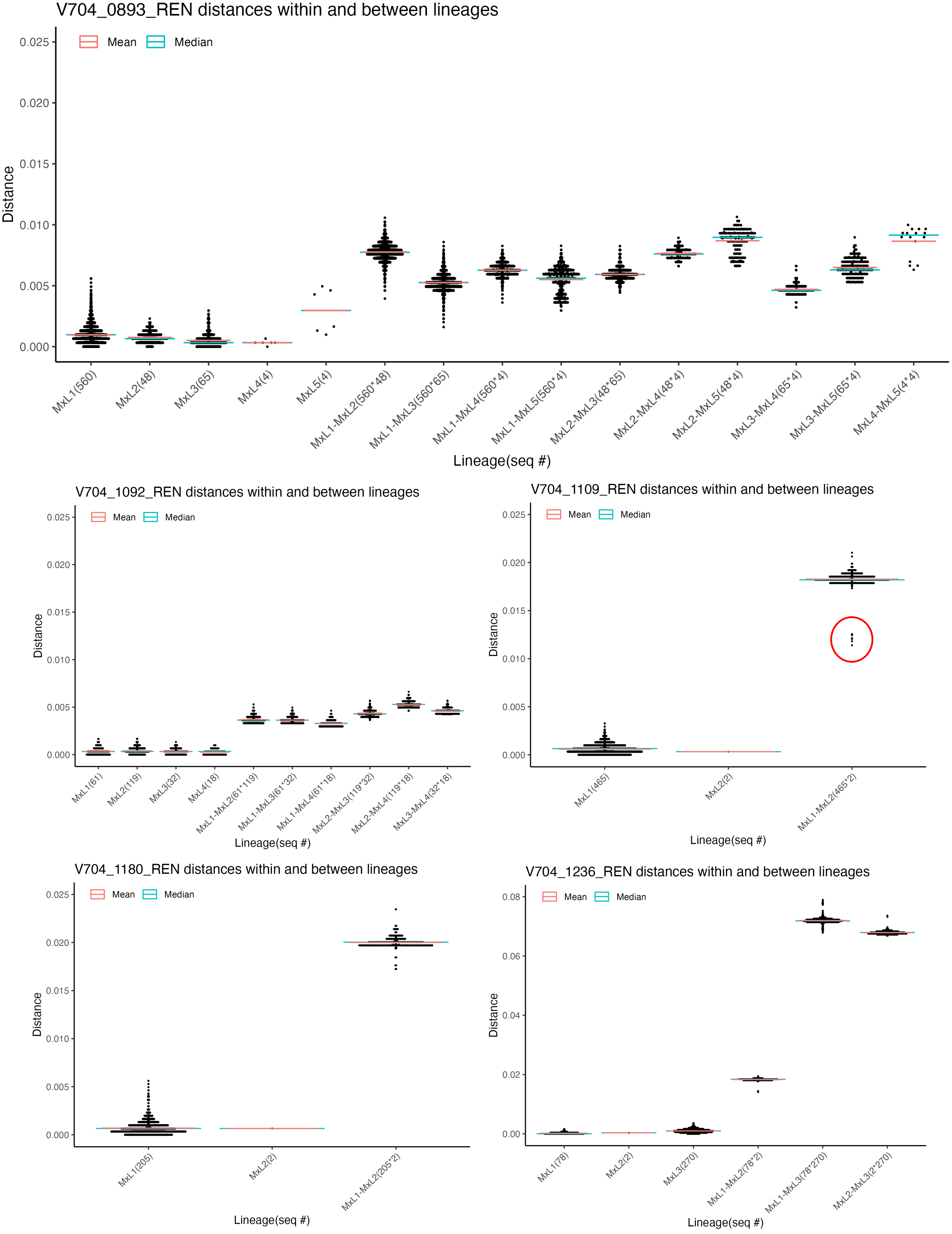
V704_0893_REN distances within and between lineages
Mean
Median
Distance
0.025
0.020
0.015
0.010
0.005
0.000
MxL1(560)
MxL2(48)
MxL3(65)
MxL4(4)
MxL5(4)
MxL1-MxL2(560*48)
MxL1-MxL3(560*65)
MxL1-MxL4(560*4)
MxL1-MxL5(560*4)
MxL2-MxL3(48*65)
MxL2-MxL4(48*4)
MxL2-MxL5(48*4)
MxL3-MxL4(65*4)
MxL3-MxL5(65*4)
MxL4-MxL5(4*4)
Lineage(seq #)
V704_1092_REN distances within and between lineages
Mean
Median
Distance
0.025
0.020
0.015
0.010
0.005
0.000
MxL1(61)
MxL2(119)
MxL3(32)
MxL4(18)
MxL1-MxL2(61*119)
MxL1-MxL3(61*32)
MxL1-MxL4(61*18)
MxL2-MxL3(119*32)
MxL2-MxL4(119*18)
MxL3-MxL4(32*18)
Lineage(seq #)
V704_1109_REN distances within and between lineages
Mean
Median
Distance
0.025
0.020
0.015
0.010
0.005
0.000
MxL1(465)
MxL2(2)
MxL1-MxL2(465*2)
Lineage(seq #)
V704_1180_REN distances within and between lineages
Mean
Median
Distance
0.025
0.020
0.015
0.010
0.005
0.000
MxL1(205)
MxL2(2)
MxL1-MxL2(205*2)
Lineage(seq #)
V704_1236_REN distances within and between lineages
Mean
Median
Distance
0.08
0.06
0.04
0.02
0.00
MxL1(78)
MxL2(2)
MxL3(270)
MxL1-MxL2(78*2)
MxL1-MxL3(78*270)
MxL2-MxL3(2*270)
Lineage(seq #)

**S3.31 Fig.**

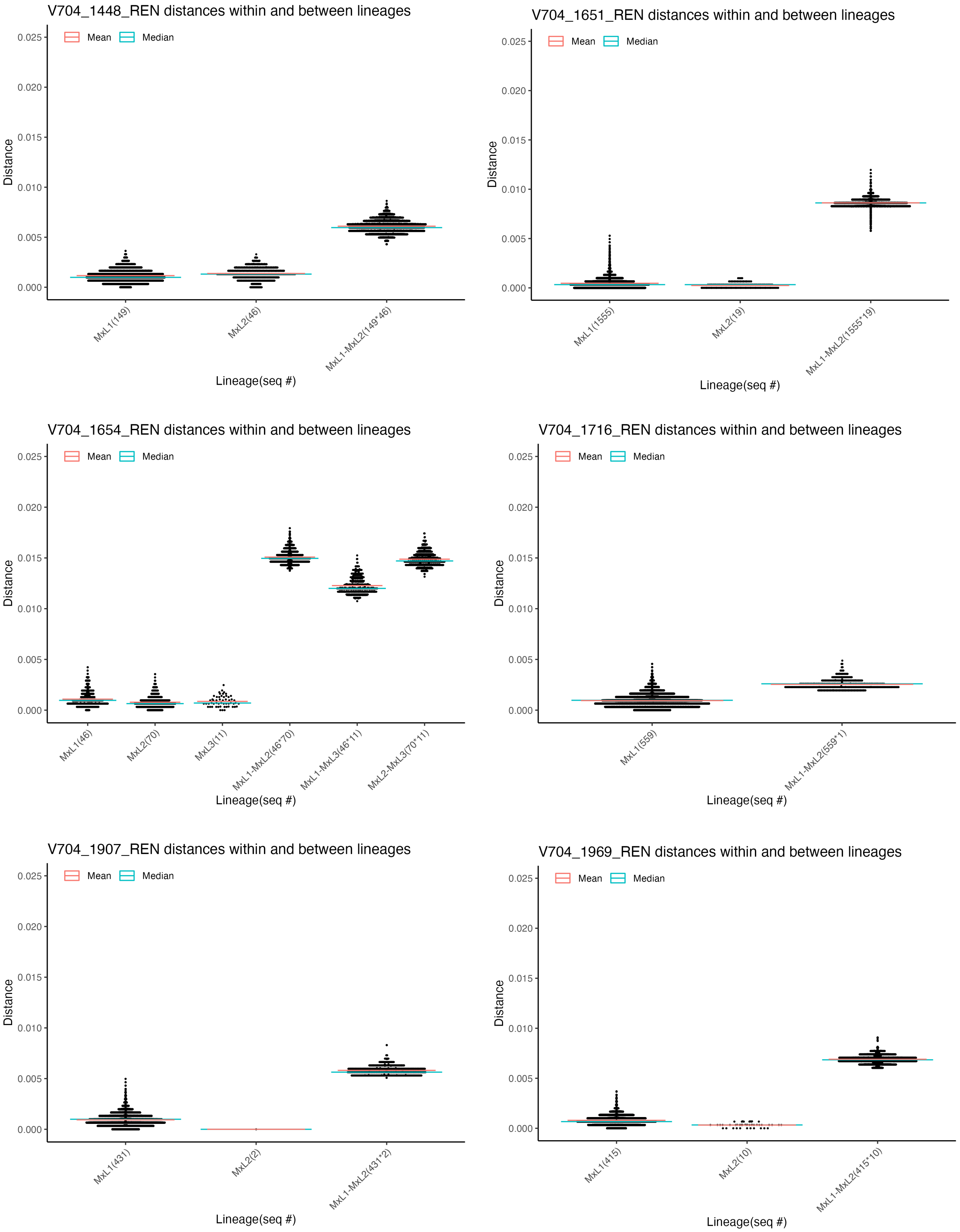

V704_1448_REN distances within and between lineages
Mean
Median
Distance
0.025
0.020
0.015
0.010
0.005
0.000
MxL1(149)
MxL2(46)
MxL1-MxL2(149*46)
Lineage(seq #)
V704_1651_REN distances within and between lineages
MxL1(1555)
MxL2(19)
MxL1-MxL2(1555*19)
V704_1654_REN distances within and between lineages
MxL1(46)
MxL2(70)
MxL3(11)
MxL1-MxL2(46*70)
MxL1-MxL3(46*11)
MxL2-MxL3(70*11)
V704_1716_REN distances within and between lineages
MxL1(559)
MxL1-MxL2(559*1)
V704_1907_REN distances within and between lineages
MxL1(431)
MxL2(2)
MxL1-MxL2(431*2)
V704_1969_REN distances within and between lineages
MxL1(415)
MxL2(10)
MxL1-MxL2(415*10)

**S3.32 Fig.**

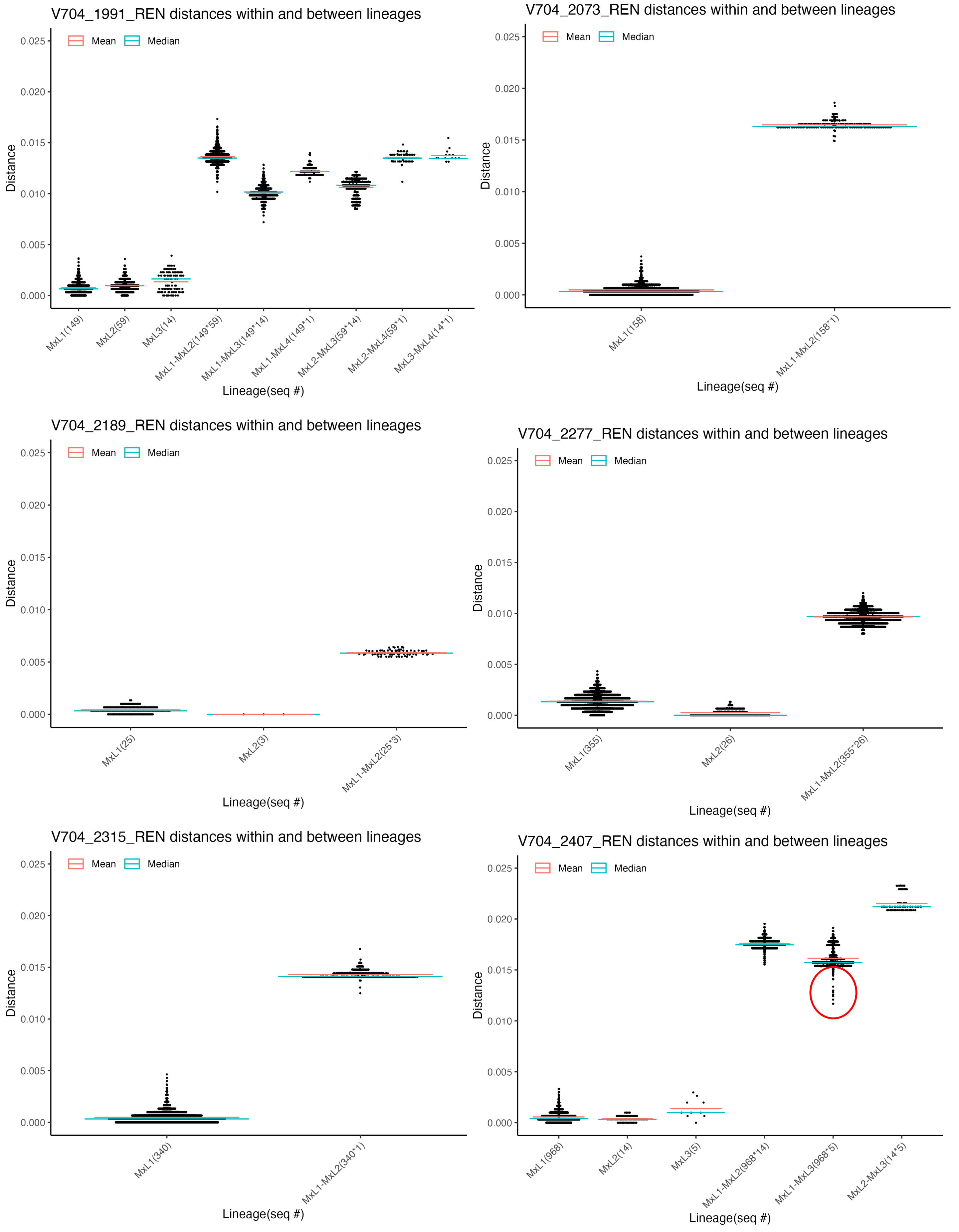
V704_1991_REN distances within and between lineages
Mean
Median
Distance
0.025
0.020
0.015
0.010
0.005
0.000
MxL1(149)
MxL2(59)
MxL3(14)
MxL1-MxL2(149*59)
MxL1-MxL3(149*14)
MxL1-MxL4(149*1)
MxL2-MxL3(59*14)
MxL2-MxL4(59*1)
MxL3-MxL4(14*1)
Lineage(seq #)
V704_2073_REN distances within and between lineages
MxL1(158)
MxL1-MxL2(158*1)
V704_2189_REN distances within and between lineages
MxL1(25)
MxL2(3)
MxL1-MxL2(25*3)
V704_2277_REN distances within and between lineages
MxL1(355)
MxL2(26)
MxL1-MxL2(355*26)
V704_2315_REN distances within and between lineages
MxL1(340)
MxL1-MxL2(340*1)
V704_2407_REN distances within and between lineages
MxL1(968)
MxL2(14)
MxL3(5)
MxL1-MxL2(968*14)
MxL1-MxL3(968*5)
MxL2-MxL3(14*5)

**S3.33 Fig.**

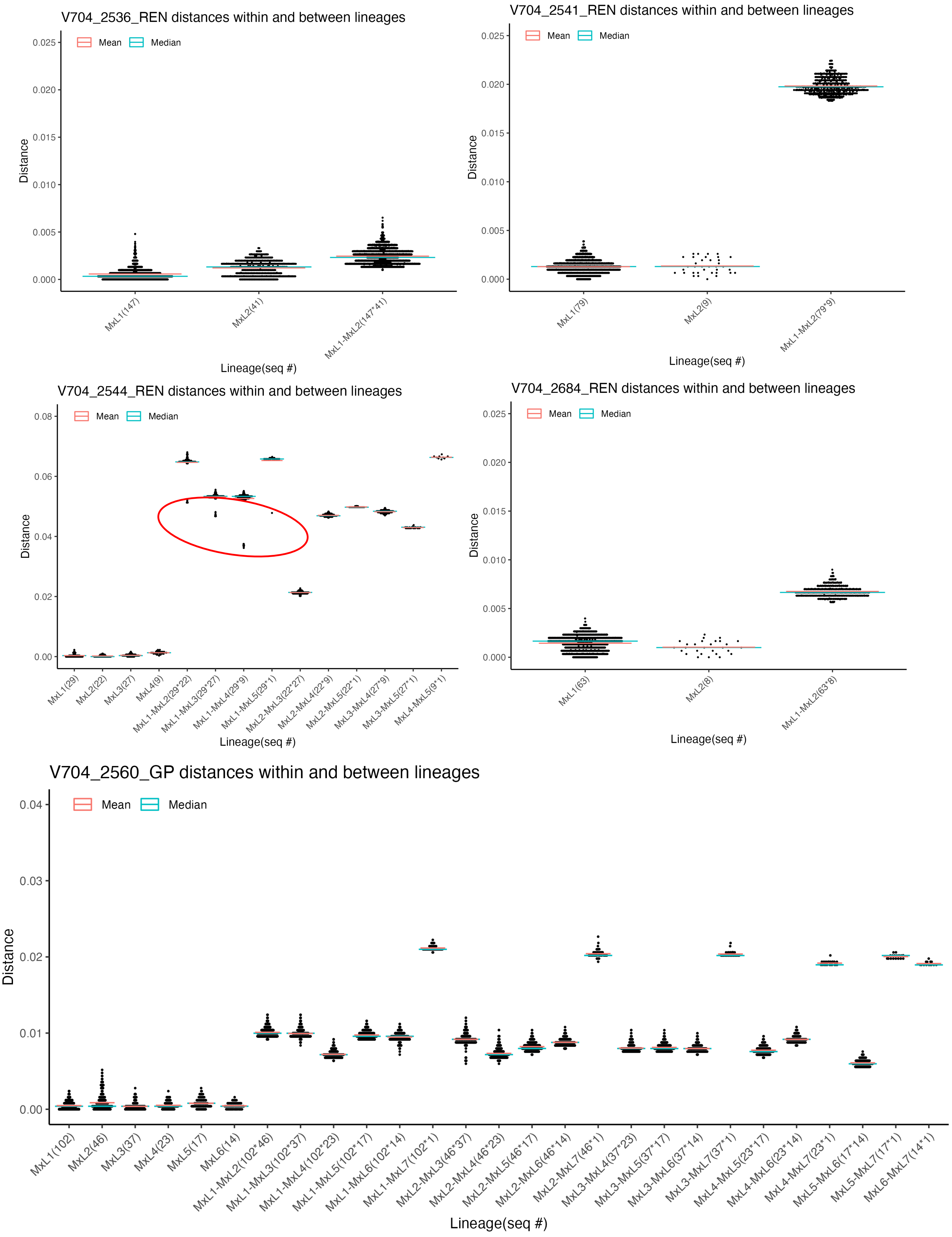
V704_2536_REN distances within and between lineages
Mean
Median
Distance
0.025
0.020
0.015
0.010
0.005
0.000
MxL1(147)
MxL2(41)
MxL1-MxL2(147*41)
Lineage(seq #)
V704_2541_REN distances within and between lineages
Mean
Median
Distance
0.025
0.020
0.015
0.010
0.005
0.000
MxL1(79)
MxL2(9)
MxL1-MxL2(79*9)
Lineage(seq #)
V704_2544_REN distances within and between lineages
Mean
Median
Distance
0.08
0.06
0.04
0.02
0.00
MxL1(29)
MxL2(22)
MxL3(27)
MxL4(9)
MxL1-MxL2(29*22)
MxL1-MxL3(29*27)
MxL1-MxL4(29*9)
MxL1-MxL5(29*1)
MxL2-MxL3(22*27)
MxL2-MxL4(22*9)
MxL2-MxL5(22*1)
MxL3-MxL4(27*9)
MxL3-MxL5(27*1)
MxL4-MxL5(9*1)
Lineage(seq #)
V704_2684_REN distances within and between lineages
Mean
Median
Distance
0.025
0.020
0.015
0.010
0.005
0.000
MxL1(63)
MxL2(8)
MxL1-MxL2(63*8)
Lineage(seq #)
V704_2560_GP distances within and between lineages
Mean
Median
Distance
0.04
0.03
0.02
0.01
0.00
MxL1(102)
MxL2(46)
MxL3(37)
MxL4(23)
MxL5(17)
MxL6(14)
MxL1-MxL2(102*46)
MxL1-MxL3(102*37)
MxL1-MxL4(102*23)
MxL1-MxL5(102*17)
MxL1-MxL6(102*14)
MxL1-MxL7(102*1)
MxL2-MxL3(46*37)
MxL2-MxL4(46*23)
MxL2-MxL5(46*17)
MxL2-MxL6(46*14)
MxL2-MxL7(46*1)
MxL3-MxL4(37*23)
MxL3-MxL5(37*17)
MxL3-MxL6(37*14)
MxL3-MxL7(37*1)
MxL4-MxL5(23*17)
MxL4-MxL6(23*14)
MxL4-MxL7(23*1)
MxL5-MxL6(17*14)
MxL5-MxL7(17*1)
MxL6-MxL7(14*1)
Lineage(seq #)

**S3.34 Fig.**

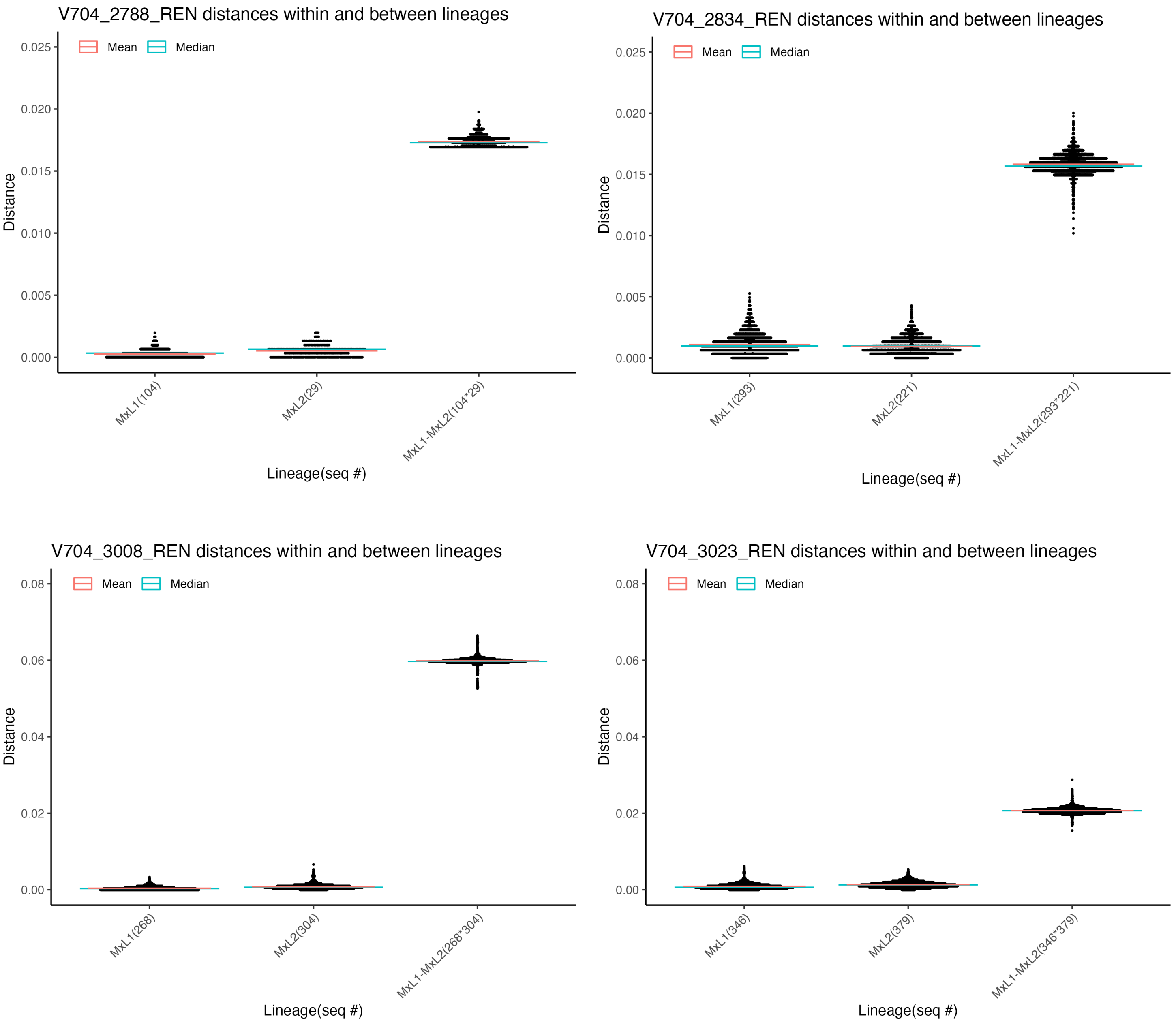
V704_2788_REN distances within and between lineages
Mean
Median
Distance
0.025
0.020
0.015
0.010
0.005
0.000
MxL1(104)
MxL2(29)
MxL1-MxL2(104*29)
Lineage(seq #)
V704_2834_REN distances within and between lineages
Mean
Median
Distance
0.025
0.020
0.015
0.010
0.005
0.000
MxL1(293)
MxL2(221)
MxL1-MxL2(293*221)
Lineage(seq #)
V704_3008_REN distances within and between lineages
Mean
Median
Distance
0.08
0.06
0.04
0.02
0.00
MxL1(268)
MxL2(304)
MxL1-MxL2(268*304)
Lineage(seq #)
V704_3023_REN distances within and between lineages
Mean
Median
Distance
0.08
0.06
0.04
0.02
0.00
MxL1(346)
MxL2(379)
MxL1-MxL2(346*379)
Lineage(seq #)

**S3.35 Fig.**

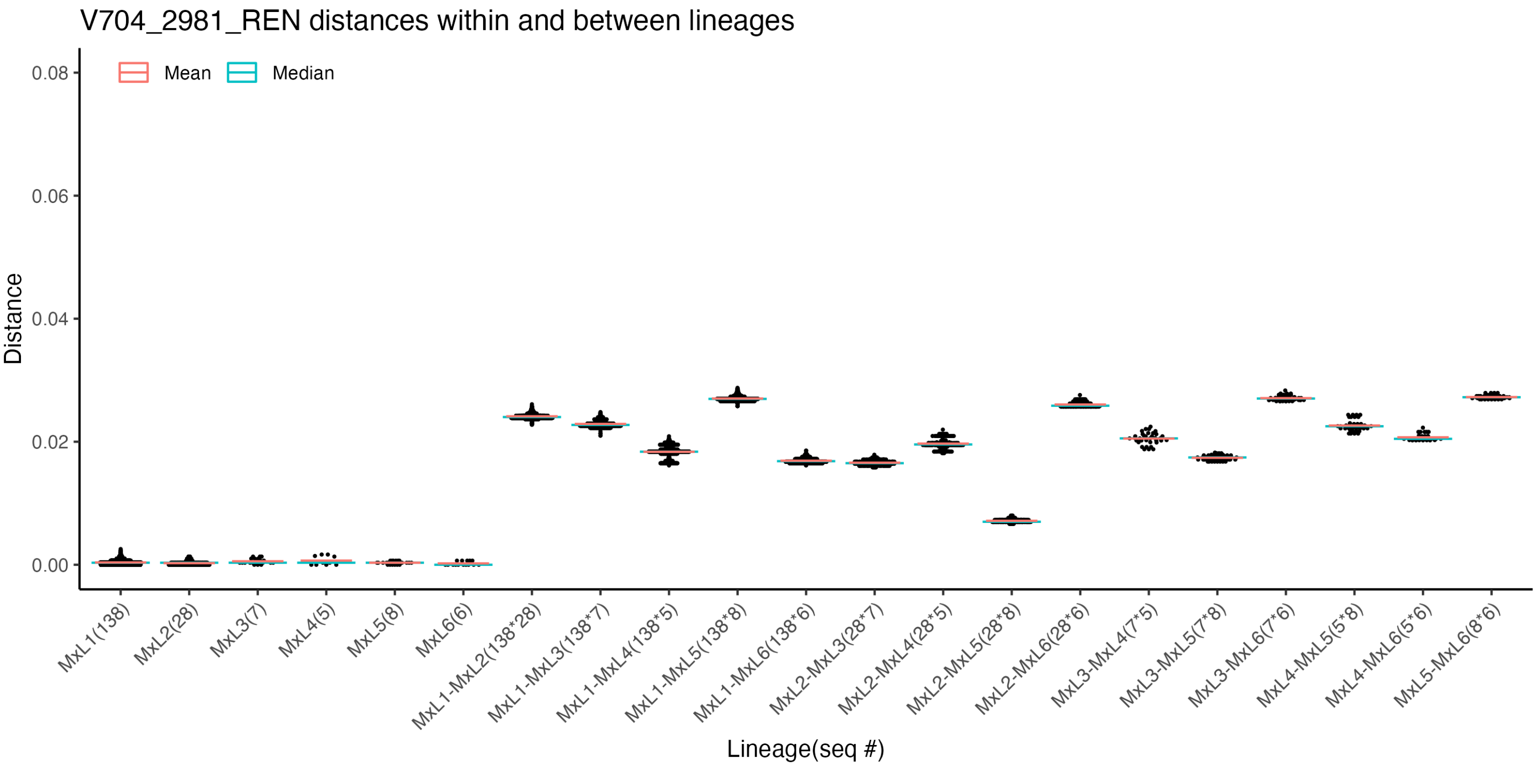
V704_2981_REN distances within and between lineages
Mean
Median
Distance
0.08
0.06
0.04
0.02
0.00
MxL1(138)
MxL2(28)
MxL3(7)
MxL4(5)
MxL5(8)
MxL6(6)
MxL1-MxL2(138*28)
MxL1-MxL3(138*7)
MxL1-MxL4(138*5)
MxL1-MxL5(138*8)
MxL1-MxL6(138*6)
MxL2-MxL3(28*7)
MxL2-MxL4(28*5)
MxL2-MxL5(28*8)
MxL2-MxL6(28*6)
MxL3-MxL4(7*5)
MxL3-MxL5(7*8)
MxL3-MxL6(7*6)
MxL4-MxL5(5*8)
MxL4-MxL6(5*6)
MxL5-MxL6(8*6)
Lineage(seq #)

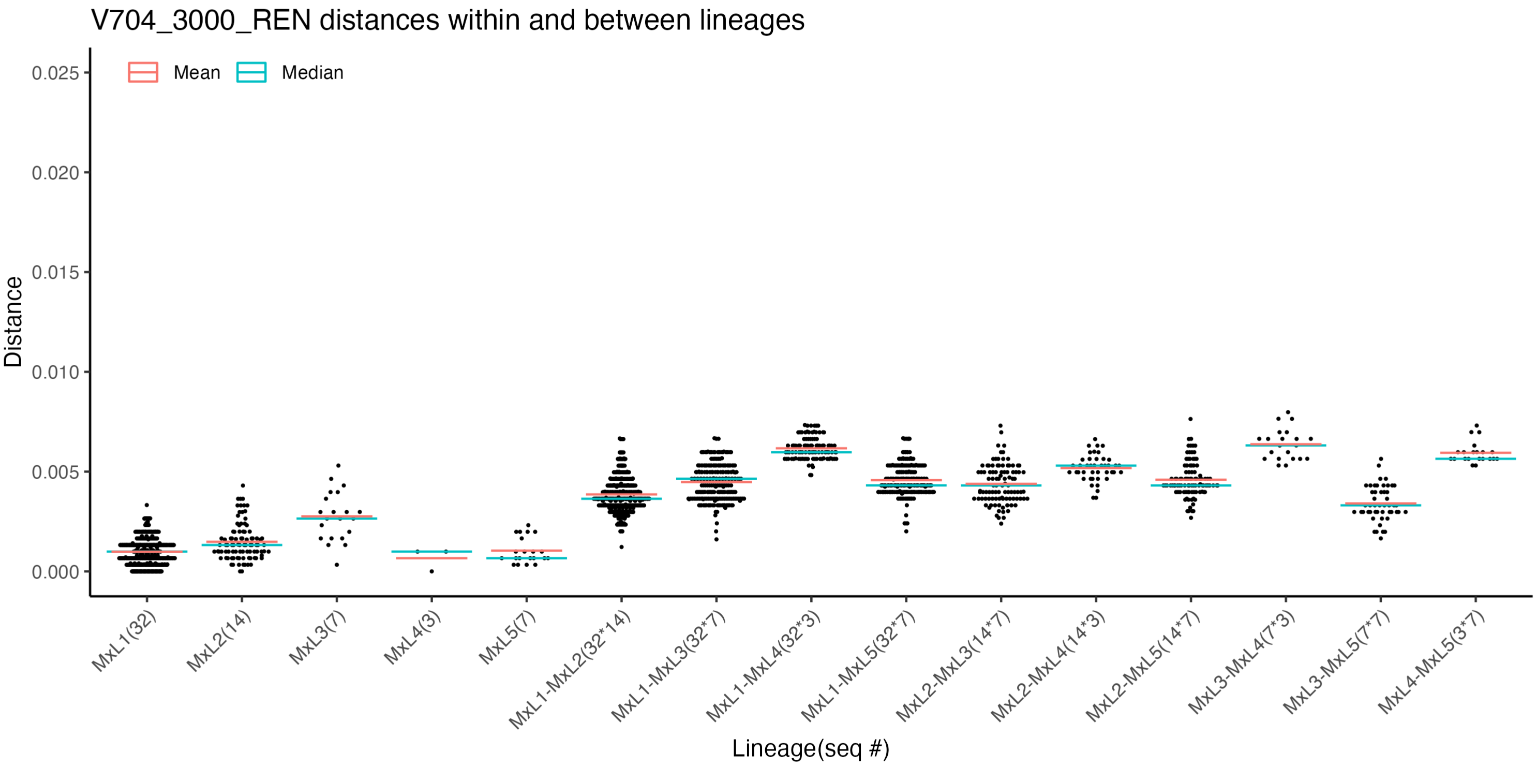
V704_3000_REN distances within and between lineages
Mean
Median
Distance
0.025
0.020
0.015
0.010
0.005
0.000
MxL1(32)
MxL2(14)
MxL3(7)
MxL4(3)
MxL5(7)
MxL1-MxL2(32*14)
MxL1-MxL3(32*7)
MxL1-MxL4(32*3)
MxL1-MxL5(32*7)
MxL2-MxL3(14*7)
MxL2-MxL4(14*3)
MxL2-MxL5(14*7)
MxL3-MxL4(7*3)
MxL3-MxL5(7*7)
MxL4-MxL5(3*7)
Lineage(seq #)

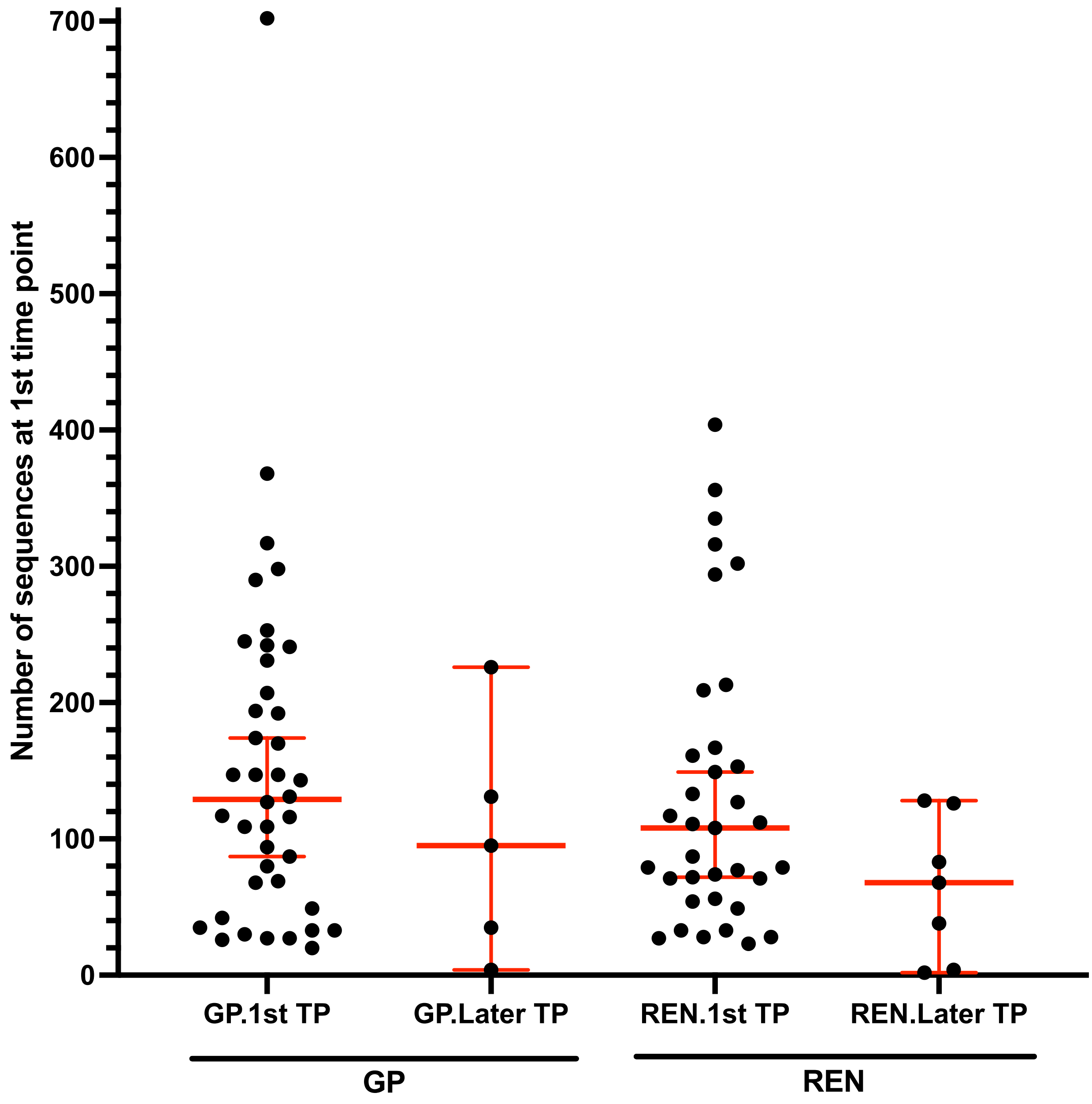

**Supplementary Fig. 7. Number of sequence obtained at the first time point as a function of when multiple lineages were detected.** "XX.1$^{st}$ TP" refers to detection of multiple lineages at the 1$^{st}$ time point examined. "XX.Later TP" refers to detection of multiple lineages only at later time point(s) examined.

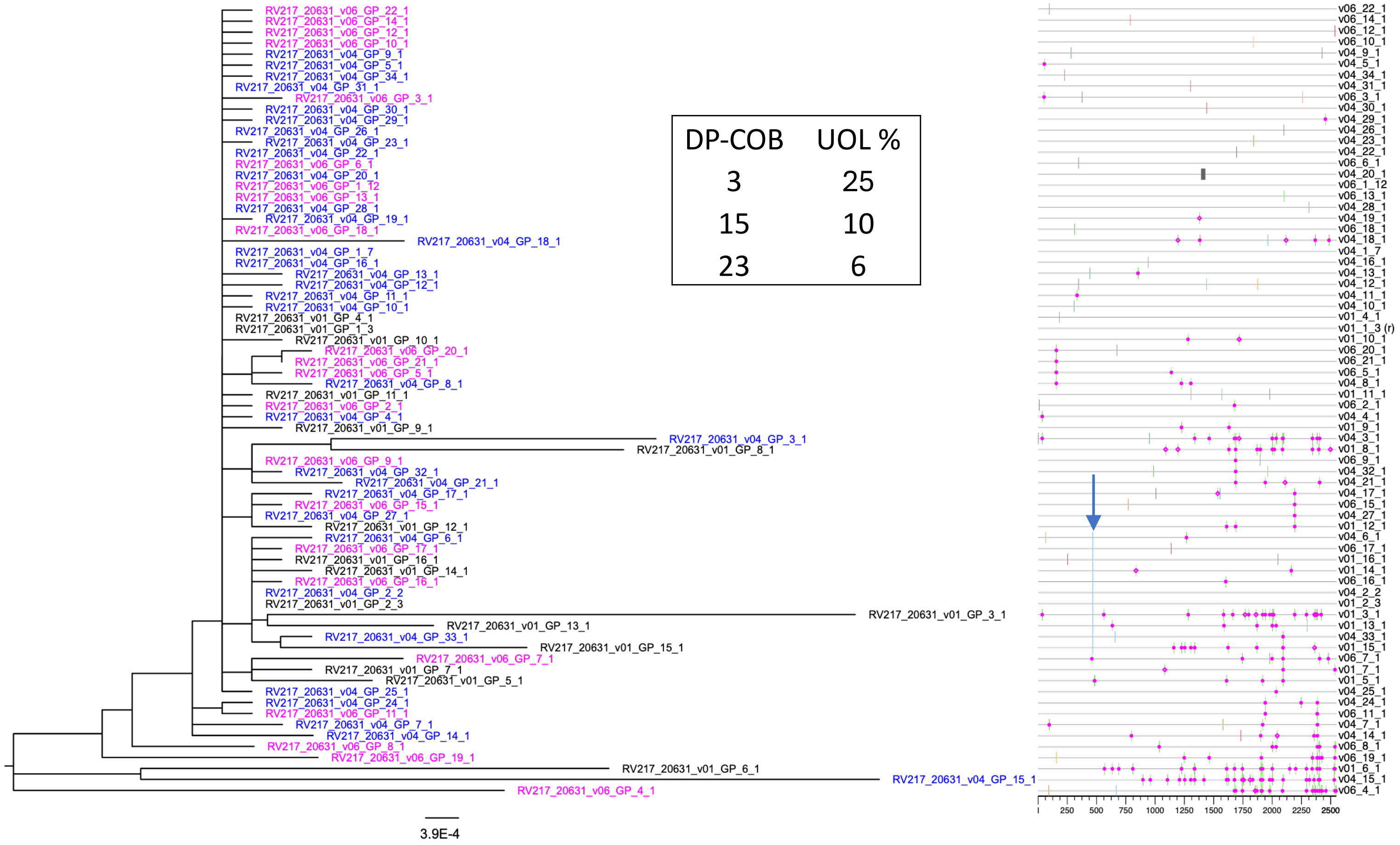

**Supplementaty Fig. 8. Uncertain-Origin-Lineage (UOL).** Phylobook output showing a phylogenetic tree and highlighter plot from a RV217 cohort participant. UOL sequences are shown as a blue stripe in the highlighter plot (blue arrow). The days post COB and the relative representation of the UOL at each time point are shown in the inset.

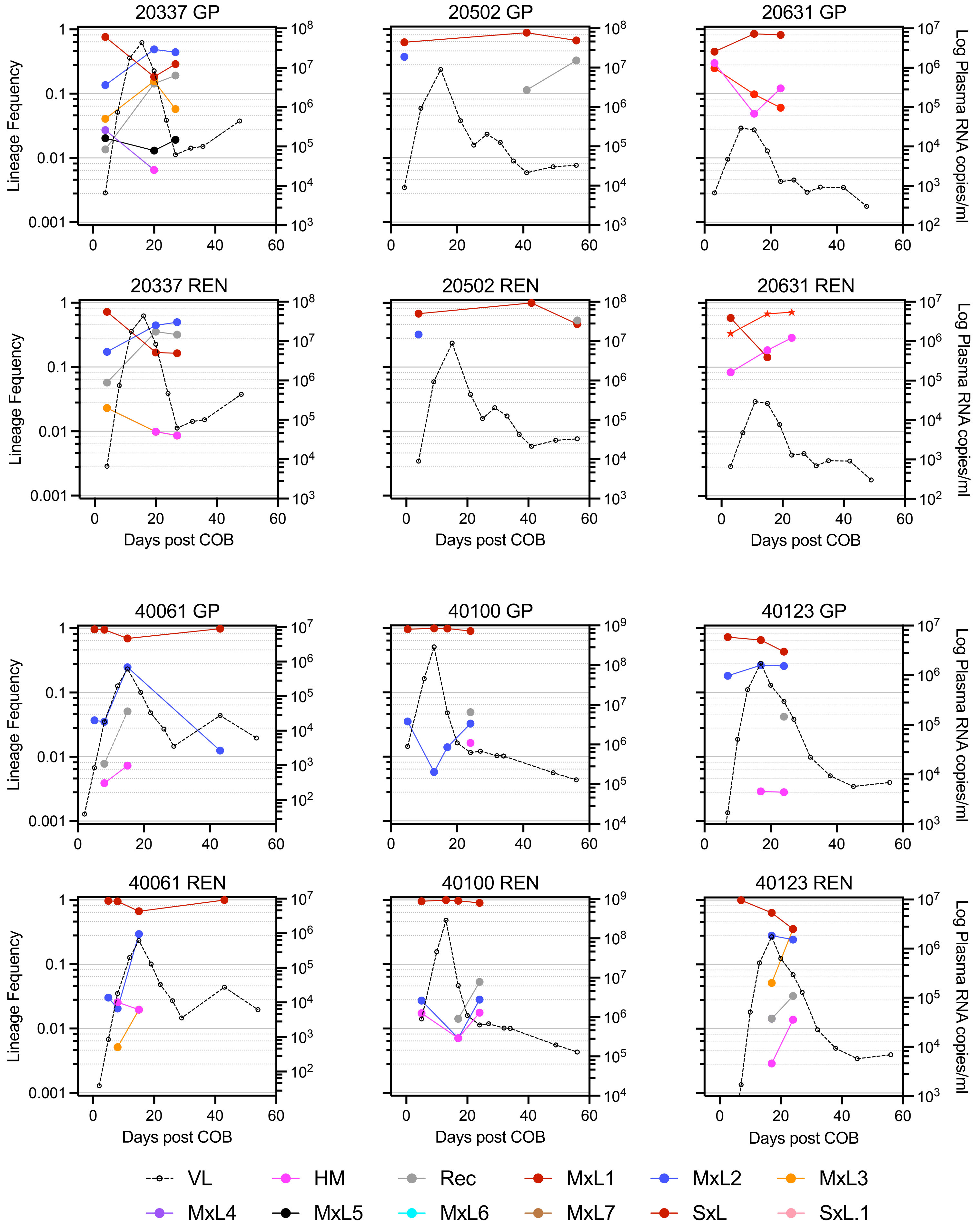

**Supplementary Fig. 9.1. Plasma HIV RNA viral load and lineage frequencies over time.** Data from participants from the RV217 cohort and with multiple transmitted lineages are shown. See legend to Fig 2 for key.

**Supplementary Fig. 9.2.**

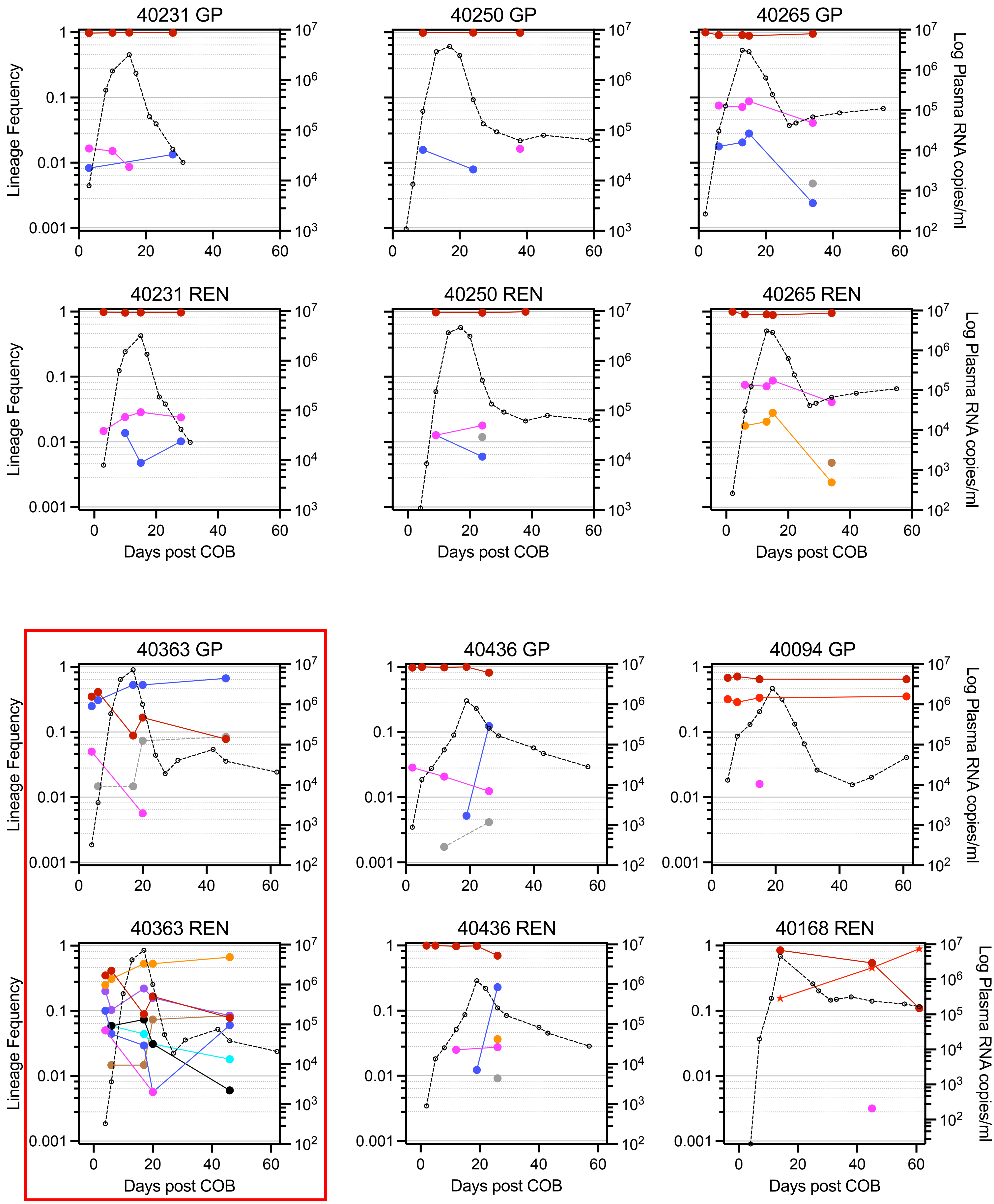

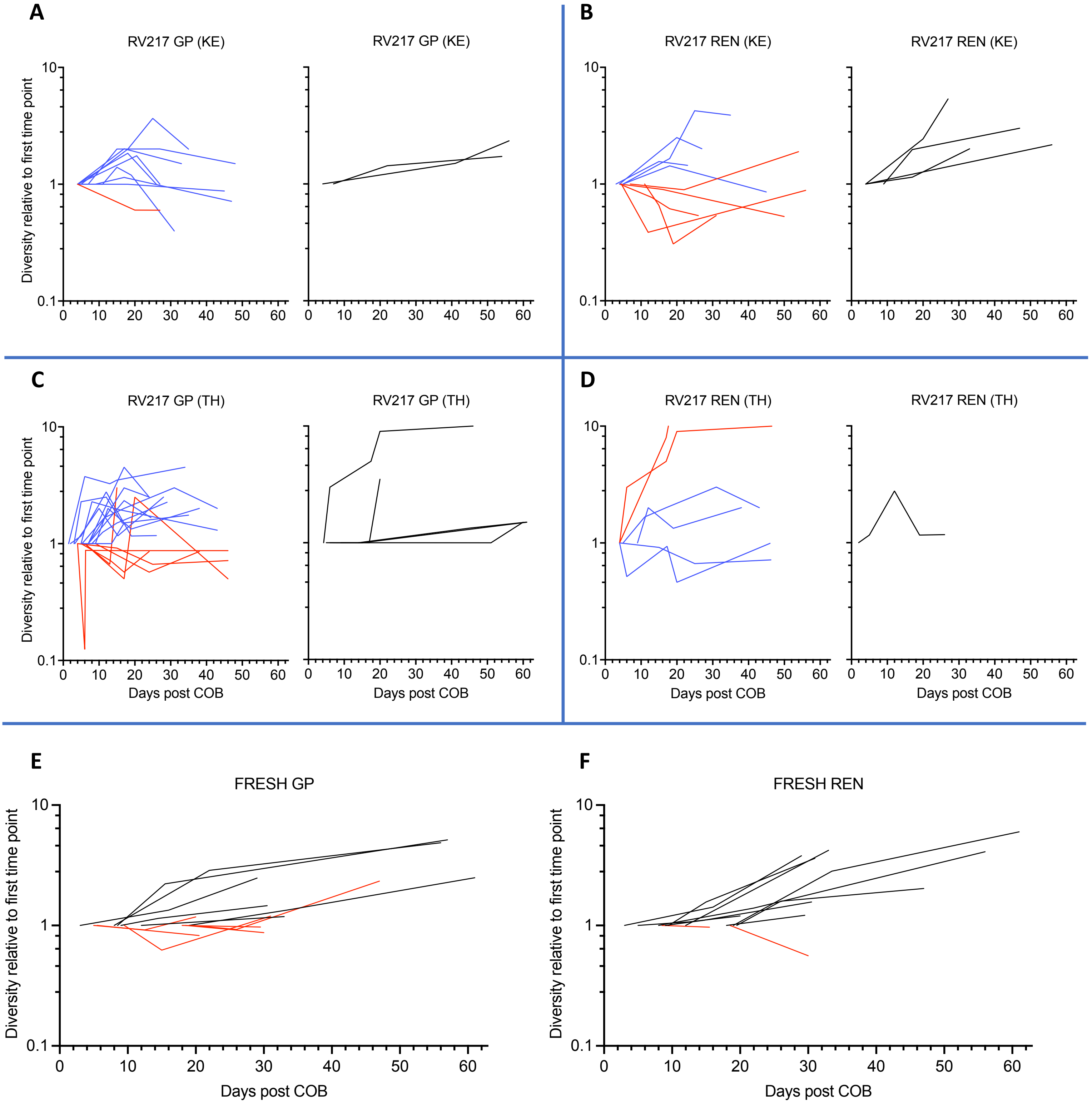

**Supplementary Fig. 10 Viral population diversification.** Maximum likelihood pairwise distances within all lineages from RV217 (**A-D**) and FRESH (**E-F**) cohorts are plotted as a function of days post COB for the GP (**A, C, E**) and REN (**B, D, F**) regions. All diversity measurements are plotted in relation to the diversity at the first time point (i.e., "1" on the y-axis). Red lines indicate instances in which the diversity decreased from the 1st to the 2nd time point, blue lines indicate instance in which the diversity decreased at any later time point, and black lines (plotted separately in A-D) indicate instances in which no drop in diversity was detected.

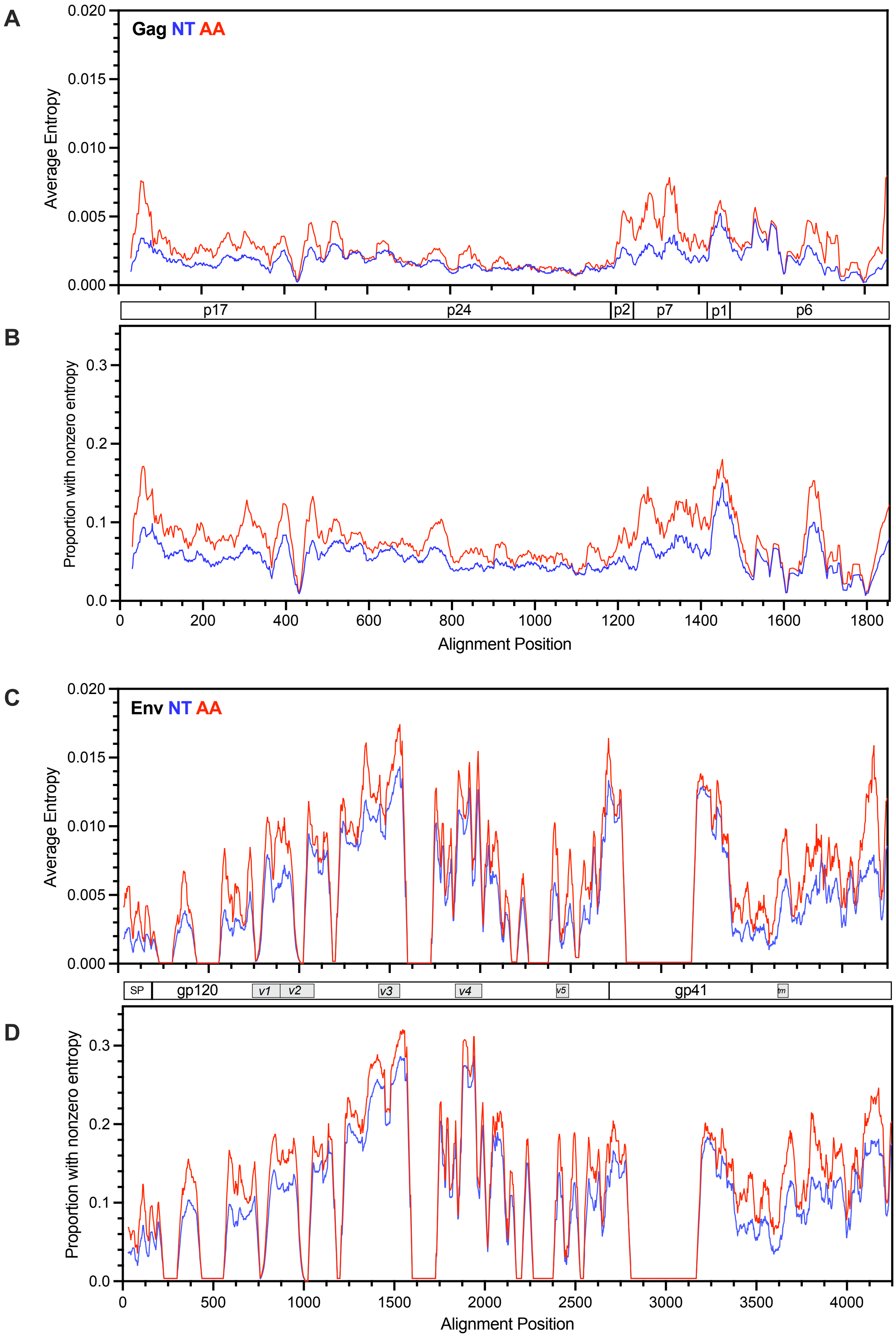

**Supplementary Fig. 11. Entropy in the Gag and Env genes.** Nucleotide (NT, blue lines) and amino acid (AA, red lines) alignments of sequences from all lineage were compared, in 30 NT or 10 AA sliding windows. Panels A and C show Shannon entropies, whereas panels B and D show the proportion of lineages that had nonzero entropy. Data was plotted using a full alignment of all lineages having at least 10 members derived from all participants.

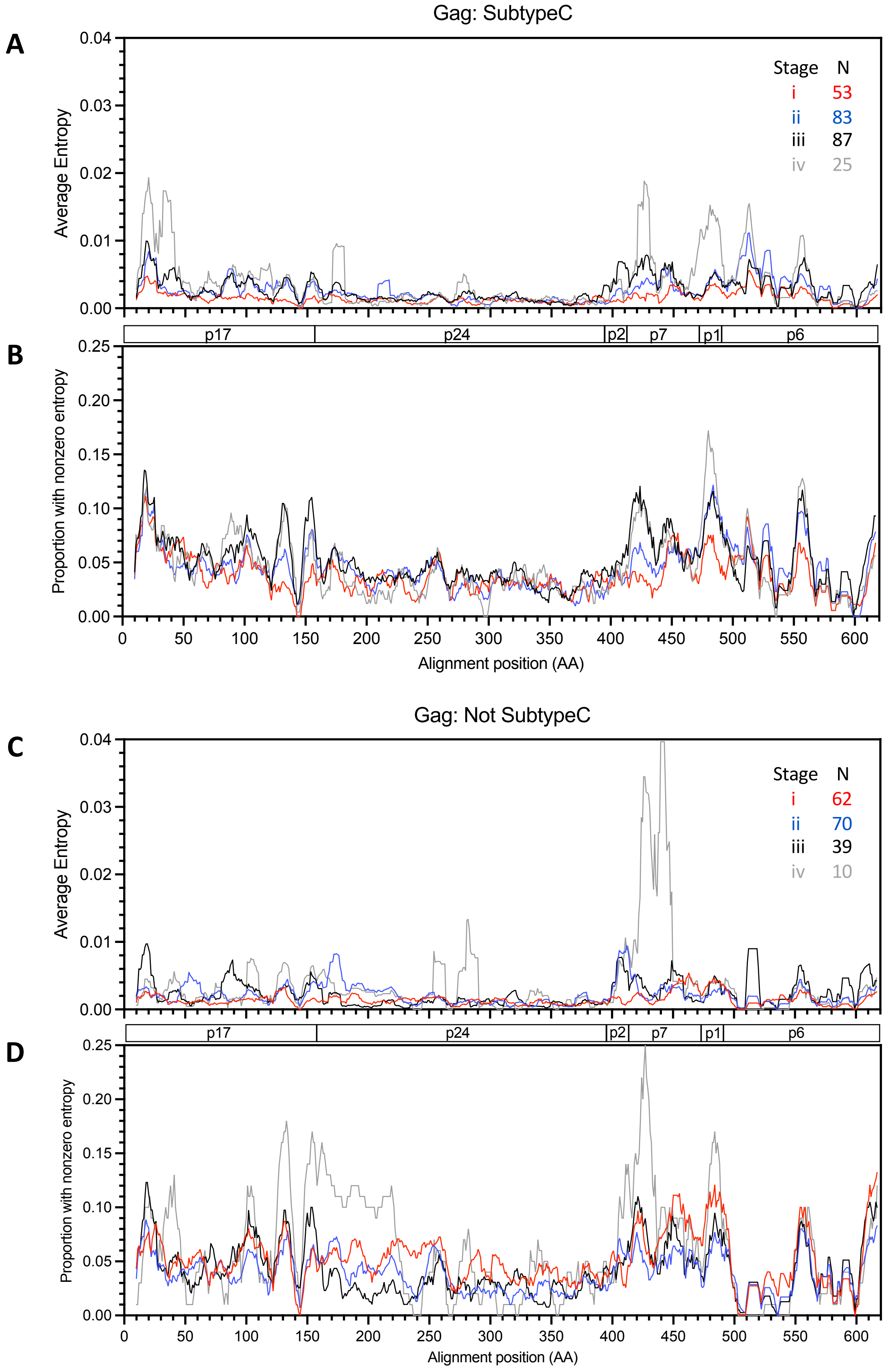

**Supplementary Fig. 12.** Entropy of Gag proteins during the i-iv stages of early infection. To limit variation due to small sample sizes, lineage-based datasets were split into Subtype C (**A, B**) and not Subtype C (**C, D**). Panels A and C show Shannon entropy values and B and D show the proportion of lineages with any entropy at a given amino acid position. Data is plotted in sliding windows of 10 amino acids using a full alignment of all lineages having at least 10 members derived from all participants. The keys in panels A and C show the color coding and the number of lineage datasets used to plot each stage pattern.

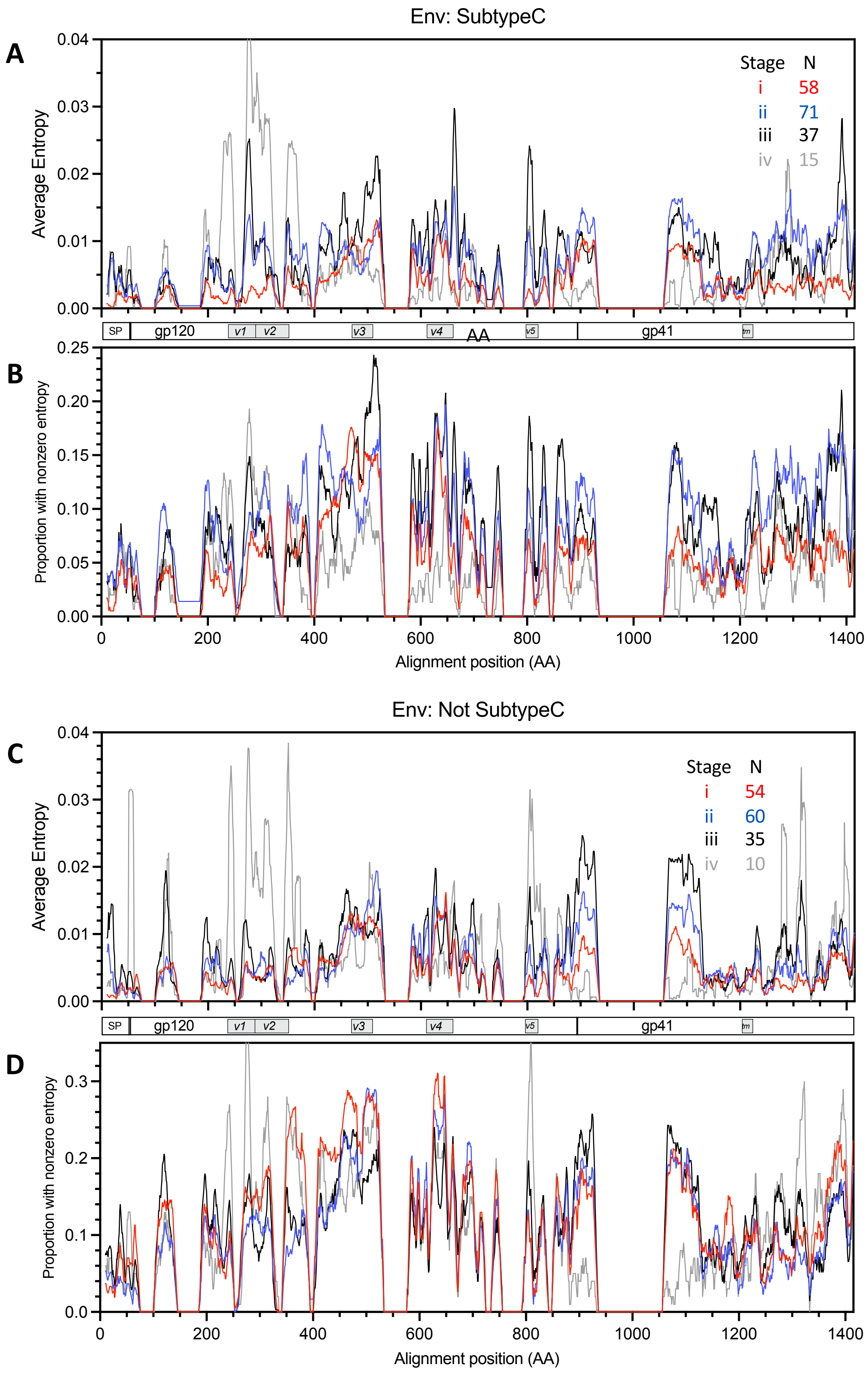

**Supplementary Figure 13.** Entropy of Env proteins during the i-iv stages of early infection. To limit variation due to small sample sizes, lineage-based datasets were split into Subtype C (**A, B**) and not Subtype C (**C, D**). Panels A and C show Shannon entropy values and B and D show the proportion of lineages with any entropy at a given amino acid position. Data is plotted in sliding windows of 10 amino acids using a full alignment of all lineages having at least 10 members derived from all participants. The keys in panels A and C show the color coding and the number of lineage datasets used to plot each stage pattern.

**Supplementary Fig. 14. Number of potential N-linked glycosylation site by stage and subtype groupings.** The red lines in panel **A** indicate medians and 95% CI.

**A**

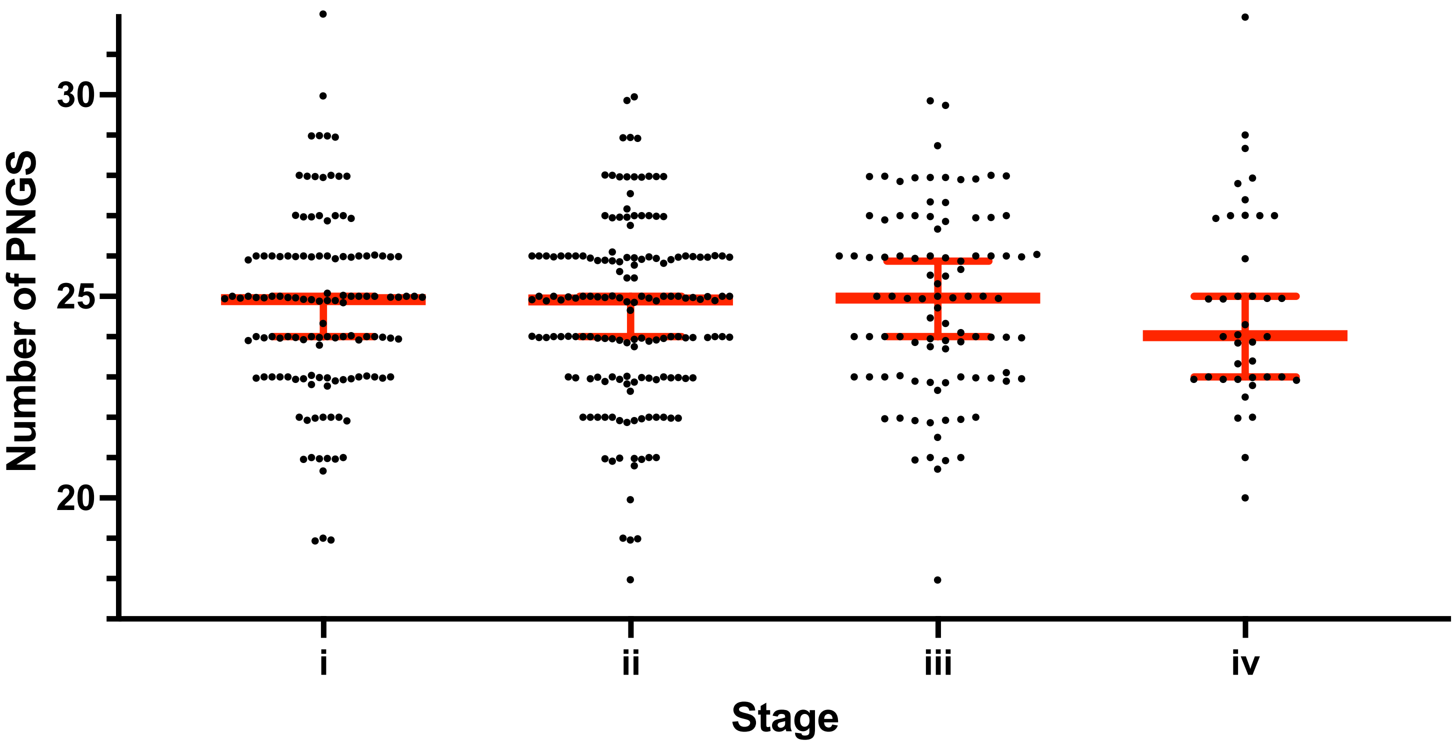

**B**

| Numbers of PNGS by stage of acute infection and Subtype | | | | |
|---|---|---|---|---|
| **Stage** | **i** | **ii** | **iii** | **iv** |
| **All cohorts** | | | | |
| Mean | 24.6 | 24.6 | 24.8 | 24.7 |
| Median | 24.9 | 24.9 | 25.0 | 24.0 |
| **Subtype C** | | | | |
| Mean | 24.2 | 24.2 | 24.3 | 24.6 |
| Median | 24.0 | 24.0 | 24.0 | 24.2 |
| **NonC** | | | | |
| Mean | 25.1 | 25.0 | 25.5 | 24.9 |
| Median | 25.0 | 25.0 | 25.3 | 24.0 |

**Supplementary Fig. 15. Selected sites within and immediately surrounding PNGS.** The number of AA sites within and nearby NLGS sequons that experienced positive and negative selection above a 0.9 probability threshold are shown. Positively (**A**) and negatively (**B**) selected sites that impact sequons and adjacent sequences are indicated. Selected sites N-terminal (negative numbers on the x-axis) and C-terminal (positive numbers) to the sequons are shown with blue bars. Loss (panel A) or preservation (panel B) of PNGS are shown in shades of red with the positions within the NX(T/S) sequon. The cyan bar in the N column of panel A indicates a single gain of a PNGS, and the black bars in the X positions indicate the selection of a Proline at this position, which greatly diminishes the likelihood of glycosylation.

**A**

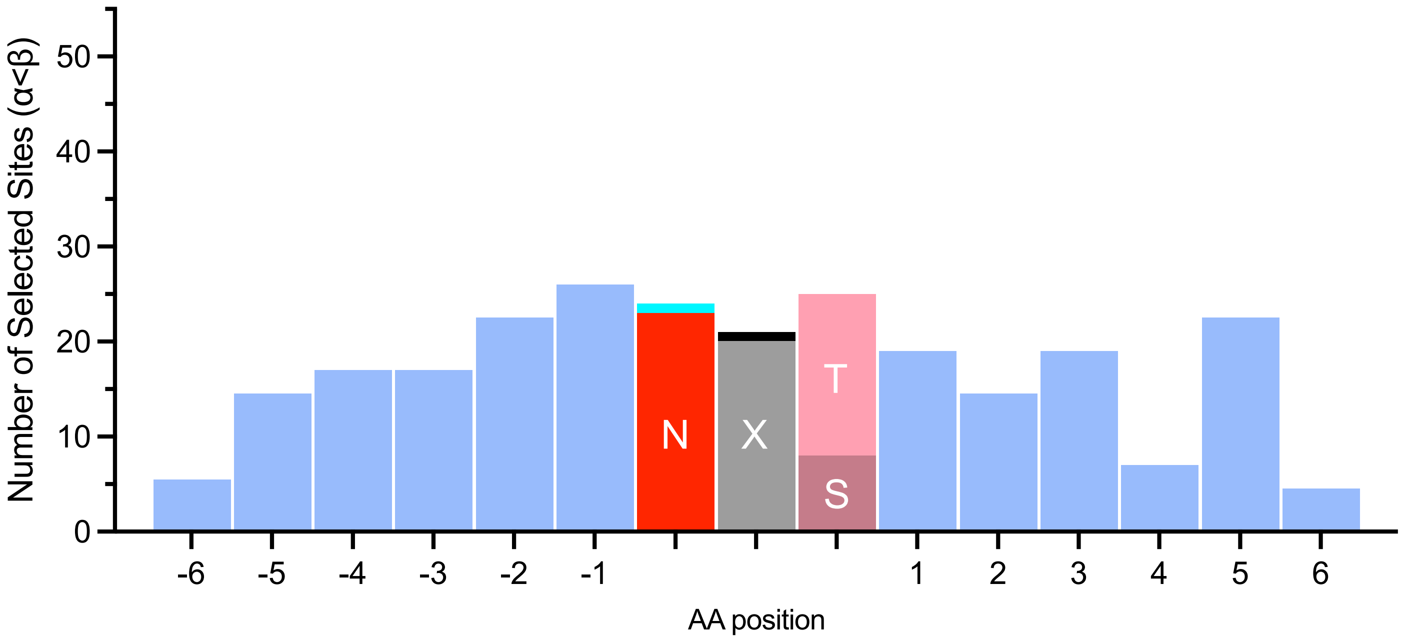

**B**

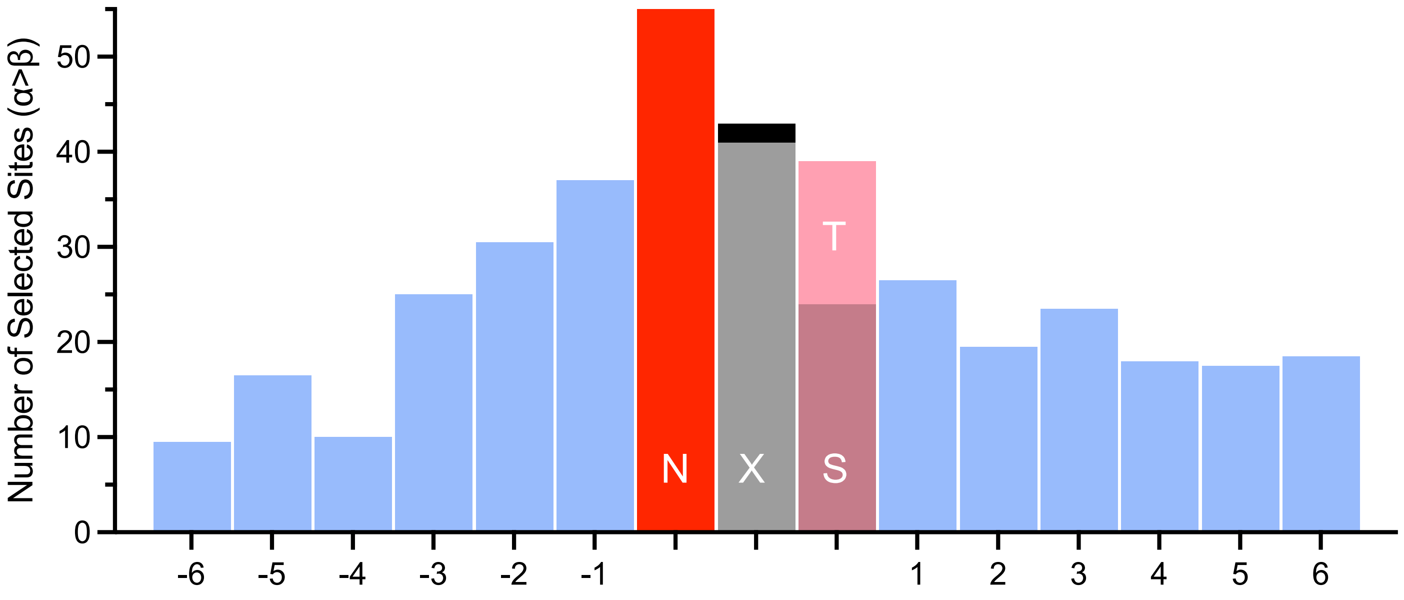

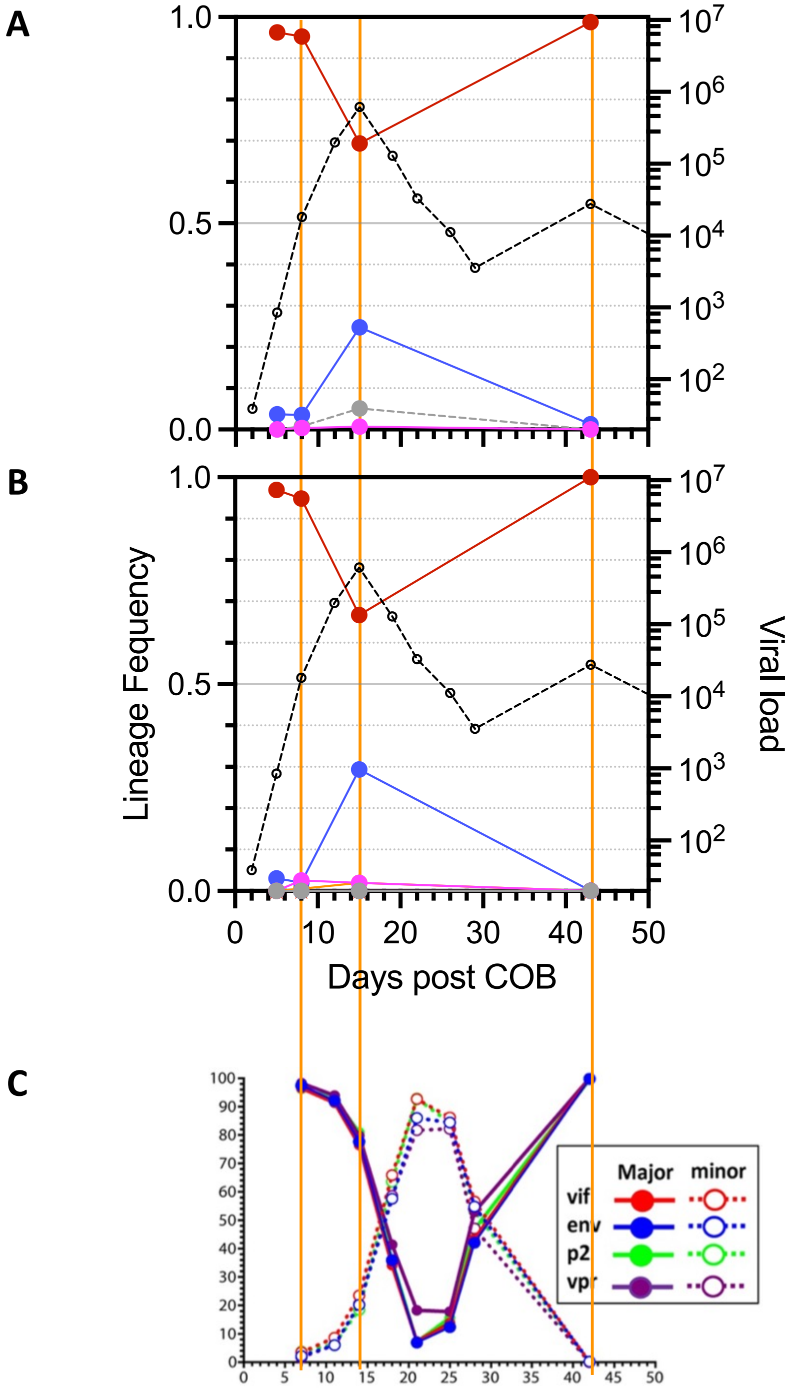

**Supplementary Fig. 15. Comparison of lineage frequencies over time from the current and a previous study.** Data from participant 40061 from the RV217 cohort and with multiple transmitted lineages is shown. Lineage frequencies in GP (**A**) and REN (**B**) from this study as well as major and minor lineages detected in 4 viral genome regions from a previous study by Kijak et al (PLoS pathogens. 2017;13(7):e1006510) (**C**). The major and minor lineages described by Kijak et al correspond to the 1st (red symbols) and 2nd (blue) most prevalent lineages from the current study. As shown by the intervening time points sequenced by Kijak et al, the minor lineage was transiently the most abundant in the interval between 14-42 days COB.
